# Design and synthesis of mirror image fluorescent nucleotide reversible terminator, (L)-3ʹ-O-azidomethyl-dGTP-N_3_-fluorophore and (D)-9°N mutant DNA polymerase for mirror image (L)-DNA sequencing by synthesis

**DOI:** 10.64898/2025.12.03.692211

**Authors:** Dae H. Kim, Bo-Shun Huang, Eric J. Yik, Jacob R. Baker, Rajasekaran Ganesan, Paramesh Jangili, Guangyan Zhou, A Michael Sismour, Esau L. Medina, Jennie S. Lee

## Abstract

Functional DNA molecules such as aptamers, catalytic DNAzymes, and dynamic nanostructures have many promising applications in diagnostics and therapeutics yet suffer from limited utility due to immunogenicity and unwanted interactions with other biomolecules, leading to nuclease degradation and unintended sequence driven binding to native sample DNA. Bio-orthogonal, (L)-form or mirror image DNA overcome these challenges yet suffers from the inability to sequence these molecules in a high-throughput manner, a necessary step in the rapid creation of functional (L)-DNA. Here, we report for the first time, design, synthesis, and functional demonstration of (L)-DNA sequencing by synthesis (SBS) using a fluorescent nucleotide reversible terminator, (L)-3ʹ-*O*-azidomethyl-dGTP-N_3_-fluorophore paired with a (D)-9°N mutant DNA polymerase, both enantiomers (i.e., mirror image) of their natural biologically active forms.

**TABLE OF CONTENTS (GRAPHICAL ABSTRACT):** 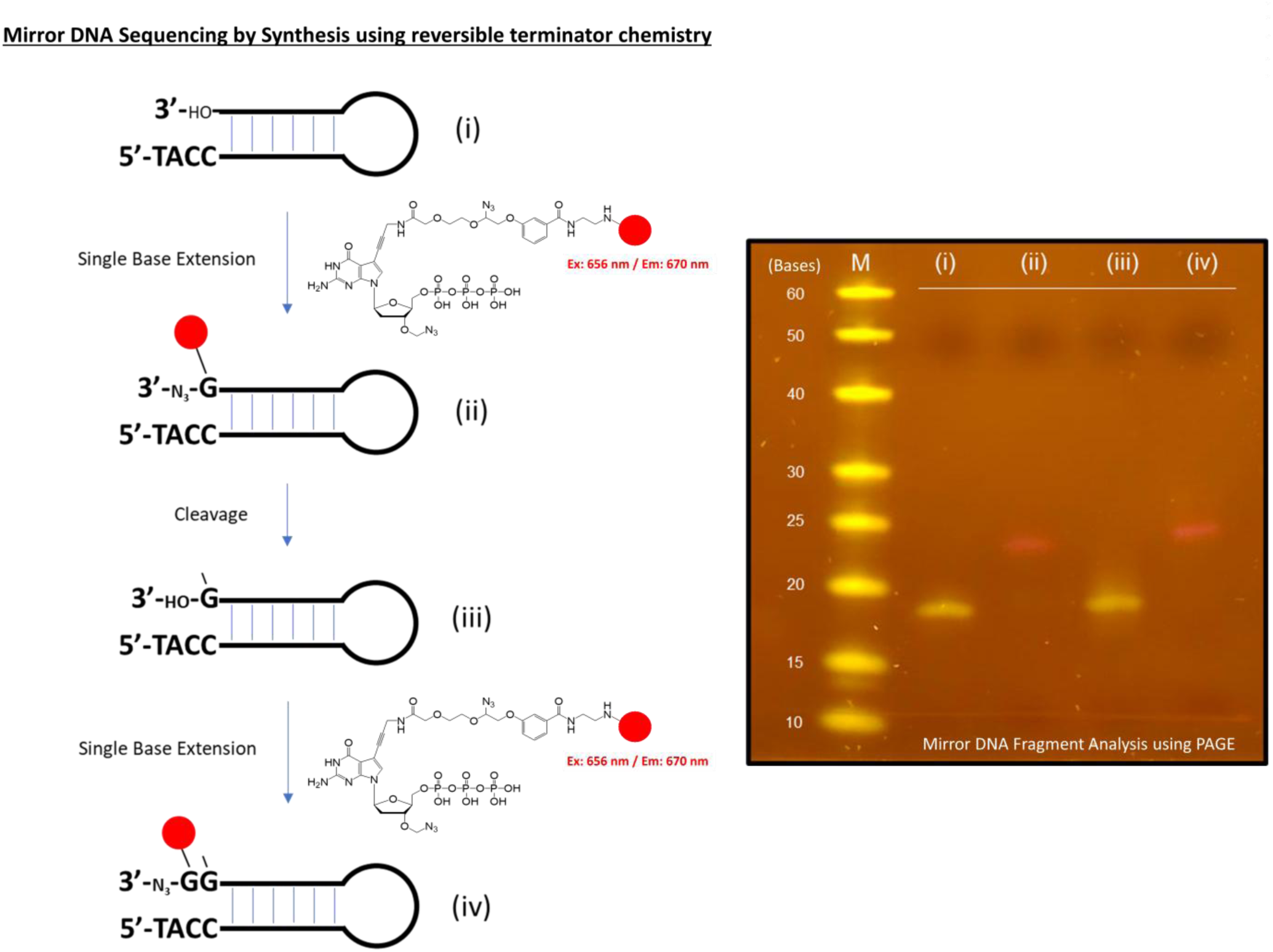

## INTRODUCTION

Molecular chirality has intrigued the scientific minds for centuries.^1,2,3^ While chirality exists universally in chemistry defining geometric properties of molecules, life on earth has evolved unequivocally as a homochiral system consisting of “right-handed” (D)-nucleic acids to encode and store genetic information in the form of RNA and DNA, accompanied by “left-handed” (L)-amino acids (except for glycine, which is achiral) that function as the building blocks for all proteins found in nature. The chirality of the biological system facilitates the folding of proteins into functional structures that allow enzymes to interact with their substrates in a precise “lock-and-key” manner, and any deviation from this (i.e., mixed-chirality life form or interaction) are not observed. The mirror image (MI) version of functional biological molecules, while mostly non-existent and not produced in nature^4,5^, are capable of being synthetically produced through chemical synthesis. Mirror image nucleic acids and peptides, or (L)-nucleic acids and (D)-peptides, can be synthesized using the same phosphoramidite and Fmoc chemistry, respectively, used to produce (D)-nucleic acids and (L)-peptides. These mirror image (L)-nucleic acids and (D)-peptides have identical physical properties to their natural counterparts, including hybridization kinetics, secondary and tertiary structure formation, but these physical properties are strictly maintained within the ‘walls’ of each chirality, meaning they are incapable of hybridization or Watson-Crick base pairing with natural (D)-nucleic acids, are completely recalcitrant to nuclease and protease recognition, and exhibit markedly reduced immunogenicity.^6^ Given these set of attributes pointing to remarkable bio-orthogonality and virtually exact physical behavior as natural biomolecules, it has long been one of the focus for scientific communities and industries needing next generation biomaterials to advance diagnostics and therapeutic solutions.

The absence of a biological system capable of directly producing large amounts of mirror-image (L)-nucleic acids and or (D)-proteins has not completely limited the advancement of synthetic versions of these MI molecules. Chemical synthesis of (D)-amino acids and (L)-nucleosides can be dated back to as early as 1854^7^ and 1964^8,9^, respectively. Since then, advancements in triphosphate, phosphoramidite, and Fmoc methodologies have fostered growth in capabilities of synthetically producing mirror-image biomolecules with increasing complexities and function^10^. In particular, construction of (D)-proteins of increasing size have been recently reported with improvements of two primary techniques that are key to producing larger (D)-proteins: first, Fmoc-based solid phase peptide synthesis (SPPS) to generate polypeptides of various lengths^11,12^ and secondly, native chemical ligations (NCL)^13,14^ to conjoin individual peptide strands generated by SPPS. Fmoc-based SPPS has evolved from a slow and manual process where small peptides (6-15 amino acids) were produced over several days or weeks, to a fully automated process. The state-of-the-art automated SPPS platforms can currently produce synthetic peptides that are 40-60 amino acids long^15^ in less than a day. With SPPS having a limit on the size of an individual peptide that it can generate, synthesis of proteins that are larger in size (60+ amino acids) require multiple peptides to be ligated to produce a full-length polypeptide using NCL. Recent advances in chemical reaction steps required for NCL, such as hydrazine-based activation to thioester-modified peptide^16^, desulfurization^17–18^, and acetamidomethyl (Acm) deprotection^19,20^ have led to product synthesis with improved final yield for longer polypeptides. Yet, proteins that are larger than ∼400 amino acids (aa) are still considered very difficult to chemically synthesize, and only a few reports of any MI proteins larger than 400 aa. Examples of full-length MI enzymes synthesized using SPPS and NCL strategy include: human immunodeficiency virus type 1 protease (HIV-1, 99 aa, 10.7 kDa)^21^, 4-hydroxy-tetrahydrodipicolinate synthase (DapA, 312 aa, 33.5 kDa)^22^, *Bacillus amyloliquefaciens* ribonuclease (barnase, 110 aa, 12.6 kDa)^23^, African swine fever virus polymerase X (ASFV pol X, 174 aa, 20.3 kDa)^24^, *Sulfolobus solfataricus* P2 DNA polymerase IV (Dpo4, 352 aa, 40.8 kDa)^25^, *Pyrococcus furiosus* DNA polymerase (Pfu, 775 aa split into 2 parts, 54.8 kDa and 35.6 kDa)^26^ and bacteriophage T7 RNA polymerase (883 aa split into 3 parts, 41.3 kDa, 26.8 kDa, and 31.5 kDa)^27^. The successful synthesis and activity shown in these works demonstrate that a total chemical synthesis strategy is entirely capable of producing MI enzymes with wide ranging sizes and activity levels virtually equivalent to its natural counterpart and sets the stage for the development of a more advanced mirror image molecular tool.

Here we report for the first time the total chemical synthesis, characterization, and functional demonstration of mirror image molecular system capable of sequencing (L)-DNA using a sequencing by synthesis (SBS) strategy. First, (L)-form deoxyguanosine, chirally inverted from natural (D)-form, was modified by using an azidomethyl moiety as a 3’-OH capping group and fluorescent dye attached to the base through a cleavable linker to produce a mirror image fluorescent nucleotide reversible terminator, (L)-3’-*O*-azidomethyl-dGTP-N_3_-fluorophore (**1**). Next, to complete the chirally inverted version of the substrate and enzyme system, we chemically synthesized a mirror image (D)-9°N mutant DNA polymerase, (775 aa split into 2 parts, 54.5 kDa and 35.5 kDa) using SPPS and NCL. The mirror image polymerase was refolded, purified, and activity verified by nucleotide incorporation assays using (L)-form nucleotide reversible terminator. Finally, proof of concept SBS of mirror image (L)-DNA template was established by confirming expected product formation at each SBS step (single nucleotide extension, fluorophore cleavage and 3’-azidomethyl deprotection, followed by 2^nd^ nucleotide extension) showing accurate mirror image sequencing by synthesis (MI-SBS) reactions through a homopolymer-C region of the (L)-DNA template. In demonstrating the ability to sequence (L)-DNA using SBS, a dominant sequencing strategy implemented in current high-throughput sequencing technologies, we have made the critical step forward to unlocking the full potential of (L)-DNA and subsequent build out of more advanced and functional mirror image molecular tools.

## RESULTS

### Synthesis of mirror image fluorescent nucleotide reversible terminator - fluorescently labeled (L)-7-deaza-7-propargylamino-3ʹ-*O*-azidomethyl deoxyguanosine triphosphate with cleavable azidomethyl linker (1)

The design principles of a reversible terminator nucleotide used in SBS-based next generation sequencing technologies are well known^28–30^. The 3’-OH capping moiety allows only a single base to incorporate during a polymerase reaction, giving time to identify the incorporated nucleotide, typically through fluorescence. Once the nucleotide identification is completed, both the 3’-OH capping moiety and fluorophore are removed to allow the polymerase reaction to continue and incorporate the subsequent base. This cyclic incorporation, detection, and cleavage, allows a base-by-base sequence identification compatible with a massively parallel nucleotide extension reaction strategy for high-throughput sequencing. To establish (L)-DNA can be sequenced using the same SBS strategy, but with a mirror version, we set out to synthesize a mirror image fluorescent nucleotide reversible terminator. Due to the lack of commercial sources of (L)-7-deaza-7-iodo deoxyguanosine **(5)**, which would typically serve as a starting synthetic precursor for nucleotide synthesis, we initially executed a glycosylation reaction with N-(4-chloro-5-iodo-7H-pyrrolo[2,3-d]pytimifin-2-yl)2,2-dimethylpropionaide **(2)** and 3,5-di-*O*-(4-methylbenzoyl)-2-deoxy-a-L-*erythro*-pentofuranosyl chloride (**3**). The glycosylation reaction was conducted with the reaction condition in the presence of powdered potassium hydroxide (KOH) and tris[2-(2-methoxy)ethyl]amine (TDA-1) as the phase transfer catalyst^31^. The resulting glycosylated product **4** was then transferred to desired precursor (L)-7-deaza-7-iodo deoxyguanosine (**5**) by incubating 2M NaOH at reflux. The precursor **5** was mixed with 2,2,2-trifluoro-N-(prop-2-yn-1-yl)acetamide to introduce a propargylamino group at 7-position by Sonogashira coupling to obtain compound **6**, followed by a TBDPS group at 5ʹ-hydroxyl group to give **7**. The 2-amine of guanine was then protected by dimethylamino group to prevent the generation of acetyl protection when introducing methylthiomethyl (MTM) group at 3ʹ-OH. After the introduction of MTM group on 3ʹ-OH of **8** in the condition of DMSO, AcOH and Ac_2_O, the replacement of thiomethyl group with azido group was achieved through the following steps: (1) *in situ* activation of the thiomethyl group with sulfuryl chloride and (2) nucleophilic replacement with NaN_3_ in dimethylforamide. The resulting product **10** was then treated with 1M TBAF in THF to obtain the intermediate **11**. With this intermediate **11**, we triphosphorylated^32,33^ at 5’-hydroxyl position to give **12** in 21% yield over four steps and then attached a cleavable azidomethyl linker (**S-3**) (Supplementary scheme 1) at 7-propargylamino group to complete **13** with a 66% yield. The commercial fluorophore (iFluor647-NHS ester, AAT Bioquest) was attached to **13** by a simple coupling reaction to give the desired fluorescently labeled (L)-7-deaza-7-propargylamino-3ʹ-*O*-azidomethyl deoxyguanosine triphosphate with cleavable azidomethyl linker (**1**, (L)-3’-*O*-azidomethyl-dGTP-N_3_-fluorophore).

### Design rationale for the selection of (D)-form 9°N DNA polymerase variant for mirror image SBS

To develop a high-throughput mirror image sequencing by synthesis platform, we needed a polymerase capable of incorporating highly modified nucleotide analogues. We initially started with a 9°N DNA polymerase variant that contained the critical aa substitutions (D141A, E143A, Y409V and A485L, Fig. 1B) well established for its ability to recognize and faithfully incorporate 3ʹ-*O*-azidomethyl-modified nucleotides. These set of aa substitutions provide key functional enhancements (gain-of-function mutation in the finger domain (Y409V, A485L) and inactivation of 3’-5’ exonuclease activity (D141A, E143A) critical for efficient incorporation of highly modified and bulkier nucleotides. Our objective was to retain this polymerase function by minimizing aa sequence changes while recognizing a handful of aa residue substitutions would be required to chemically synthesize a polymerase of this size.

**Figure 1.**
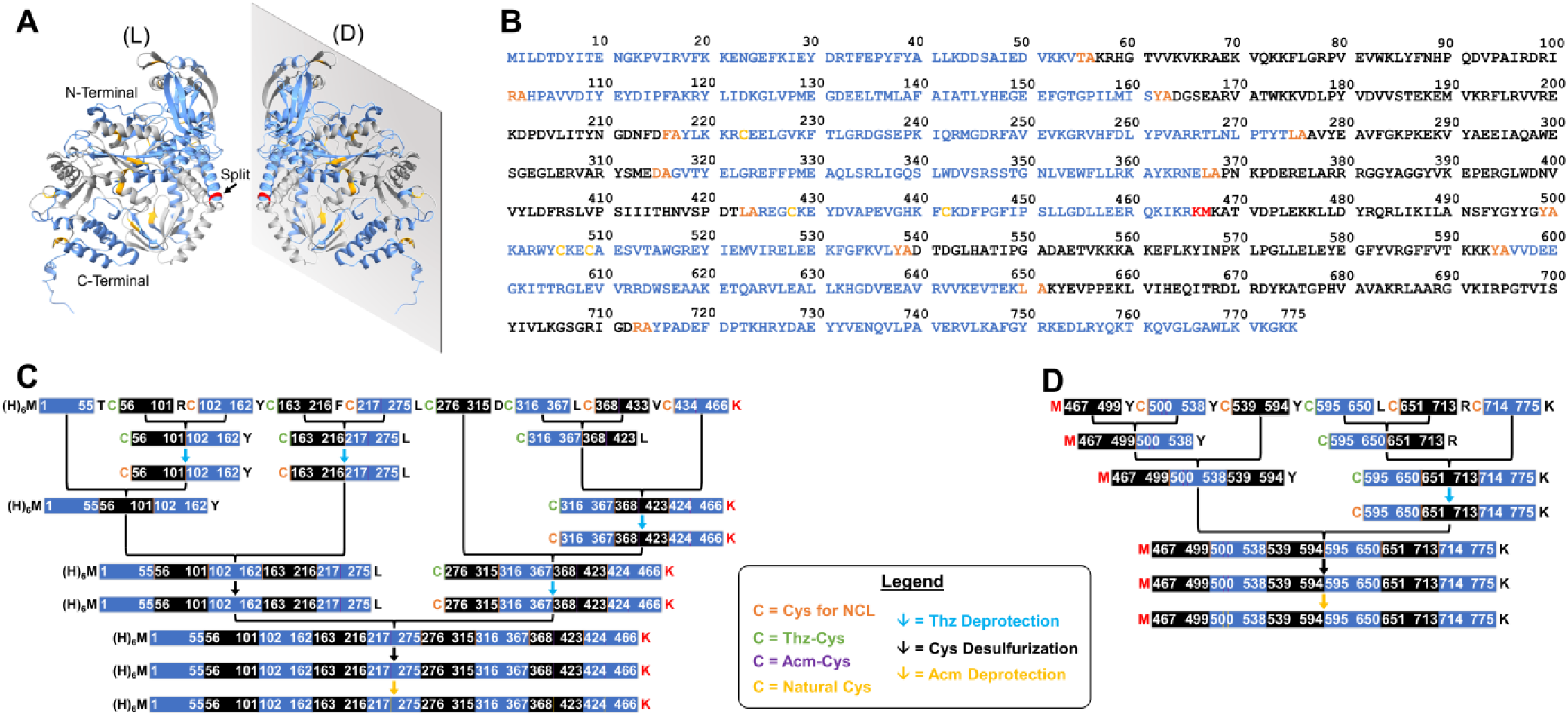
Synthesis assembly of mirror image (D)-9°N DNA polymerase using native chemical ligation (NCL). Ribbon structure representation (A) of (L)- and (D)-9°N DNA polymerase variant 1 indicating alternating fragmentation of the amino acid sequence, native chemical ligation junction sites, and the split location of the N- and C terminal of the polymerase in an alpha-helix. (B) Full amino acid sequence of (D)-9°N DNA polymerase variant 1 indicating all junction sites (in orange), split location (in red) and natural cysteines (in yellow). Individual peptide fragments ranged between 33-63 amino acids in length (highlighted in alternating blue and black fonts). (C) Native chemical ligation synthesis scheme for the N-terminal (466 amino acids) and (D) C-terminal (309 amino acids) peptides. Assembly scheme depicts all Thz-Cys deprotection, cysteine desulfurization and Acm- deprotection to recover natural cysteines post-NCL.

Next, we took advantage of series of both established and emerging methods that enable effective synthesis of large proteins (>500 aa) chemically. First, to identify potential ligation junctions between peptides for native chemical ligation (NCL), we identified the site of every natural cysteine (C223, C428, C442, C506, C509, Fig. 1B) across the entire 9°N DNA polymerase (total length of 775 aa). NCL requires an amino-terminal thiol functional group, typically cysteine, to perform a transthioesterification between peptides generated via SPPS^11,12^. However, due to the size of the polymerase, the five natural cysteine sites would require synthesis of large peptides with sizes up to 267 aa, which is not feasible and practical in general SPPS due to the significant drop off in crude purity and yield with larger peptides. We then took advantage of the ability to convert unprotected cysteine back to alanine (i.e. post-NCL reaction) via a radical desulfurization reaction^17^, leading to 54 additional possible ligation junction sites that led to individual peptide length in the range of 2 to 49 aa. We also performed a proactive systematic mutation of non-conserved isoleucine residues, a bulky hydrophobic aa, into less bulky and more neutral aa to better navigate through solubility difficulties during NCL^34,35^. With these parameters in mind, we designed a retrosynthesis scheme of the (D)-9°N DNA polymerase of the most advantageous junctions^14^ for NCL with mutation of amino acid sites (E276A and N424A, including isoleucine mutations at I80V, I127V, I171A, I176V, I191V, I228V, I256V, I264A, I268L, I400V, I597V, I610V, I618A, I630L, I642V, I715Y, I733V, I744V)^26^ that differs from the published 9°N DNA polymerase sequence. In total, the (D)-9°N DNA polymerase was fragmented into 15 peptide strands ranging from 33-63 amino acids (Fig. 1C and D) (D-9°N-N-1 to -9 and D-9°N-C-1 to -6) that will allow us to build the full protein in two large halves (N-terminal and C-terminal fragments)^26,36^, with a final non-covalent assembly process to produce a functional polymerase. As an additional confirmation of the final design, we recombinantly produced a natural (L)-form 9°N polymerase with the same aa sequence (Supplementary Fig. 1 and 37) and verified comparable activity (Supplementary Fig. 40).

### Synthesis of (D)-form mutant 9°N DNA polymerase via SPPS and NCL

We utilized key orthogonal protecting groups: (S)-thiazolidine (Thz)^19,37,38^ and S-acetamidomethyl (Acm)^19,20^ to facilitate parallel NCL reactions and preservation of natural cysteine, respectively. While cysteine is naturally less abundant in protein sequences, they play an outsized role in protein folding and stability via disulfide bridge formation. Preservation of key physiochemical components that lead to accurate protein refolding and activity guided our design and synthesis procedure. With the scheme in hand, we synthesized the starting peptides using Fmoc chemistry with an attached hydrazine^16^, a thioester synthon, on the C-terminal of each designated peptide (D-9°N-N-1 to -8; Supplementary Fig. 2-9 and D-9°N-C-1 to -5; Supplementary Fig. 22-26) by SPPS on CEM Liberty Blue 2.0^TM^. The hydrazine is a robust reactive group, which can be easily converted to a thioester for NCL reactions, and a compatible group under NCL reaction conditions. D-9°N-N-9 (Supplementary Fig. 10) and D-9°N-C-6 (Supplementary Fig. 27) were synthesized with C-terminal carboxylic acid, since C-terminal ligations were unnecessary. All peptides were purified by Reverse Phase High Performance Liquid Chromatography (RP-HPLC) and mass confirmed by Liquid Chromatography–Mass Spectrometry (LC-MS).

After the synthesis of all starting peptides (D-9°N-N-1 to -9 and D-9°N-C-1 to -6), the intermediate protein products were assembled using NCL (peptidyl-thioester and Cys-peptide). In general, the peptidyl hydrazine was first activated by acetylacetone (Acac) and 4-mercaptophenylacetic acid (MPAA) to generate an activated peptidyl-MPAA, then the second peptide (Cys-peptide) was *in situ* added to the solution of activated peptydyl-MPAA. All the NCL reactions were performed at room temperature, monitored and analyzed by analytical RP-HPLC and LC-MS and further purified by preparative RP-HPLC (N-terminal; Supplementary Fig.11-21 and C-terminal; Supplementary Fig. 28-34). The activation method (Acac and MPAA) was chosen instead of the common activation method (Sodium nitrite (NaNO_2_) and MPAA) because the thiazolidine group is incompatible^39,40^ under NaNO_2_ activation. Two rounds of complete synthesis procedure yielded a total of 19.1 mg of D-9°N-N-20 (calculated/observed; 54541.58/54541.42 Da) and 31.2 mg of D-9°N-C-13 (calculated/observed; 35524.27/35525.26 Da) (Fig. 2A).

**Figure 2.**
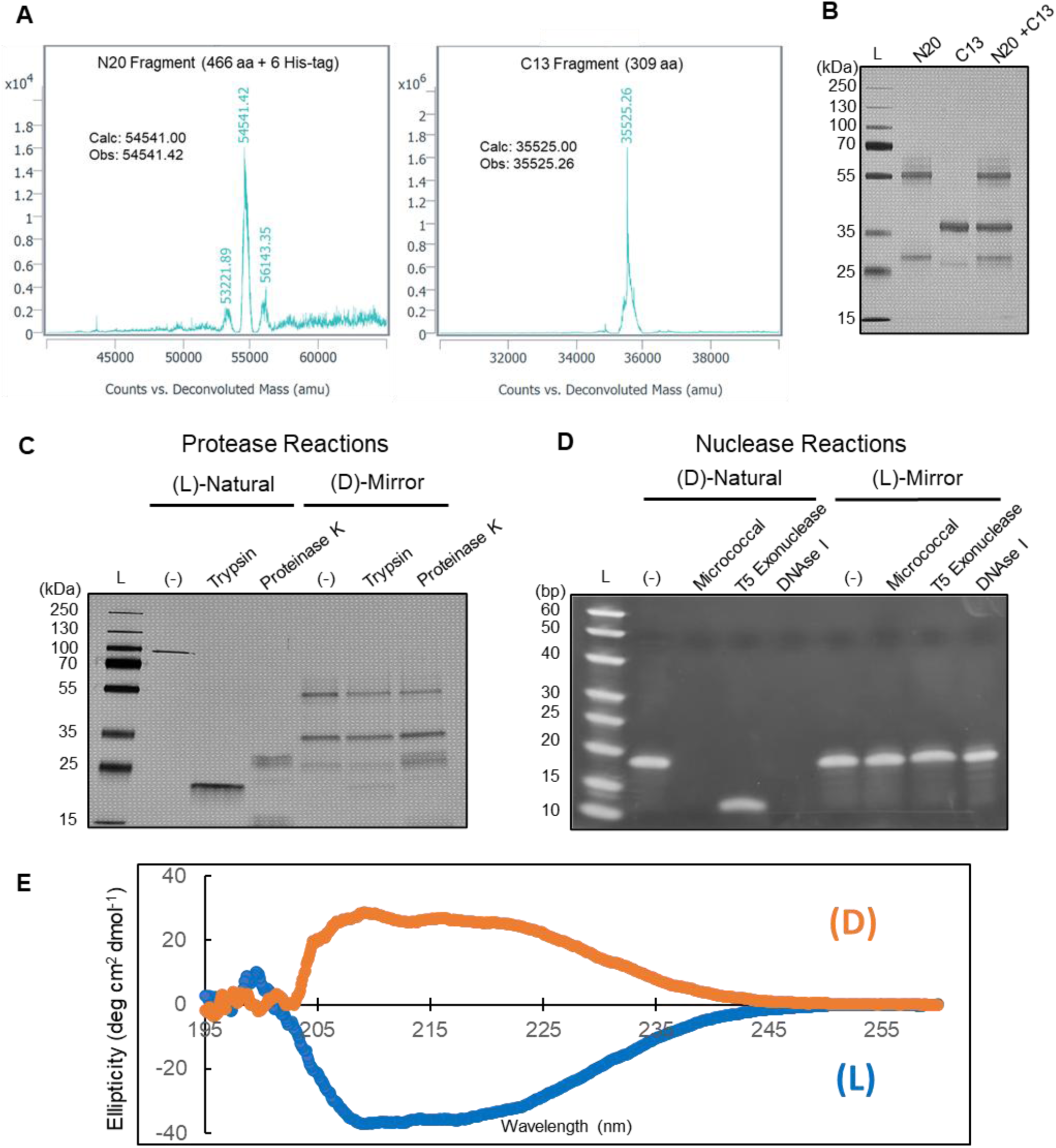
Characterization of synthetic (D)-9°N DNA polymerase variant 1. (A) LC-MS analysis of the N20 and C13 fragments indicating full length products. (B) SDS-PAGE analysis of starting N20 (lane 2), C13 (lane 3) and co-refolding of N20 and C13 post heat shock (lane 4). (C) Protease degradation analysis via SDS-PAGE of (L)-9°N DNA polymerase (lane 1) in the presence of Trypsin (lane 2), Proteinase K (lane 3) and (D)-9°N DNA polymerase (lane 4) in the presence of Trypsin (lane 5) and Proteinase K (lane 6). (D) Demonstrating the stability of (L)-DNA against nucleases. Natural (D)-Oligonucleotide template was incubated with DNAse I (lane 1), lambda (lane 2), T7 exonuclease (lane 3) versus the (L)-Oligonucleotide template in lanes 4-6, respectively. (E) Circular dichroism analysis of (L)-9°N DNA polymerase (blue) and (D)-9°N DNA polymerase variant 1 (orange).

### Final assembly and characterization of mirror image (D)-form mutant 9°N DNA polymerase

Previous work has established that under the right conditions and design, intermolecular interaction can drive multiple smaller protein fragments to refold into its singular full-size protein maintaining both structural accuracy, stability and activity^36^. We performed the final assembly between D-9°N-N-20 and D-9°N-C-13 fragments through gradual dialysis procedure changing the medium from a highly denaturing environment to long-term storage compatible buffered environment to encourage accurate refolding into its proper active structure. In further assessing correct refolding, we took advantage of the fact that a properly refolded polymerase must be heat-proof, especially if they are thermostable, which is true for 9°N DNA polymerase. We confirmed using SDS-PAGE analysis that after incubation at 90°C for 15 minutes, incorrectly folded proteins were precipitated and only the correctly folded products remained in-tact (Fig. 2B).

To demonstrate the bio-stability of the (D)-9°N DNA polymerase, we performed a protease degradation reaction using trypsin and proteinase K. The SDS-PAGE analysis shows the inability of the proteases to degrade mirror image (D)-9°N DNA polymerase (Fig. 2C, Supplementary Fig. 38). As verification that a similar bio-stability exists for (L)-DNA, a nuclease degradation reaction was performed using DNAse I, lambda (λ), and T7 exonuclease. As shown in Fig. 2D, (L)-DNA is recalcitrant to nuclease degradation, while full or partial degradation of (D)-DNA is complete within 15-30 minutes. Even in reactions where (L)-DNA is incubated with excess nucleases, it withstands these extreme cases with remarkable stability (nuclease titration, Supplementary Fig. 36). Finally, optical measurements of both (D)- and (L)-form 9°N DNA polymerase was performed using circular dichroism (CD) spectroscopy, verifying the chirality of the synthesized products. As shown in Fig. 2E, mirror image (D)-polymerase has optical measurements that are reversed in intensity (Fig. 2E, orange-line) compared to the natural (L)-polymerase (Fig. 2E, blue-line).

### Verification of mirror image (D)-9°N DNA polymerase activity by nucleotide incorporation assay using non-fluorescent (L)-3ʹ-*O*-azidomethyl-dGTP

To validate (D)-9°N DNA polymerase variant with specific mutations outlined in this work was refolded properly and functional, the polymerase was challenged to accurately incorporate a non-fluorescent (L)-3ʹ-*O*-azidomethyl-dGTP (**S-9**, Supplementary scheme 2) into a self-priming (L)-DNA hairpin template in solution. The extended product was analyzed using Matrix-Assistant Laser Desorption/Ionization–Time of Flight (MALDI-TOF) mass spectrometry (MS) enabling high-resolution accurate molecular product determination. As shown by the mass measurement of the extended product at m/z 8335.826 (calculated m/z 8335, Supplementary Fig. 39), (L)-3’-*O*-azidomethyl-dGTP (**S-9**) was found to efficiently incorporate into the growing (L)-DNA strand in a base specific manner and subsequently terminated the DNA synthesis to allow for base identification. This result indicates that the aa changes that were incorporated into the mirror image (D)-9°N DNA polymerase did not have detrimental effect toward polymerase activity and specificity. As comparison, an extension reaction using a recombinantly expressed (L)-9°N DNA polymerase variant and (D)-3’-*O*-azidomethyl-dGTP (TriLink) was carried out and analyzed using MALDI-TOF MS, and found to have similar activity to its mirror image counterpart m/z 8350.957 (calculated m/z 8350, Supplementary Fig. 40).

### Demonstration of MI-SBS using fluorescent (L)-3ʹ-*O*-azidomethyl-dGTP-N_3_-flurophore and (D)-9°N DNA polymerase

Finally, to verify that mirror image fluorescent nucleotide analogues can be used for MI-SBS, two continuous steps of (L)-DNA extension and cleavage were carried out in solution using (L)-3’-*O*-azidomethyl-dGTP-N_3_-fluorophore (**1**) and a self-priming (L)-DNA template (5’-<u>GACC</u>GCGCCGCGCCTTGGCGCGGCGC-3’). The four nucleotides in the template immediately adjacent to the annealing site of the primer are underlined. This experimental design allows the isolation of the (L)-DNA product at each step for detailed molecular structure characterization by MALDI-TOF MS as shown in Fig. 3. The first extension product (L)-5’-G(N_3_-fluorophore)-3’-O-azidomethyl (Fig. 3A, ii) was purified by HPLC and analyzed using MALDI-TOF MS (Fig. 3B, ii). This product was then incubated in aqueous TCEP solution^41,42^ to simultaneously cleave both the fluorophore and the 3’-*O*-azidomethyl group. The cleaved product (Fig. 3A, iii) was also analyzed by using MALDI-TOF MS (Fig. 3B, iii, m/z 8435). As can be seen in Fig. 3B, ii, the MALDI-TOF MS spectrum consists of a distinct peak at m/z 9692 corresponding to the (L)-DNA extension product (L)-5’-G(N_3_-fluorophore)-3’-O-azidomethyl, which confirms that the nucleotide analogue can be accurately incorporated as base terminator by mirror image DNA polymerase into a growing DNA strand. Fig. 3B, iii shows the cleavage result of the above extended (L)-DNA product. The peak at m/z 9692 has completely disappeared, whereas the peak corresponding to the dual cleaved product (L)-5’-G-3’-OH appears as the sole dominant peak at m/z 8435, which establishes that the incubation with TCEP completely cleaves both the fluorophore and the 3’-*O*-azidomethyl group with high efficiency without damaging the (L)-DNA. The next extension reaction was carried out by using this cleaved (L)-DNA product (Fig 3A, iii) with a free 3’-OH group regenerated as a primer along with (L)-3ʹ-*O*-azidomethyl-dGTP-N_3_-fluorophore (**1**) to yield an extension product (L)-5’GG(N_3_-fluorophore)-3’-O-azidomethyl. This extension product (Fig 3A, iv) was purified by HPLC and analyzed using MALDI-TOF MS producing a dominant peak at m/z 10176 (Fig. 3C, iv). These results demonstrate that the above-synthesized chemically cleavable fluorescent nucleotide analogue is successfully incorporated with high fidelity into the growing DNA strand in a polymerase reaction, and furthermore, both the fluorophore and the 3’-*O*-azidomethyl group are efficiently removed by using a mild and short incubation with TCEP, which makes it feasible to use them for SBS of (L)-DNA.

**Figure 3.**
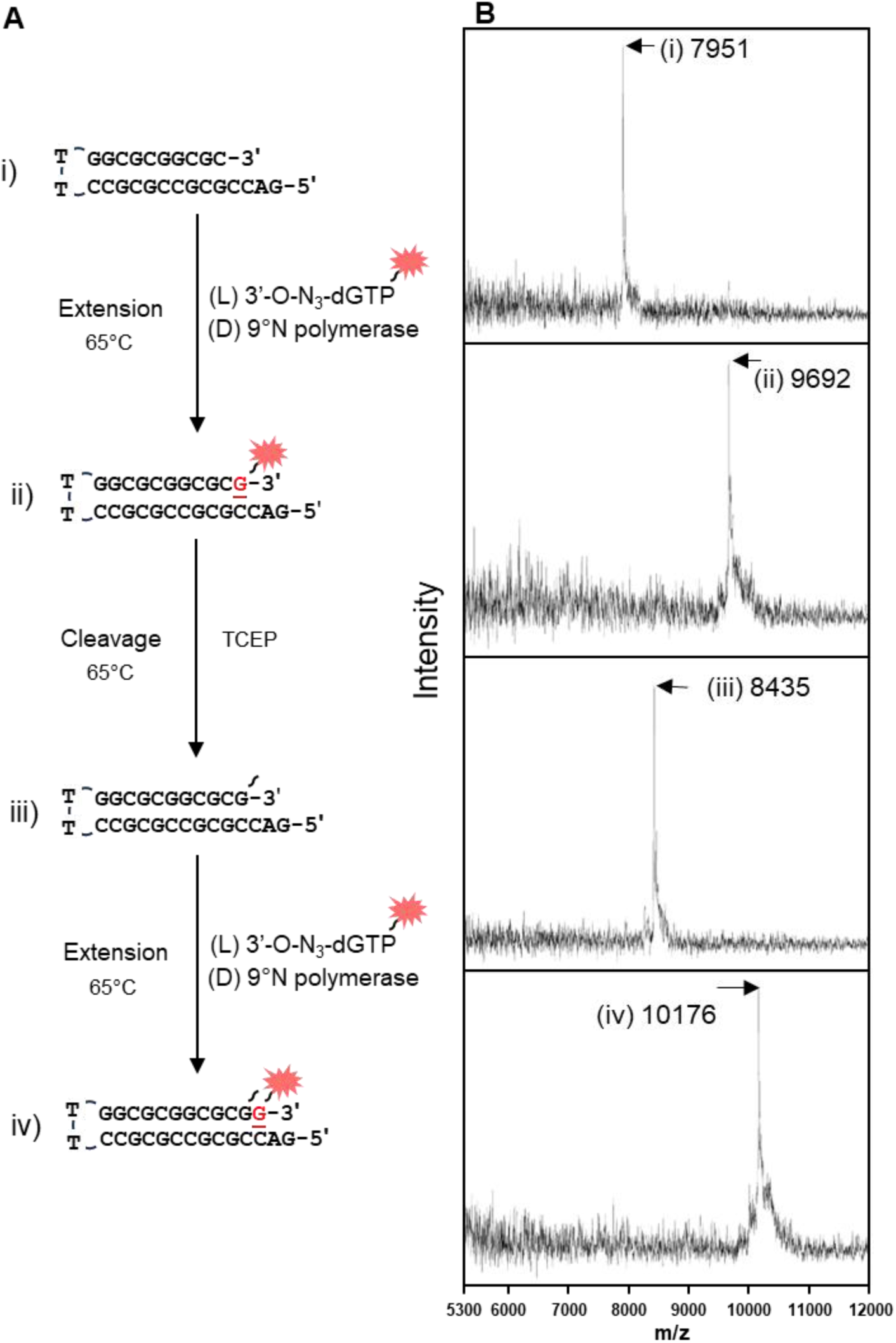
Iterative incorporation of fluorescent (L)-3ʹ-*O*-azidomethyl-dGTP with (D)-9°N DNA polymerase. (A) Primer extension scheme starting with a self-priming (L)-DNA template (i) incubated with fluorescent (L)-3ʹ-*O*-azidomethyl-dGTP and (D)-9°N DNA polymerase variant 1 at 65°C. Reactions were purified by RP-HPLC and (B) analyzed by MALDI-TOF MS to validate the incorporation of the fluorescent (L)-3ʹ-*O*-azidomethyl-dGTP (ii). The fluorescent product was cleaved by TCEP at 65°C for 30 minutes, mass confirmed by MALDI-TOF MS and utilized as the starting template (iii) for the second incorporation (iv).

## DISCUSSION

Beyond sequencing genomes and transcriptomes, high-throughput DNA sequencing has transformed both biomedical research and human health by providing the capability of hypothesis-free experimentation at scale. Powerful technologies such CRISPR screens, saturation mutagenesis, aptamer/antibody selection, and rapid feedback on designed molecules have advanced on account of NGS sequencing advancements.^43,44^ SBS chemistry and methods have been the engine beneath the continuous improvement of DNA sequencing throughput over the past two decades because of the ability to readout sequences in a massively parallel format. More advanced technologies and new biological insights will undoubtedly be achieved as throughput power improves. In this work, we have chemically synthesized the reagents necessary to develop a MI-SBS system to enable high-throughput mirror image (L)-DNA sequencing. The two prerequisite materials to perform SBS on (L)-DNA are i) a set of fluorescently labeled, 3ʹ-*O*-azidomethyl protected (L)-form nucleotide triphosphates and ii) a DNA polymerase constructed from D-amino acids capable of incorporating the mirror image nucleotide analogues. First, we successfully synthesized the fluorescent-(L)-3ʹ-*O*-azidomethyl-dGTP (**1**) starting from a glycosylation of a required modified 7-iodinated guanine base (**2**) and a (L)-1-Cl-3,5-protected-deoxyribose (**3**), followed by installation of a reversible terminator group (azidomethyl) and a cleavable azidomethyl linker tethered with a fluorophore. The azidomethyl functional group on both 3ʹ-OH and the linker were designed as a reversible terminator to allow for single nucleotide incorporation and detection, respectively.^41,42^ Utilization of the azidomethyl group allows for rapid and simultaneous chemical removal of the 3ʹ-*O*-cap and fluorophore with TCEP. We accomplished the synthesis and verification of the second prerequisite for MI-SBS - a mirror image (D)-9°N DNA polymerase variant capable of faithful incorporation of the heavily modified mirror image nucleotide reversible terminator. The functional (D)-9°N DNA polymerase variant was successfully constructed using SPPS and NCL, demonstrating feasibility of mirror image molecular engineering using total chemical synthesis and cell-free refolding process that can potentially scale with further optimization across various chemical manufacturing process. Taken together, we show the ability to create an advanced mirror image molecular system, in this case a MI-SBS system, which will lead to advancements in molecular tools and technologies based on mirror image nucleic acids.

In an effort to complete the development of MI-SBS especially for broad access, the synthesis of the complete set of fluorescent nucleotide reversible terminators (dA, dC, and dU) must be finished. We started with the synthesis of (L)-3’-*O*-azidomethyl-dGTP-N_3_-fluorophore (**1**), described in this work, because it required the synthesis of a precursor, (L)-7-deaza-7-iodo deoxyguanosine (**5**), a compound that had not been previously described or reported. Glycosylation of iodinated purine-base and (L)-1-Cl-3,5-protected-deoxyribose (**3**) led to successful production of precursor and establishes a valid synthesis path for (L)-3’-*O*-azidomethyl-dATP-N_3-_fluorophore. The remaining pyrimidine-based (L)-3’-*O*-azidomethyl-dCTP-N_3_-fluorophore and (L)-3’-*O*-azidomethyl-dUTP-N_3_-fluorophore will follow a more traditional and established synthetic path.^45^ The complete set of the fluorescent nucleotide reversible terminators (dA, dC, dG and dU) are currently in the final stages of synthesis, and when complete, will be described elsewhere. Additionally, development in the area of scalable mirror image protein (>500 aa) process in a cost-effective manner will be paramount and we have taken initial steps at improving the yield and cost structure associated with mirror image polymerase production. In the SPPS front, we have found yield improvement from many factors, including from non-obvious sources such as vendor specific resin types and loading capacity^46^, to more straightforward synthesis scale increases. During our initial nominally increased synthesis scale, we have observed up to 41% increase in final product yield and 49% decrease in reagent cost simply starting the SPPS at a higher scale (0.1 mmol to 0.4-0.8 mmol). During NCL, in addition to ligation junction selection^14^, general improvement and optimization in peptide solubility, separation of ligated and un-ligated products, strategic protection and deprotection designs/methods have led to significant improvement in the final product yield, but as we work with many different polymerase variants with varying aa sequences, data-driven systematic optimization process tailored for ‘tough’ aa sequence regions must be derived and able to be utilized in a predictable manner.

Mirror image molecular systems hold amazing promise for medicine and bioengineering applications because they are bio-orthogonal and do not interact with natural DNA or proteins, including nucleases and proteases. While bio-orthogonality yields many advantages, development of mirror image molecular systems has thus far been limited due to the absence of an advanced toolkit to manipulate and analyze these molecules. Previous attempts at analyzing (L)-DNA by sequencing utilized a Sanger-based method using error-prone (D)-Dpo4 polymerase variant.^47^ This method is inherently low-throughput and typically cannot support applications and experiments requiring large amount of sequencing data. With a development of a high-throughput mirror image sequencing, many new technology and applications using mirror image DNA can proceed with expanded complexity. For example, high-throughput discovery of (L)-DNA based aptamers can be accomplished using the full power of in-vitro selection methodologies such as Systematic Evolution of Ligands by Exponential Enrichment (SELEX)^48,49^ where the starting unique number of DNA sequences can exceed 10^14^. Our MI-SBS technology uniquely enables direct mirror image (L)-DNA SELEX (MI-SELEX), producing aptamers that are bio-orthogonal to endogenous DNA and proteins making them nuclease resistant and free from off-target binding. MI-SELEX can provide rapid, efficient, and scalable selection and characterization of mirror-image aptamers without needing mirror-image protein targets, fundamentally transforming (L)-DNA aptamer discovery. Thus, previously inaccessible molecular targets become feasible, and the efficiency and speed of aptamer development improve dramatically. Furthermore, massively parallel array format utilized by SBS can lead to multiplexed structure-function profiling where highly multiplexed characterization of binding activities and sequencing data can be obtained within clonal (L)-DNA populations to yield large data sets required to train machine learning and artificial intelligence algorithms for de novo design and prediction. This powerful capability allows direct generation of comprehensive sequence-function datasets, streamlining translation into precise structure-function relationships. Such integration accelerates identification and validation of therapeutic aptamers.

MI-SBS technology is a major step forward in our ability to make useful mirror image DNA systems as we can now read the DNA sequence in a high-throughput manner in addition to writing it. This enables numerous uses of (L)-DNA systems that require properties such as their orthogonality towards natural nucleic acids and nuclease resistance including as mirror image DNA data storage to alleviate cold and sterile storage requirements and serve as an ultra-secure stealth (undetectable, unreadable, ultra-secure) information storage system, rapid mirror-image aptamer selection for ‘Print-on-Demand’ nuclease resistant precision diagnostics and therapeutics, and bio-orthogonal DNA barcode tags for *in situ* and *in vivo* applications such as spatial biology and multiplexed drug screening.

**Scheme 1.**
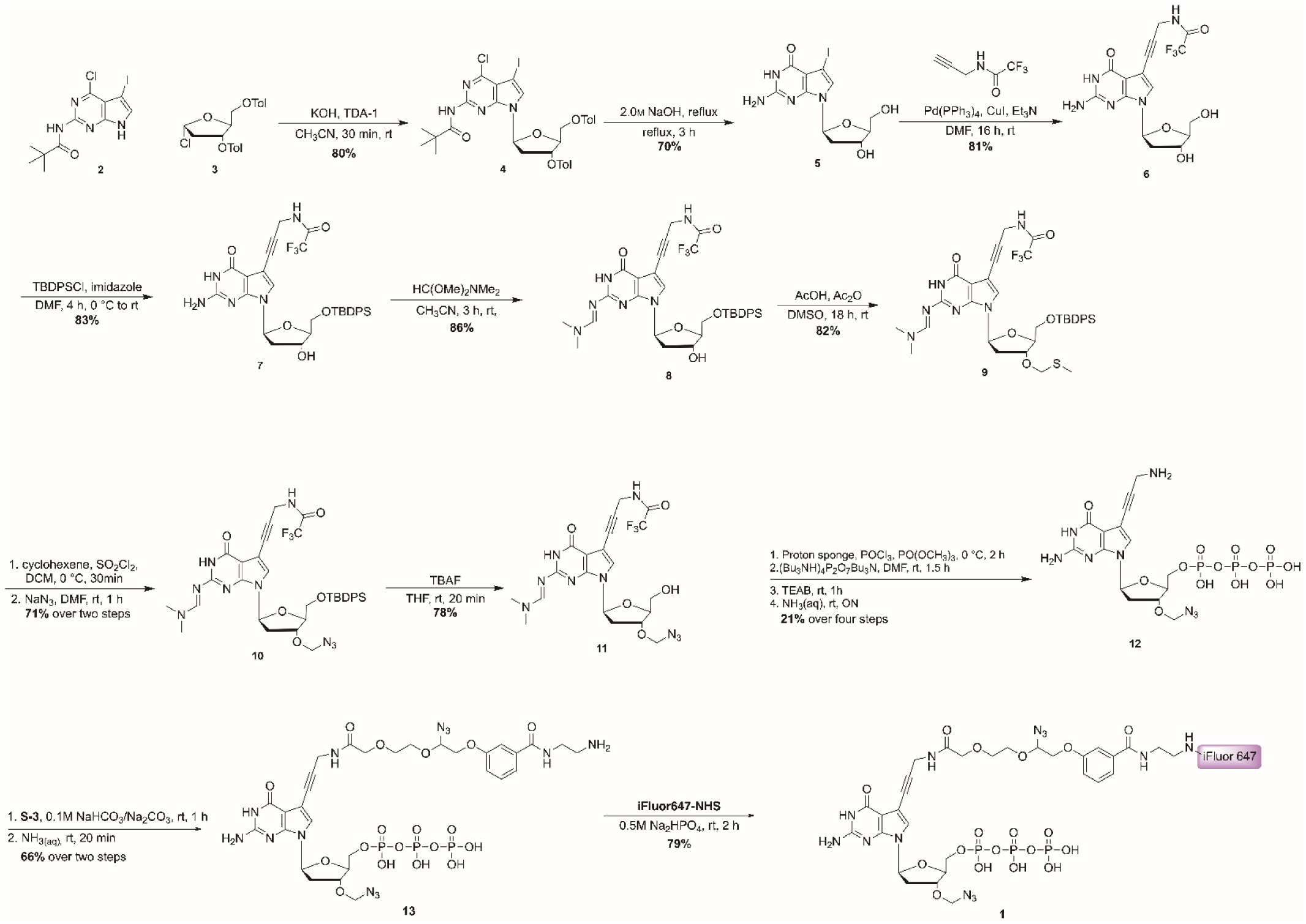
Synthesis scheme of fluorescently labeled (L)-3ʹ-*O*-azidomethyl deoxyguanosine triphosphate with cleavable azidomethyl linker **3**.

## METHODS

### Materials

All solvents and reagents for chemical synthesis were reagent grades, purchased commercially, and used without further purification unless specified. Chemicals without specific indication for synthesis of compound **3** were purchased from Sigma-Aldrich. All other chemicals were purchased from suppliers as described below. Triethylamine was purchased from Thermo Fisher Scientific. N-(4-Chloro-5-iodo-7h-pyrrolo[2,3-d] pyrimidin-2-yl)-2,2-dimethylpropionamide **(2)** was purchased from Combi-Blocks (San Diego, CA). 1-Chloro-2-deoxy-3,5-di-*O*-toluoyl-L-ribofuranose (**3**) and 2,2,2-trifluoro-N-(prop-2-yn-1-yl)acetamide were purchased from Ambeed (Buffalo Grove, IL). 1M Triethylammonium bicarbonate buffer (pH 8.5, HPLC grade) was purchased from Boston Bioproducts (Milford, MA). iFluor647 NHS ester were purchased from AAT Bioquest (Sunnyvale, CA). (D)-Oligonucleotide G was purchased from IDT (Coralville, IA) and (L)-Oligonucleotide G was purchased from ChemGenes (Wilmington, MA).

### Synthesis of fluorescently labeled (L)-3ʹ-*O*-azidomethyl deoxyguanosine triphosphate with cleavable azidomethyl linker (1)

#### 3’*-O-*azidomethyl-7-deaza-7-[3-acetamido(2-(2-(2-(3-((2-aminoethylcarbamoyl)phenoxy)-1-adizoethoxy)ethoxy)-prop-1-ynyl]-L-2’-deoxyguanosine 5’-*O*-triphosphate iFluor647 conjugate (1)

To a solution of **13** (1.9 mg, 0.0020 mmol) in 0.5M Na_2_HPO_4_ (100 uL) was added a solution of iFluor647-NHS (5.0 mg, 0.0039 mmol) in DMF (10 uL) and stirred at room temperature for 2 hours. The reaction was monitored by HPLC until **2** was consumed completely. The crude was purified on preparative HPLC (Polaris C18, 10.0 x 250 mm, 180Å, 5 mm) with gradient 0%B to 90%B over 50 mins (A: 0.1M TEAB, B: ACN) to obtain **1** (3.2 mg, 79%). HRMS (ESI): m/z [M] calcd 1919.5; Found [M – 3H]^3–^ 638.8356. Compound **1** was further evaluated by performing a single base extension (MALDI-TOF: m/z 9692), cleavage of cleavable linker (MALDI-TOF: m/z 8435) and another single base extension (MALDI-TOF: m/z 10176). (Fig. 3).

#### 6-chloro-7-deaza-7-iodo-L-3’,5’-di-*O*-toluoyl-2’-deoxyguanosine (4)

To suspension of KOH (680 mg, 8.71 mmol) and TDA-1 (0.1 mL, 0.32 mmol) in CH_3_CN (30 mL), N-(4-Chloro-5-iodo-7h-pyrrolo[2,3-d] pyrimidin-2-yl)-2,2-dimethylpropionamide (**2**) (756 mg, 2.00 mmol) was added at room temperature. After stirring for 10 min, 1-chloro-2-deoxy-3,5-di-*O*-toluoyl-L-ribofuranose (**3**) (1.02 g, 2.6 mmol) was added. The reaction mixture was stirred at room temperature for 30 minutes. The insoluble material was filtered off, and the precipitate was washed with 3 times of CH_3_CN (15 mL x 3), and the resulting filtrate was evaporated to dryness. The residue was purified by column chromatography (100% CH_2_Cl_2_) to obtain **4** (1.15 g, 80%) as brown foam. ^1^H NMR (400 MHz, CDCl_3_) δ 8.21 (s, 1H), 8.00 (d, *J* = 7.9 Hz, ArH), 7.93 (d, *J* = 7.9 Hz, ArH), 7.45 (s, 1H, H-8), 7.27 (app t, *J* = 7.9 Hz, ArH), 6.74 (dd, 1H, *J* = 6.2 Hz, *J* = 7.6 Hz, H-1’), 5.81 (dt, 1H, *J* = 2.7 Hz, *J* = 6.7 Hz, H-3’), 4.77 (dd, 1H, *J* = 3.8 Hz, *J* = 12.0 Hz, H-5’a), 4.67 (dd, 1H, *J* = 4.0 Hz, *J* = 12.0 Hz, H-5’b), 4.61–4.58 (m, 1H, H-4’), 3.02–2.93 (m, 1H, H-2’a), 2.83 (ddd, 1H, *J* = 2.7, 6.1, 14.3 Hz, H-2’b), 2.45 (s, 3H, C*H_3_*Ph), 2.44 (s, 3H, C*H_3_*Ph) 1.37 (s, 9H, CH_3_ of *t*Bu); ^13^C NMR (100 MHz, CDCl_3_) δ 175.75, 166.27, 166.21, 152.97, 151.69, 151.47, 144.51, 144.25, 131.17, 130.01, 129.74, 129.51, 129.37, 129.26, 126.85, 126.57, 113.97, 84.73, 82.81, 75.09, 64.11, 60.51, 53.25, 40.44, 38.14, 27.54, 21.88, 21.86, 14.34; HRMS (ESI) m/z [M] calcd for C_32_H_32_ClN_4_O_6_ 730.1055; Found [M + H]^+^ 731.1159.

#### 7-deaza-7-iodo-L-2’-deoxyguanosine (5)

Compound **4** (319 mg, 0.437 mmol) was treated with NaOH solution (2.0 M, 5.0 mL) at room temperature. The solution was heated to reflux for 3 hours. After completion of **4**, the mixture was cooled to room temperature and neutralized by addition of 2.0M HCl to pH = 5. The mixture was then evaporated to dryness. The residue was purified by column chromatography (100% CH_2_Cl_2_ to 85:15 CH_2_Cl_2_– CH_3_OH) to give **5** (120 mg, 70%) as yellowish foam. ^1^H NMR (400 MHz, CD_3_OD) δ 7.00 (s, 1H, H-8), 6.27 (dd, 1H, *J* = 6.2 Hz, *J* = 8.0 Hz, H-1’), 4.37–4.32 (m, 1H), 3.85–3.80 (m,1H), 3.64 (dd, 1H, *J* = 3.7, 11.7 Hz, H-5’a), 3.58 (dd, 1H, *J* = 4.3, 11.7 Hz, H-5’b), 2.38 (ddd, 1H, *J =* 6.2, 8.0, 14.2 Hz, H-2’a), 2.14 (ddd, 1H, *J =* 3.2, 6.2, 14.2 Hz, H-2’b); ^13^C NMR (100 MHz, CD_3_OD) δ 161.30, 154.14, 152.51, 124.84, 102.37, 88.76, 85.34, 72.98, 63.68, 54.85, 41.42, 27.78; HRMS (ESI) m/z [M] calcd for C_11_H_13_IN_4_O_4_ 391.9981; Found [M + H] ^+^ 393.0067.

#### 7-deaza-7-[3-(2,2,2-trifluoroacetamido)-prop-1-ynyl]-L-2’-deoxyguanosine (6). To a solution of 5

(120 mg, 0.306 mmol) in DMF (3.0 mL) was added CuI (10 mg, 0.061 mmol) and Et_3_N (80 µL) at room temperature. After stirring at room temperature for 5 min under N_2_, the mixture was added 2,2,2-trifluoro-N-(prop-2-yn-1-yl)acetamide (138 mg, 0.918 mmol) and Pd(PPh_3_)_4_ (30.2 mg, 0.031 mmol) at room temperature. The mixture was stirred for 16 hours at room temperature, covered by aluminum foil. Then the solution was quenched by addition of CH_3_OH (2 mL), and then concentrated under reduced pressure. The residue was purified by column chromatography (1:1, Hexanes–EtOAc to 2:2:1, Hexanes–EtOAc–CH_3_OH) to give **6** (103 mg, 81%) as brown foam. ^1^H NMR (400 MHz, CD_3_OD) δ 7.23 (s, 1H, H-8), 6.37 (dd, 1H, *J* = 7.8, 6.1 Hz, H-1’), 4.50–4.44 (m, 1H), 4.31 (s, 2H, C*H_2_*N), 3.98–3.93 (m, 1H), 3.76, (dd, 1H, *J* = 4.2, 11.7 Hz, H-5’a), 3.70, (dd, 1H, *J* = 4.2, 11.7 Hz, H-5’b), 2.50 (ddd, 1H, *J* = 6.3, 7.6, 13.7 Hz, H-2’a), 2.28 (m, 1H, *J* = 2.6, 6.1, 13.7 Hz H-2’b); ^13^C NMR (100 MHz, CD_3_OD) δ 161.43, 158.60 (q, *J* = 37.2 Hz), 154.59, 152.27, 124.77, 117.74 (q, *J* = 290.01 Hz), 101.65, 100.05, 88.83, 85.49, 85.04, 78.27, 77.96, 63.66, 41.4, 31.3; HRMS (ESI) m/z [M] calcd for C_16_H_16_F_3_N_5_O_5_ 415.1104; Found [M + H] ^+^ 416.1211.

#### 5’-*O*-(*tert*-Butyldiphenyl)-7-deaza-7-[3-(2,2,2-trifluoroacetamido)-prop-1-ynyl]-L-2’-deoxyguanosine (7)

To a solution of **6** (103 mg, 0.248 mmol) in DMF (3.0 mL) was added imidazole (25.3 mg, 0.372 mmol) and TBDPSCl (84.2 µL, 0.322 mmol) at 0 °C. The temperature was gradually warmed to room temperature, and then the mixture was stirred at room temperature for another 4 hours. After 4 hours, the mixture was quenched by addition of CH_3_OH (2 mL), and concentrated under reduced pressure. The residue was purified by column chromatography (100% EtOAc to 85:15 EtOAc–MeOH) to obtain **7** (134 mg, 83%) as yellow syrup. ^1^H NMR (400 MHz, CD_3_OD) δ 7.69–7.62 (m, 4H, ArH), 7.44–7.32 (m, 6H, ArH), 7.12 (s, 1H, H-8), 6.40 (app t, 1H, *J* = 6.7 Hz, H-1’), 4.56 (dt, 1H, *J* = 3.8 Hz, *J* = 6.0 Hz, H-3’), 4.25 (s, 2H, C*H_2_*N), 3.97–3.94 (m, 1H, H-4’), 3.85 (dd, 1H, *J* = 3.3, 11.3 Hz, H-5’a), 3.78 (dd, 1H, *J* = 4.1, 11.3 Hz, H-5’b), 2.46–2.37 (m, 1H, H-2’a), 2.31 (ddd, 1H, *J* = 3.8, 6.3, 13.4 Hz, H-2’b), 1.05 (s, 9H, (C*H_3_*)_3_CSi)); ^13^C NMR (100 MHz, CD_3_OD) δ 161.50, 158.51 (q, *J* = 37.7 Hz), 154.65, 152.44, 136.92, 136.82, 134.41, 134.21, 131.14, 131.09, 129.02, 124.08, 117.49 (q, *J* = 286.5 Hz), 101.45, 100.23, 88.39, 85.06, 84.56, 78.26, 77.29, 65.43, 41.63, 31.26, 27.59, 20.20; HRMS (ESI) m/z [M] calcd for C_32_H_34_F_3_N_5_O_5_Si 653.2281; Found [M + H] ^+^ 654.2410.

#### 5’-*O*-(*tert*-Butyldiphenyl)-7-deaza-2-N,N-dimethylamino-7-[3-(2,2,2-trifluoroacetamido)-prop-1-ynyl]-L-2’-deoxyguanosine (8)

To a solution of **7** (70 mg, 0.107 mmol) in acetonitrile (5 mL) was added N, N-dimethylformamide dimethyl acetal (71.2 µL, 0.535 mmol). After stirring at room temperature for 3 hours, the solution was concentrated under reduced pressure. The residue was purified by column chromatography (100% EtOAc to 90:10, EtOAc–CH_3_OH) to give **8** (65 mg, 86%) as yellow syrup. ^1^H NMR (400 MHz, CD_3_OD) δ 8.60 (s, 1H, C*H*=N), 7.69–7.62 (m, 4H, ArH), 7.46–7.31 (m, 6H, ArH), 7.22 (s, 1H, H-8), 6.40 (app t, 1H, *J* = 6.8 Hz, H-1’), 4.60–4.54 (m, 1H, H-3’), 4.26 (s, 2H, C*H_2_*N), 4.01–3.96 (m, 1H, H-4’), 3.86 (dd, 1H, *J* = 3.4, 11.4 Hz, H-5’a), 3.80 (dd, 1H, *J* = 4.1, 11.4 Hz, H-5’b), 3.13 (s, 3H, (C*H_3_*)_2_N), 3.07 (s, 3H, (C*H_3_*)_2_N), 2.50–2.41 (m, 1H, H-2’a), 2.35 (ddd, 1H, *J* = 3.9, 6.3, 13.6 Hz, H-2’b), 1.06 (s, 9H, (C*H_3_*)_3_CSi)); ^13^C NMR (100 MHz, CD_3_OD) δ 162.25, 159.50, 158.53 (q, *J* = 36.9 Hz), 158.18, 151.58, 136.94, 136.83, 134.45, 134.21, 131.17, 131.12, 129.04, 125.12, 117.43, (q, *J* = 287.1 Hz), 104.38, 100.29, 88.51, 85.21, 84.57, 78.21, 77.37, 65.52, 41.68, 41.57, 35.42, 31.27, 27.6, 20.23. HRMS (ESI) m/z [M] calcd for C_35_H_39_F_3_N_6_O_5_Si 708.2703; Found [M + H] ^+^ 709.2829.

#### 5’-*O*-(*tert*-Butyldiphenyl)-7-deaza-2-N,N-dimethylamino-3’*-O-*methylthiomethyl-7-[3-(2,2,2-trifluoroacetamido)-prop-1-ynyl]-L-2’-deoxyguanosine (9)

To a solution of **8** (65 mg, 0.092 mmol) in DMSO (2.0 mL) was added acetic acid (1 mL) followed by acetic anhydride (Ac_2_O, 3 mL). The solution was stirred at room temperature for 18 hours. The mixture was diluted with 10.0 mL EtOAc and quenched at 0 °C by addition of saturated NaHCO_3_. The aqueous layer was washed with EtOAc, and the collected organic layer was dried over Na_2_SO_4_, filtered and concentrated under reduced pressure. The residue was purified by flash column chromatography (100% EtOAc to 9:1 EtOAc–CH_3_OH) to give **9** as yellow syrup (58 mg, 82%). ^1^H NMR (400 MHz, CDCl_3_) δ 10.26 (brs, 1H, NH), 8.61 (s, 1H, C*H*=N), 8.25 (brs, 1H, NH), 7.73–7.63 (m, 4H, ArH), 7.48–7.33 (m, 6H, ArH), 7.12 (s, 1H, H-8), 6.53 (dd, 1H, *J* = 6.1 Hz, *J* = 7.8 Hz, H-1’), 4.74–4.57 (m, 3H), 4.40–4.24 (m, 2H), 4.10 (brs, 1H), 3.85–3.70 (m, 2H), 3.16 (s, 3H, (C*H_3_*)_2_N), 3.08 (s, 3H, (C*H_3_*)_2_N), 2.47–2.29 (m, 2H, H-2’a, H-2’b), 2.15 (s, 3H, SC*H_3_*), 1.10 (s, 9H, (C*H_3_*)_3_CSi). ^13^C NMR (100 MHz, CDCl_3_) δ 160.71, 158.27, 156.95 (q, *J* = 38.0 Hz), 156.26, 150.14, 135.81, 135.72, 132.95, 132.88, 130.16, 130.11, 128.07, 123.39, 115.93, (q, *J* = 288.6 Hz), 103.64, 99.38, 84.73, 83.86, 83.09, 78.27, 76.76, 73.82, 64.25, 41.61, 38.08, 35.38, 31.18, 27.13, 19.42, 14.13; HRMS (ESI) m/z [M]calcd for C_37_H_43_F_3_N_6_O_5_Si 768.2737, found [M + H]^+^ 769.2831.

#### 3’*-O-*azidomethyl-5’-*O*-(*tert*-Butyldiphenyl)-7-deaza-2-N,N-dimethylamino-7-[3-(2,2,2-trifluoroacetamido)-prop-1-ynyl]-L-2’-deoxyguanosine (10)

To a solution of **9** (58.0 mg, 0.076 mmol) in CH_2_Cl_2_ (5.0 mL) was added cyclohexene (400 uL) and 1.0 M sulfuryl chloride in dichloromethane (378 uL, 0.378 mmol) at 0 °C. After stirring for 30 min, the solution was concentrated under reduced pressure at 0 °C, then dried *in vacuo* for 20 min. The residue was re-dissolved in DMF (5.0 mL) at room temperature, and the resulting solution was added NaN_3_ (49 mg, 0.76 mmol). After stirring for 1 hour, the mixture was diluted with EtOAc and washed with saturated NaHCO_3_ solution. The organic layer was dried over Na_2_SO_4_, filtered and concentrated under reduced pressure. The residue was purified by column chromatography (100% EtOAc to 9:1 EtOAc–CH_3_OH) to obtain **10** (41 mg, 71%) as pale-yellow gum. ^1^H NMR (400 MHz, CDCl_3_) δ 9.70 (br, 1H), 8.71–8.57 (m, 2H), 7.72–7.58 (m, 4H, ArH), 7.46–7.32 (m, 6H, ArH), 7.09 (s, 1H, H-8), 6.52 (dd, 1H, *J* = 5.9 Hz, *J* = 8.1 Hz, H-1’), 4.77 (d, 1H, *J* = 9.0 Hz, C*H_2_*N_3_), 4.65 (d, 1H, *J* = 9.0 Hz, C*H_2_*N_3_), 4.53–4.48 (m, 1H), 4.38–4.22 (m, 2H, C*H_2_*N), 4.14–4.10 (m, 1H), 3.82 (dd, 1H, *J* = 3.7, 11.2 Hz, H-5’a), 3.77 (dd, 1H, *J* = 5.0, 11.2 Hz, H-5’b), 3.17 (s, 3H, C*H_3_*N), 3.09 (s, 3H, C*H_3_*N), 2.49 (ddd, 1H, *J* = 2.5, 5.9, 13.7 Hz, H-2’a), 2.37 (m, 1H, H-2’b), 1.10 (s, 9H, (C*H_3_*)_3_CSi). ^13^C NMR (100 MHz, CDCl_3_) δ 158.36, 157.09 (q, *J* = 37.8 Hz), 156.22, 150.03, 135.83, 135.74, 132.89, 132.84, 130.26, 130.21, 128.13, 123.13, 115.92 (q, *J* = 286.6 Hz), 99.59, 84.81, 84.01, 83.01, 80.88, 79.21, 78.18, 77.43, 64.07, 41.67, 38.54, 35.44, 31.20, 29.91, 27.16, 19.48; HRMS (ESI) m/z [M] calcd for C_36_H_40_F_3_N_9_O_5_Si 763.2874, found [M + H]^+^ 764.3012.

#### 3’*-O-*azidomethyl-7-deaza-2-N,N-dimethylamino-7-[3-(2,2,2-trifluoroacetamido)-prop-1-ynyl]-L-2’-deoxyguanosine (11)

To a solution of **10** (41 mg, 0.053 mmol) in THF (1.0 mL) was added TBAF in THF (1.0 M, 0.1 mL) at room temperature. After stirring for 20 min at room temperature, the mixture was concentrated, then washed with EtOAc and saturated aqueous NaHCO_3._ The organic layer was dried over Na_2_SO_4_, filtered and concentrated under reduced pressure. The residue was purified by column chromatography to give **11** as a yellow foam (22 mg, 78%). ^1^H NMR (CD_3_OD) δ 8.64 (s, 1H, NH), 7.35 (s, 1H, H-8), 6.47 (dd, *J* = 6.1 Hz, *J* = 8.1 Hz, 1H, H-1’), 4.84–4.77 (m, 2H, CH_2_N_3_), 4.52 (app dt, *J* = 2.3 Hz, *J* = 6.1 Hz, 1H, H-3’), 4.30 (s. 2H, C*H_2_*N), 4.12-4.07 (m, 1H, H-4’), 3.77–3.68 (m, 2H, H-5’a, H-5’b), 3.19 (s. 3H, (C*H_3_*)_2_CH=N), 3.10 (s. 3H, (C*H_3_*)_2_CH=N), 2.64-2.55 (m. 1H, H-2’a), 2.48 (ddd, 1H, *J* = 2.3, 6.1, 13.7 Hz, H-2’b); ^13^C NMR (100 MHz, CD_3_OD) δ 158.60, 157.33, 150.62, 124.63, 103.55, 99.29, 85.7,0, 84.28, 84.17, 82.07, 79.68, 77.20, 62.45, 40.53, 38.21, 34.42, 30.29, 29.90; HRMS (ESI) m/z [M] calcd for C_20_H_22_F_3_N_9_O_5_ 525.1696, found [M + H]^+^ 526.1797.

#### 3’-*O*-azidomethyl-7-deaza-7-[3-amino-prop-1-ynyl]-L-2’-deoxyguanosine 5’-*O*-triphosphate (12)

The starting material compound **11** (22 mg, 0.042mmol), together with sponge (16.0 mg, 0.084 mmol) was dried for 72 hours *in vacuo*. To a solution of starting materials in trimethyl phosphate (0.5 mL) was injected phosphoryl oxychloride (10.5 µL, 0.105 mmol) at 0 °C. The mixture was stirred at 0 °C for 2 hours and monitored by HPLC, until HPLC indicated **11** was completely consumed. Then a well-vortexed mixture of tributylammonium pyrophosphate (93 mg, 0.17 mmol) and tributylamine (80.8 µL, 0.39 mmol) in an anhydrous DMF (1.0 mL) was injected to the above solution. The mixture was stirred for another 1.5 hours at room temperature. After which, triethylammonium bicarbonate buffer (0.1 M TEAB buffer, 5.0 mL, pH 8.0) was added and stirred for another hour at room temperature, then concentrated NH_4_OH (10.0 mL) was added and stirred overnight at room temperature. The resulting mixture was concentrated, and the residue was diluted with HPLC water (10.0 mL). The mixture was extracted with CH_2_Cl_2_ (3 x 10.0 mL) and the aqueous layer was concentrated under reduced pressure to 5.0 mL. The residue was subjected to preparative C18 HPLC (0%B to 90%B over 50 min, mobile phase A: 0.1M TEAB, B: ACN). The fractions with products were collected and concentrated under reduced pressure. The residue was purified with semi-prep HPLC column on an ion-exchange column (Agilent PL-SAX, 10 µm, 1000Å), mobile phase: A, 15% acetonitrile in water; B, 15% acetonitrile in 1M TEAB buffer. Elution was performed in gradient condition (0%B to 80%B over 50 min). The fractions with products were collected, and concentrated under reduced pressure, and then washed 3 times with HPLC grade water (5 mL). After NMR analysis, the pure product was lyophilized to give **12** as a pale-yellow powder (5.5 mg, 21%). ^1^H NMR (D_2_O) δ 7.55 (s, 1H, H-8), 6.41 (dd, *J* = 2.1 Hz, *J* = 6.3 Hz, 1H, H-1’), 4.96–4.84 (m, 2H, CH_2_N_3_), 4.73 (brs, 1H, H-3’), 4.40 (s, 1H, H-4’), 4.21–4.17 (m, 2H, H-5’), 3.98 (brs, 2H, CH_2_N), 2.73–2.54 (m. 2H, H-2’). ^31^P NMR (D_2_O) δ –10.50 (d, *J* =19.6 Hz, 1P, Pγ), –11.40 (m, *J* =19.6 Hz, 1P, Pα.), –23.14 (t, *J* =19.4 Hz, 1P, P_β_). HRMS (ESI) calcd for [M] C_15_H_21_N_8_O_13_P_3_ 614.0441, found 613.1009 (M-H)^-^.

#### 3’*-O-*azidomethyl-7-deaza- 7-[3-acetamido-(2-(2-(2-(3-((2-aminoethylcarbamoyl)phenoxy)-1-adizoethoxy)ethoxy)-prop-1-ynyl]-L-2’-deoxyguanosine 5’-*O*-triphosphate (13)

To a solution of **12** (5.0 mg, 0.0081 mmol) in 0.1 M NaHCO_3_/Na_2_CO_3_ solution (20 uL) was added the solution of **S-3** (12 mg, 0.0162 mmol) in ACN (20 uL). The reaction was stirred at room temperature and monitored by HPLC until the completion of **12.** After reaction completion in one hour, the reaction mixture was added ammonia (40 uL) and then stirred at room temperature for 30 min. The crude was purified on preparative HPLC (Polaris C18, 10.0 x 250 mm, 180Å, 5 mm) with gradient 0%B to 90%B over 50 mins (A: 0.1M TEAB, B: ACN). The fractions were collected and dried under reduced pressure. The residue was washed with HPLC grade water 3 times (5 mL) and lyophilized to give **13** as white powder (5.2 mg, 66%).^1^H NMR (D_2_O) δ 7.44–7.35 (m, 2H), 7.30–7.25 (m, 1H), 7.21–7.17 (m, 1H), 7.09 (s, 1H, H-8), 6.96–6.70 (m 1H), 6.19–6.10 (m, 1H, H-1’), 5.15 (dd, *J* = 4.6 Hz, *J* = 10.2 Hz, 1H), 4.95 (dd, *J* = 4.6Hz, *J* = 10.2 Hz, 1H), 4.86 (dd, *J* = 4.6 Hz, *J* = 10.2 Hz, 1H), 4.60–4.55 (m, 1H), 4.35 (br, 1H), 4.30–4.10 (m, 10H), 4.07 (s, 1H), 4.00–3.80 (m, 4H), 3.71–3.59 (m, 3H), 2.61– 2.40 (m, 2H). ^31^P NMR (D_2_O) δ –9.00 (m, 1 P. P*γ*), –11.32 (d, *J* = 18.98 Hz, 1 P, P_α_.), –22.56 (t, *J* = 19.06 Hz, 1 P, P_β_). HRMS (ESI): m/z [M] calcd for C_30_H_40_N_13_O_18_P_3_ 963.1827; Found [M – H]^−^ 962.1768.

### NMR measurement

All deuterated solvents were purchased commercially, and used without further purification unless specified. Proton nuclear magnetic resonance (^1^H NMR) spectra were recorded on a Bruker Ascend^TM^ (400 MHz) spectrometer at Chapman University (Irvine, CA) and reported in parts per million (ppm) from a CDCl_3_ (7.26 ppm), CD_3_OD (4.78 ppm) or D_2_O. Data were reported as follows: (s = singlet, d = doublet, t = triplet, q = quartet, m = multiplet, dd = doublet of doublets, *J* = coupling constant in Hz, integration). Carbon-13 nuclear magnetic resonance (^13^C NMR) was recorded on a Bruker Ascend^TM^ (100 MHz) spectrometer at Chapman University and reported in parts per million (ppm) from a CDCl_3_ (77.23 ppm), CD_3_OD (48.15 ppm) or D_2_O. Proton-decoupled ^31^P NMR spectra were recorded on a Bruker Ascend^TM^(121.4 MHz) spectrometer at Chapman University. Data were reported as follows: (d = doublet, t = triplet, m = multiplet, *J* = coupling constant in Hz, integration).

### Purification of synthetic triphosphate 1, 12, 13

All solvents were purchased commercially, and used without further purification unless specified. Water (HPLC grade) was purchased from Lab Alley (Austin, TX), acetonitrile (HPLC grade) was purchased from Oakwood Chemicals (Estill, SC) and triethylammonium bicarbonate buffer (TEAB, 1M, pH 8.5) was purchased from Boston Bioproducts. Compound **12** was purified with reverse-phase HPLC on a C18 reversed phase column (Polaris 5 C18-A, 180Å, 10.0 x 250 mm, 5 µm), mobile phase: A, 100 mM TEAB buffer in water; B, acetonitrile. Elution was performed in gradient condition (0%B to 90%B over 50 min). Fractions were collected and evaporated under reduced pressure. The resulting residue was washed with HPLC grade water (5 mL x 3) three times under reduced pressure, then was dried *in vacuo*. The dried residue was resuspended in HPLC grade water (3 mL), and purified with semi-prep HPLC column on an ion-exchange column (Agilent PL-SAX, 10 µm, 1000Å), mobile phase: A, 15% acetonitrile in water; B, 15% acetonitrile in 1M TEAB buffer. Elution was performed in gradient condition (0%B to 80%B over 50 min). Fractions were collected and evaporated under reduced pressure. The resulting residue was washed with HPLC grade water (5 mL x 3) for three times under reduced pressure, then was dried via lyophilization. Compound **1** and **13** were purified with reverse-phase HPLC on a C18 reversed phase column (Polaris 5 C18-A, 180Å, 10.0 x 250 mm, 5 µm), mobile phase: A, 100 mM TEAB buffer in water; B, acetonitrile. Elution was performed in gradient condition (0%B to 90%B over 50 min). Fractions were collected and evaporated under reduced pressure. The resulting residue was washed with HPLC grade water (5 mL x 3) for three times under reduced pressure, then was dried via lyophilization. All purifications were performed on Shimadzu Nexera prep system with SPD-M40 photodiode array detector and LC-20AR semi-prep HPLC pump.

### Fmoc-based solid phase peptide synthesis

All solvents and chemicals were purchased commercially, and used without further purification unless specified. All Fmoc-D-amino acids were purchased from Chempep (Wellington, FL). Piperidine was purchased from Spectrum Chemical (New Brunswick, NJ). N,N-diisopropylcarbodiimide (DIC), N,N-diisopropylethylamine (DIPEA), thioanisole, triisopropylsilane, and 1,2-ethanedithiol were purchased from Sigma-Aldrich. Oxyma and HO-TCP(Cl) Protide resin were purchased from CEM (Matthews, NC). N,N-dimethylforamide and diethyl ether were purchased from Carolina Chemical (Charlotte, NC). Trifluoroacetic acid (TFA) was purchased from Fisher Scientific. Fmoc-D-Lys(Boc)-Wang Resin was purchased from Aapptec (Louisville, KY). All peptides were synthesized by Fmoc-based SPPS on a Liberty Blue 2.0 automatic microwave peptide synthesizer (CEM). Peptides with a carboxylate group at C-terminal, such as D-9°N-N-9 and D-9°N-C-6, were synthesized on a preloaded Fmoc-D-amino acid Wang resin (Aapptec), which has preloaded first C-terminal residue. All other peptides with hydrazine at C-terminal were synthesized on a Fmoc-hydrazine 2-chlorotrityl resin to prepare peptide hydrazides. The Fmoc-hydrazine 2-chlorotrityl resin was prepared by a two-step synthesis, chlorination as suggested by the manufacturer on HO-TCP(Cl)-Protide resin (CEM) and Fmoc-hydrazine attachment^50^. All swelling, deprotections and couplings were performed on a Liberty Blue 2.0 automatic microwave peptide synthesizer (CEM). All resins were swelled in DMF for 30 min before the SPPS began. The Fmoc group of both resins and the assembled amino acids were removed by treatment with 10% piperidine supplemented with 0.1M Oxyma in DMF at 90 °C for 1 min. Coupling of amino acids except Fmoc-D-Cys(Trt)-OH, Fmoc-D-Thz-OH, Fmoc-D-His(Trt)-OH, Fmoc-D-Cys(Acm)-OH and Fmoc-D-Arg(Pbf)-OH were executed with 4 equiv. of amino acid, 4 equiv. of Oxyma (with 0.1M DIPEA) and 8 equiv. of DIC. Fmoc-D-Cys(Trt)-OH, Fmoc-D-Thz-OH, Fmoc-D-His(Trt)-OH, Fmoc-D-Cys(Acm)-OH and Fmoc-D-Arg(Pbf)-OH were carried out at 50 °C for 10 min to avoid side reactions at high temperature. After the completion of peptide chain assembly, peptides were cleaved from resin using H_2_O/triisopropylsilane/1,2-ethanedithiol//thioanisole/trifluoroacetic acid (0.5/0.5/0.5/0.25/8.25) (vol/vol). The cleavage reaction was monitored at specific time intervals by HPLC and LC-MS, and the reaction completion was observed at 4 hours at ambient temperature. After cleavage reaction, the solution was poured into conical tubes with cold diethyl ether to precipitate the crude peptides. After centrifugation, the supernatant was discarded and the precipitates were washed with cold diethyl ether twice. The crude peptides were dissolved in H_2_O/CH_3_CN with 0.1% TFA for analysis on analytical RP-HPLC. The main product peak collected from analytical RP-HPLC was analyzed on LC-MS. The crude peptides were purified by preparative RP-HPLC, and re-analyzed the purity on analytical RP-HPLC and LC-MS.

### Native chemical ligations

All chemicals were purchased commercially, and used without further purification unless specified. Guanidine hydrochloride (Gn•HCl), sodium phosphate dibasic heptahydrate (Na_2_HPO_4_•7H_2_O) and sodium phosphate monobasic dihydrate (NaH_2_PO_4_•2H_2_O) were purchased from Sigma-Aldrich. 4-Mercaptophenylacetic acid was purchased from Ambeed. Acetylacetone (acac) was purchased from Fisher Scientific. Tris-(2-carboxyethyl)phosphine hydrochloride (TCEP) was purchased Tokyo Chemical Industry. To a solution of C-terminal peptide hydrazide segment (peptide 1) in 6M Gn•HCl was added 50 equiv. MPAA and 2.5 equiv. acetylacetone. The reaction mixture was agitated at ambient temperature and monitored on analytical RP-HPLC. After completion of activation, the solution of N-terminal cysteine-peptide (peptide 2) in buffer (6M Gn•HCl, 200 mM Na_2_HPO_4_, pH 8.5) was added into the above solution with equal volume, then adjusted pH to 6.5. The reaction mixture was agitated at ambient temperature overnight and monitored on analytical RP-HPLC. After completion of NCL reaction, the NCL reaction was added 100 mM TCEP in buffer (6M Gn•HCl, 100 mM Na_2_HPO_4_, pH 7.0). The reaction was kept at room temperature for 30 min with stirring. Finally, the ligation product was purified by preparative HPLC and analyzed by analytical HPLC and LC-MS.

### Desulfurization

All chemicals were purchased commercially, and used without further purification unless specified. Guanidine hydrochloride (Gn•HCl), sodium phosphate dibasic heptahydrate (Na_2_HPO_4_•7H_2_O), sodium phosphate monobasic dihydrate (NaH_2_PO_4_•2H_2_O), and reduced L-glutathione (reduced L-GSH) were purchased from Sigma-Aldrich. Tris-(2-carboxyethyl)phosphine hydrochloride (TCEP) and 2,2-azobis[2-(2-imidazolin-2-yl)propane] dihydrochloride (VA-044) were purchased Tokyo Chemical Industry. The Cysteine-containing peptide (2 mg/mL) was dissolved in desulfurization buffer (6M Gn•HCl, 100 mM Na_2_HPO_4_, 200 mM TCEP, 40 mM reduced L-GSH and 20 mM VA-044, pH 7.0). The mixture was stirred at 37 °C overnight with covered aluminum foil, and the desulfurized product was analyzed by HPLC and purified by preparative HPLC. The collected fraction from preparative HPLC was analyzed on analytical HPLC and LC-MS.

### Acm deprotection

All chemicals were purchased commercially, and used without further purification unless specified. Guanidine hydrochloride (Gn•HCl), sodium phosphate dibasic heptahydrate (Na_2_HPO_4_•7H_2_O), sodium phosphate monobasic dihydrate (NaH_2_PO_4_•2H_2_O), Palladium(II) chloride (PdCl_2_) and dithiothreitol (DTT) were purchased from Sigma-Aldrich. Tris-(2-carboxyethyl)phosphine hydrochloride (TCEP) was purchased Tokyo Chemical Industry. The Acm-protected cysteine-containing peptide (0.5–1.0 mM) was dissolved in Acm deprotection buffer (6M Gn•HCl, 100 mM Na_2_HPO_4_ and 40 mM TCEP, pH 7.0). The solution of PdCl_2_ in 6M Gn•HCl (1 mg PdCl_2_/20 uL) was added to the above solution to a final concentration of 3 mg peptide/1 mg PdCl_2_. The reaction mixture was incubated with stirring at 30 °C until reaction completion by determination on analytical HPLC and ESI-MS. A solution of 2M DTT in 6M Gn•HCl was added to the above Acm deprotection solution to quench the reaction. The reaction mixture was stirred at 30 °C for 1 hour and purified by preparative HPLC. The collected fraction was analyzed on analytical HPLC and LC-MS.

### Analytical RP-HPLC

All solvents were purchased commercially, and used without further purification unless specified. Water with 0.1% TFA (HPLC grade) and acetonitrile with 0.1% TFA (HPLC grade) were purchased from Fisher Scientific. All peptides were analyzed on a C4 reversed phase column (Welch C4, 4.6 x 250 mm, 5 µm, 300Å) with elution gradient (20%B to 70%B over 30 min), mobile phase: A, 0.1%TFA in water; B, 0.1%TFA in acetonitrile. The analysis of peptides was performed on a Shimadzu Nexara analytical system with SPD-M40 photodiode array detector and LC-40D solvent delivery module.

### Preparative RP-HPLC

All solvents were purchased commercially, and used without further purification unless specified. Water with 0.1% TFA (HPLC grade) and acetonitrile with 0.1% TFA (HPLC grade) were purchased from Fisher Scientific. All crude peptides were dissolved in H_2_O/CH_3_CN with 0.1% TFA and purified on a C18 reversed phase column (Polaris C18-A, 21.2 x 250 mm, 5 µm, 180Å) with varies elution gradient (indicated on each peptide in SI) on Waters Delta 4000, mobile phase: A, 0.1%TFA in water; B, 0.1%TFA in acetonitrile. The NCL peptides, desulfurization peptides and Acm deprotection peptides were diluted in H_2_O/CH_3_CN with 0.1% TFA and purified on a C4 reversed phase column (Welch C4, 21.2 x 250 mm, 5 µm, 300Å) with varies elution gradient (indicated on each peptide in SI) on Shimadzu Nexara prep system with SPD-M40 photodiode array detector and LC-20AR semi-prep HPLC pump, mobile phase: A, 0.1%TFA in water; B, 0.1%TFA in acetonitrile.

### MALDI-TOF MS and LC-MS

All solvents and chemicals were purchased commercially, and used without further purification unless specified. 3-hydropicolinic acid and diammonium citrate were purchased from Sigma-Aldrich. Water with 0.1% FA (LCMS grade) and acetonitrile with 0.1% FA (LCMS grade) were purchased from Fisher Scientific. All purified peptides were analyzed on Agilent 6545XT AdvanceBio LC/QTOF with Agilent 1260 HPLC system. Elution was performed in gradient condition (30%B to 95%B over 5 min), mobile phase: A, 0.1%FA in water; B, 0.1%FA in acetonitrile. All purified synthetic compounds **1**-**13** were analyzed on Bruker Impact II^TM^ Ultra-High Resolution Q-TOF at Chapman University. Elution was performed in gradient conditions (30%B to 95%B over 5 min), mobile phase: A, 0.1%FA in water; B, 0.1%FA in acetonitrile. All purified oligonucleotides were analyzed on Autoflex Speed MALDI-TOF mass spectrometer at Chapman University. The purified oligonucleotides were dissolved in water (500 µL), then added DAC solution (500 µL, 25 mg/mL in water) and matrix solution (1 mL, 60 mg/mL in 50% ACN/water). The above oligonucleotide solution was spotted on target plate and then was dried with air.

### Refolding, Heat Shock and Storage

Mass confirmed N- and C-terminal of the 9°N polymerase was dissolved in 5 M guanidine hydrochloride (GnHCl) and 10 mM BME at 5 mg/ml concentrations. Concentration of the N- and C-terminal were calculated to ∼1 μM (22.4 μL and 17.5 μL, respectively) for a final volume of 1 mL in 5 M GnHCl with 10 mM BME and incubated at room temperature for 4 hours. After incubation, dialysis samples are placed into Thermo Fisher 10 kDa Slide-A-Lyzer and placed into dialysis buffer 1 (40 mM Tris-HCl at pH 7.5, 100 mM KCl, 0.1 mM EDTA, 10% glycerol) at 4°C for 15-16 hours with stirring. After overnight incubation, the dialysis cassette was transferred to dialysis buffer 1 and furthered dialyzed at 4°C for 4 hours with stirring. Dialyzed solution was removed from dialysis cassette and placed into a fresh 2 mL centrifuge tube, precipitation was observed, then samples were centrifuge at 20,000 x g for 10 minutes at room temperature. Dialysis supernatant was subjected to an 85°C heat shock for 15 minutes and placed at 4°C for at least 30 minutes. Samples were centrifuged at 20,000 x g for 20 minutes at room temperature. Supernatant was removed and further concentrated in 10 kDa Amicon concentrator to 50 μL in buffer 1 with the addition of 80% glycerol (50 μL), for a final buffer 1 with 50% glycerol and stored at -20°C.

### Single Base Primer Extension

Primer extension reaction (20 μL) was set up as following: Oligonucleotide G template (5’-GATCGCGCCGCGCCTTGGCGCGGCGC-3’, 2 to 20 pmol), and NEB ThermoPol buffer (1x) were mixed and melted at 90°C for 2 minutes and annealed for 5 minutes at room temperature. Followed by the addition of 200 pmol of (L)-3’-O-azidomethyl-dGTP, 4 mM MnCl_2_, and 6 µL of a 5x (D)-9°N polymerase stock to initiate the reaction. Extensions were performed as follows: 65°C for 2 minutes, followed by 25 to 100 cycles of 37°C for 30 seconds and 65°C for 2 minutes. Reactions were monitored at various time points by removing 2 μL of the reaction, mixed with 10 mM EDTA and 2x TBE Urea Sample loading buffer and melted at 90°C for 5 minutes. Samples were loaded onto Novex 15% TBE-Urea page at 180 V for 70 min. Gels were stained with 1x SYBR Gold (Invitrogen) in 1x TBE buffer for 5 minutes and imaged under blue light.

### Purification of extended primers

All solvents were purchased commercially and used without further purification unless specified. Triethylammonium acetate (1M, pH 7.0) was purchased from Sigma-Aldrich. Water (HPLC grade) was purchased from was purchased from Lab Alley. Acetonitrile (HPLC grade) was purchased from Oakwood Chemicals. The oligonucleotides in extension experiments were purified on an Agilent PLRP-S column (1000Å, 5 µm), mobile phase: A, 100 mM TEAA buffer in water; B, acetonitrile/water (7:1). Elution was performed in gadient condition (0%B to 50%B over 25 min) for extension and in gradient conditions (0%B to 25%B over 25 min) for TCEP cleavage. Fractions were collected based on the absorbance at 260 nm and 647 nm. All purifications were performed on a Shimadzu Nexara analytical system with SPD-M40 photodiode array detector and LC-40D solvent delivery module. The collected solution was dried on a lyophilzer (Labconco Freezone 4.5L, –105°C), and analyzed by MALDI-TOF MS.

### Protease Stability

Trypsin protease was purchased from Sigma Aldrich and Proteinase K was purchased from Thermo Fisher Scientific. Trypsin protease was dissolved in 1 mM HCl at a 10 mg/ml stock and further diluted to a 1 mg/ml work stock concentration. Proteinase K was diluted from a 20 mg/ml stock to 1 mg/ml working stock in 20 mM Tris-HCl, pH 7.5 and 1 mM CaCl_2_. ∼1 μg of recombinant candidate 1 and ∼1 μg of mixed N20 and C13, was suspended in Proteinase K buffer and Trypsin buffer (50 mM Tris-HCl, pH 8.0 and 20 mM CaCl_2_) with the addition of 2 μg of each protease in a total volume of 20 μL. All samples were incubated at 37°C for 3 hr. Proteases were heat denatured at 95°C for 5 minutes. Samples were mixed with 4 μL of 6x Laemmli SDS sample buffer (Thermo Fisher), heated at 85°C for 2 minutes and loaded onto a Novex 12% Tris-Glycine Wedge Well Gels and ran at 225 V for 35 minutes. Protein gel was stained with Pierce Silver Stain Kit (Thermo Scientific) performed using manufacturer protocol. Protein gel was imaged under a white light imager and then grey scaled.

### Nuclease Test

Nucleases were purchased from NEB: DNase I, Micrococcal, and T5 Exonuclease. For each reaction 0.5 μM concentration of either (D)-Oligonucleotide G or (L)-Oligonucleatiode G were dissolved in 1x buffer solution for each individual reaction in a final volume of 50 μL. Amounts of nuclease added were DNase I 2 units (1 μL), micrococcal 3125 gel units (0.25 uL), and T5 Exonuclease 2 units (1 μL). DNase I was incubated for 10 minutes at 37°C, quenched with 5 mM EDTA and heat inactivated at 75°C for 10 minutes. Micrococcal and T5 Exonuclease were incubated for 30 minutes at 37°C. Aliquots of 2 μL were removed and mixed with 2x TBE Urea Sample loading buffer and melted at 90°C for 5 minutes. Samples were loaded onto Novex 15% TBE-Urea page at 180 V for 70 min. Gels were stained with 1x SYBR Gold (Invitrogen) in 1x TBE buffer for 5 minutes and imaged under blue light.

## Data Availability

All data generated or analyzed during this study are included in the main text and supplemental information. Data supporting the findings of this study are also available from the corresponding author upon request. Source data are provided with this paper.

### Acknowledgement

We thank George Church (Harvard) and Jingyue Ju (Columbia) for helpful discussions. We thank Chapman University School of Pharmacy NMR Core Facility for the assistance and the usage of NMR, LC-MS and MALDI-TOF MS instruments. We also thank Analytical Chemistry Instrumentation Facility, University of California, Riverside, for the assistance and the usage of the LC-MS instrument.

## Author Contributions

D.H.K. conceived the project and provided overall guidance on experimental designs and directions. B.-S.H., G.Z., J.S.L designed and performed experiments for chemical synthesis. B.-S.H., G.Z., J.S.L analyzed and characterized all chemically synthetic compounds. B.-S.H, E.J.Y., R.G., J.R.B., P.J. designed and performed experiments for chemical synthesis of peptides and proteins including SPPS and NCL. B.-S.H, E.J.Y., R.G., J.R.B., P.J. analyzed and characterized all peptides and proteins. E.J.Y., B.-S.H., J.R.B., E.M. performed the experiments for synthetic protein refolding, primer extension and stability tests. E.J.Y., B.-S.H., J.R.B., E.M. analyzed and characterized protein activity and extension products. D.H.K. supervised this project. B.-S.H., E.J.Y., D.H.K. wrote paper and drafted figures. All authors revised and approved the final paper.

