## Supplemental Information for "Design and synthesis of mirror image fluorescent nucleotide reversible terminator, (L)-3ʹ-O-azidomethyl-dGTP-N_3_-fluorophore and (D)-9°N mutant DNA polymerase for mirror image (L)-DNA sequencing by synthesis"

Supporting Information

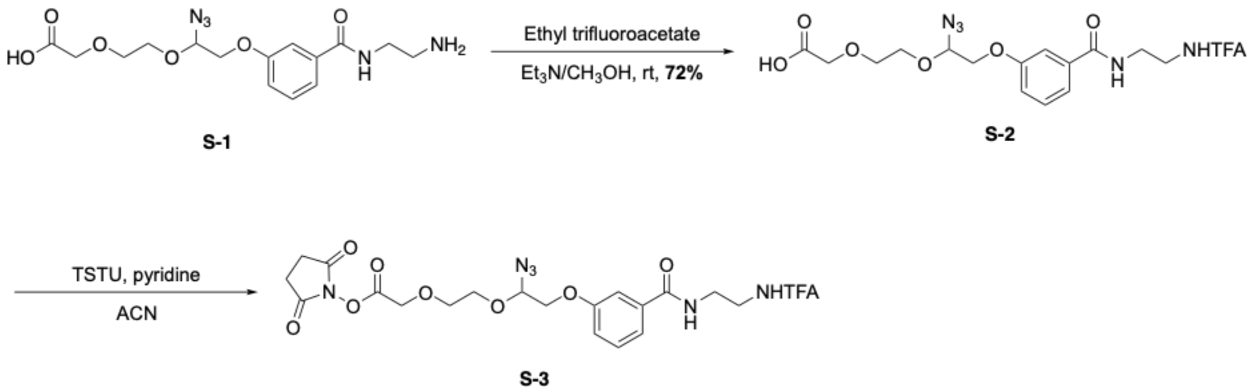

**Supplementary Scheme 1.** Synthesis of general cleavable linker NHS ester (**S-3**)

**2-(2-(2-(3-((2-trifluoroacetamidoethyl)carbamoyl)phenoxy)-1-azidoethoxy)ethoxy)acetic acid** (**S-2**). To a solution of **S-1**[1] (50.0 mg, 0.13 mmol) in MeOH (5.0 mL) was added triethylamine (0.5 mL), and then slowly added ethyl trifluoroacetate (15.6 μL, 0.14 mmol). The reaction was monitored by HPLC. After complete consumption of **S-1**, the solution was concentrated under reduced pressure and the residue was purified by column chromatography (90:10, $CH_2Cl_2$–$CH_3OH$) to obtain **S-2** as white powder (45.2 mg, 72%). $^1$HNMR (400 MHz, $D_2O$) δ 7.38–7.30 (m, 3H), 7.10–7.07 (m, 1H), 4.97 (dd, $J$ = 0.6 Hz, $J$ = 3.8 Hz, 1H), 4.56 (brs, 2H), 4.20 (dd, $J$ = 4.6 Hz, $J$ = 10.2 Hz, 1H), 4.08 (dd, $J$ = 4.6Hz, $J$ = 10.2 Hz, 1H), 4.02–3.91 (m, 1H), 3.90 (s, 2H) 3.84–3.79 (m, 1H), 3.70–3.63 (m, 2H), 3.51–3.44 (m, 4H). HRMS (ESI) M: $C_{17}H_{20}F_3N_5O_7$, calc. 463.1315; Found $[M + Na]^+$ 486.1203.

**2-(2-(2-(3-((2-trifluoroacetamidoethyl)carbamoyl)phenoxy)-1-azidoethoxy)ethoxy)acetic acid NHS ester** (**S-3**). To a solution of **S-2** (25.0 mg, 0.054 mmol) in ACN (3.0 mL) was added pyridine (0.5mL) and TSTU (16.5 mg, 0.056 mmol), and the resulting mixture was stirred at room temperature for 2 hours, and monitored by HPLC. After complete consumption of **S-2**, the reaction mixture was concentrated, and then filtered through column chromatography (97:3, $CH_2Cl_2$–$CH_3OH$) to give **S-3**. The elutions were dried and directly used for the next step without taking $^1$H NMR due to the instability of **S-3**. HRMS (ESI) M: $C_{21}H_{23}F_3N_6O_9$, calc. 560.1479; Found $[M + Na]^+$ 583.1381.

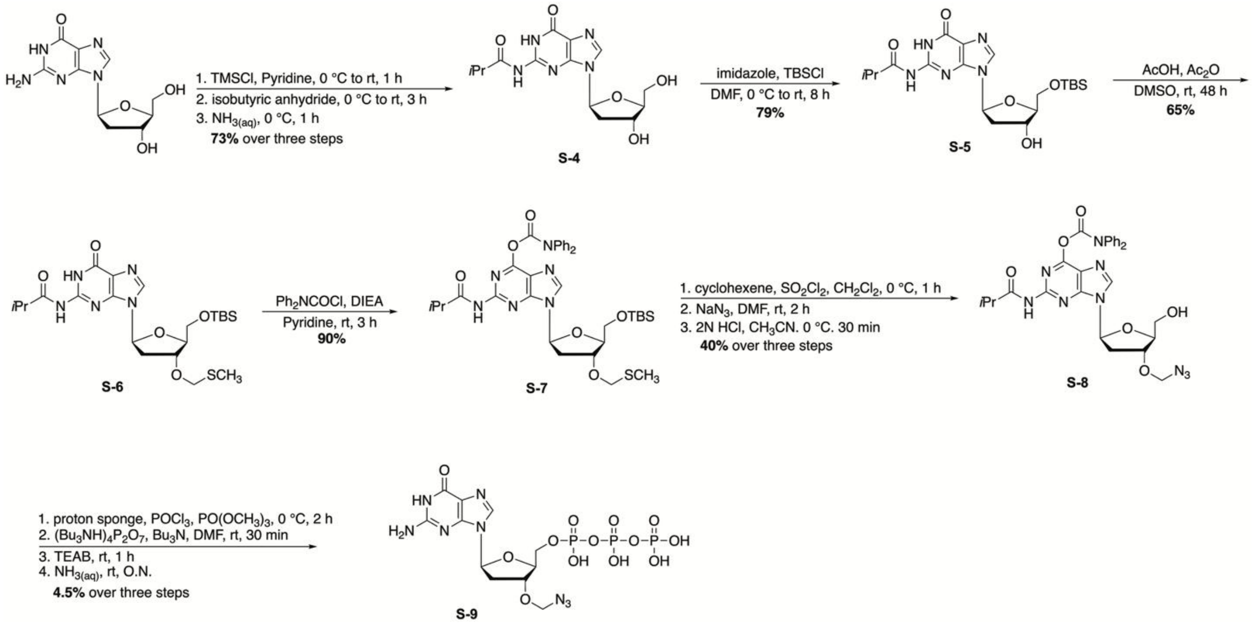

**$N^2$-Isobutyryl-L-2'-deoxyguanosine (S-4).** β-L-deoxyguanosine (2 g, 7.48 mmol) was co-evaporated (2 x 12 mL), and then added pyridine (30 mL) and cool the suspension to 0 °C. Trimethylsilyl chloride (4.8 mL, 37.4 mmol) was added drop wisely via syringe. The ice bath was then removed and the mixture was stirred for 1 hour. The solution was cooled to 0 °C and isobutyric anhydride (6.2 mL, 37.4 mmol) was added drop wisely via a syringe. The ice bath was removed and the resulting solution was stirred at room temperature for 3 hours. After stirred for 3 hours, the mixture was cooled to 0 °C and ice cold water (5 mL) was slowly added and stirred for 15 min, followed by concentrated ammonia solution (10 mL) to a final 2.5 M concentration of ammonia. The mixture was stirred on ice bath for 1 hour and then evaporated to dryness. The residue was redissolved in $CH_3OH$ and was added silica gel and concentrated until dryness. The mixture was concentrated until dryness. The resulting residue was purified by column chromatography (19:1 to 9:1, $CH_2Cl_2$–$CH_3OH$) to give **S-4** (1.85 g, 73%) as a white solid. $^1H$ NMR (400 MHz, $CD_3OD$) δ 8.24 (s, 1H, H-8), 6.36 (app t, 1H, $J_{1',2'a} = J_{1',2'b} = 6.8$ Hz, H-1'), 4.53 (app dt, 1H, $J = 6.0$ Hz, $J = 3.5$ Hz, H-3'), 4.01–3.96 (m, 1H, H-4'), 3.78 (dd, 1H, $J = 3.8$, 12.0 Hz, H-5'a), 3.72 (dd, 1H, $J = 4.3$, 12.0 Hz, H-5'b), 2.78–2.61 (m, 2H, $COC\mathit{H}(CH_3)_2$, H-2'a), 2.42 (ddd, 1H, J = 3.5, 6.3, 13.5 Hz, H-2'b), 1.23 (d, 6H, $J = 7.0$ Hz, $COCH(C\mathit{H}_3)_2$); $^{13}C$ NMR (100 MHz, $CD_3OD$) δ 180.89, 149.41, 149.00, 138.68, 120.43, 88.43, 84.65, 71.51, 62.16, 40.91, 36.09, 18.47. HRMS (ESI) m/z [M] calcd for $C_{14}H1N_5O_5$ 337.1386; Found $[M + H]^+$ 338.1476.

**5'-O-*tert*-butyldimethylsilyl-N$^2$-Isobutyryl-L-2'-deoxyguanosine (S-5).** To a solution of **S-4** (1.8 g, 5.33 mmol) in anhydrous DMF (30 mL) was added imidazole (544 mg, 8.0 mmol) and *tert*-butyldimethylsilyl chloride (885 mg, 5.87 mmol) at 0 ºC under nitrogen, and warmed to room temperature. After stirred for 8 h at room temperature, the solution was added iced cold water and extracted with EtOAc (3 x 100 mL). The combined organic layers were dried over anhydrous $Na_2SO_4$, filtered and concentrated. The resulting residue was purified by column chromatography (19:1 to 9:1, $CH_2Cl_2$–$CH_3OH$) to afford compound **S-5** (1.9 g, 79%) as a white foam. $^1$H NMR (400 MHz, $CD_3OD$) δ 8.07 (s, 1H, H-8), 6.32 (app t, 1H, $J_{1',2'a} = J_{1',2'b}$ = 6.3 Hz, H-1'), 4.53 (app dt, 1H, *J* = 6.3, 3.9 Hz, H-3'), 4.02–3.98 (m, 1H, H-4'), 3.88 (dd, 1H, *J* = 3.4, 11.4 Hz, H-5'a), 3.86 (dd, 1H, *J* = 3.4, 11.4 Hz, H-5'b), 2.72 (sep, 1H, *J* = 7.0 Hz, COC*H*($CH_3$)$_2$), 2.60 (app dt, 1H, *J* = 6.3, 13.3 Hz, H-2'a), 2.46 (ddd, 1H, *J* = 3.9, 6.3, 13.3 Hz, H-2'b), 1.22 (d, 6H, *J* = 7.0 Hz, COCH(C*H*$_3$)$_2$), 0.89 (s, 9H, (C*H*$_3$)$_3$CSi); 0.08 (s, 3H, (C*H*$_3$)$_2$Si), 0.07 (s, 3H, (C*H*$_3$)$_2$Si); $^{13}$C NMR (100 MHz, $CD_3OD$) δ 180.32, 156.04, 148.69, 148.31, 137.67, 119.89, 87.74, 84.23, 70.71, 63.01, 40.68, 35.54, 25.05, 17.92, 17.88, –6.68, –6.73; HRMS (ESI) m/z $M^+$calcd for $C_{20}H_{33}N_5O_5Si$ 451.2251; Found $[M + H]^+$ 452.2368

**5'-O-*tert*-butyldimethylsilyl-N$^2$-Isobutyryl-3'-*O*-methylthiomethyl-L-2'-deoxyguanosine (S-6).** To a stirred solution of **S-5** (1.7 g; 3.76 mmol) in DMSO (12 ml) was added acetic acid (6 ml) and acetic anhydride (18 ml). After stirred at room temperature for 48 h, the solution was added saturated $NaHCO_3$ solution at 0 °C and stirred for 30 min, and the aqueous layer was extracted with EtOAc (2 x 100 ml). The combined organic layers were dried over $Na_2SO_4$, filtered and concentrated. The crude product was purified by flash column chromatography (1:1 to 1:3, Hexanes–EtOAc) to afford **S-6** (1.2 g; 65%) as a light yellowish foam $R_f$ 0.26 (1:4 hexanes–EtOAc); $^1$H NMR (400 MHz, $CDCl_3$): δ 12.1 (s, 1H, N*H*), 9.40 (s, 1H, N*H*), 7.98 (s, 1H), 6.18 (app t, 1H, $J_{1',2'a} = J_{1',2'b}$ = 6.9 Hz, H-1'), 4.71–4.61 (m, 3H, C*H*$_2$S, H-3'), 4.15–4.09 (m, 1H, H-4'), 3.78 (d, 2H, *J* = 3.9 Hz, H-5'a, H-5'b), 2.74 (sep, 1H, *J* = 7.0 Hz, COC*H*($CH_3$)$_2$), 2.57–2.46 (m, 2H, H-2'a, H-2'b), 2.15 (s, 3H, SC*H*$_3$), 1.25 (app t, 6H, *J* = 7.0 Hz, COCH(C*H*$_3$)$_2$), 0.89 (s, 9H, (C*H*$_3$)$_3$CSi); 0.08 (s, 3H, (C*H*$_3$)$_2$Si), 0.07 (s, 3H, (C*H*$_3$)$_2$Si); $^{13}$C NMR (100 MHz, $CDCl_3$) δ 179.06, 156.04, 148.24, 147.82, 137.12, 121.32, 85.42, 84.21, 76.37, 73.77, 63.39, 38.42, 36.48, 26.06, 19.19, 19.10, 18.50, 14.03, –5.25, –5.36; HRMS (ESI) m/z $[M + H]^+$ calcd for $C_{22}H_{38}N_5O_5SSi$ 512.2363; Found 512.2360

**5’-O-*tert*-butyldimethylsilyl-O$^6$-diphenylcarbamoyl-N$^2$-Isobutyryl-3’-*O*-methylthiomethyl-L-2'-deoxyguanosine (S-7).** To a stirred solution of **S-6** (1.20 g, 2.35 mmol) in dry pyridine (25 ml) was added diphenylcarbamoyl chloride (814 mg, 3.52 mmol) and DIPEA (N,N-diisopropylethylamine) (1.23 ml, 7.03 mmol) at room temperature. After stirring at room temperature for 3 hours, the solvent was removed under high vacuum. The residue was diluted with EtOAc and washed with 2M HCl and saturated $NaHCO_3$. The organic layer was dried with $Na_2SO_4$, filtered and concentrated. The residue was purified by flash column chromatography (1:1 to 1:3, Hexanes–EtOAc) to afford **S-7** (1.5 g, 90%) as a foam-type yellowish powder. $R_f$ 0.5 (1:1 hexanes–EtOAc); $^1$H NMR (400 MHz, $CDCl_3$): δ 8.25 (s, 1H, N*H*), 8.05 (s, 1H), 7.50–7.33 (m, 8H, ArH), 7.29–7.21 (m, 2H, ArH), 6.41 (app t, 1H, $J_{1',2'a} = J_{1',2'b}$ = 6.7 Hz, H-1’), 4.76–4.66 (m, 3H, C*H*$_2$S, H-3’), 4.19–4.13 (m, 1H, H-4’), 3.88 (dd, 1H, $J_{5'a,4'}$ = 4.5 Hz, $J_{gem}$ = 11.1 Hz, H-5’a), 3.81 (dd, 1H, $J_{5'b,4'}$ = 3.6 Hz, $J_{gem}$ = 11.1 Hz, H-5’b), 2.96 (m, 1H, COC*H*($CH_3$)$_2$), 2.75 (m, 1H, H-2’a), 2.57 (ddd, 1H, $J_{2'a,1}$ = 6.7 Hz, $J_{2'a,3'}$ = 3.2 Hz, $J_{gem}$ = 13.6 Hz, H-2’b), 2.18 (s, 3H, SC*H*$_3$), 1.28 (d, 6H, *J* = 6.8 Hz, COCH(C*H*$_3$)$_2$), 0.92 (s, 9H, (C*H*$_3$)$_3$CSi); 0.11 (s, 3H, (C*H*$_3$)$_2$Si), 0.10 (s, 3H, (C*H*$_3$)$_2$Si); 13C NMR (100 MHz, $CDCl_3$) δ 175.60, 156.20, 154.71, 152.03, 150.63, 142.71, 141.99, 129.34, 121.60, 85.62, 85.02, 76.58, 73.8763.45, 38.24, 36.10, 26.13, 19.44, 18.55, 14.09, –5.19, –5.3; HRMS (ESI) m/z $[M + H]^+$ calcd for $C_{35}H_{47}N_6O_6SSi$ 707.3047; Found 707.3048

**3’-*O*-azidomethyl-O$^6$-diphenylcarbamoyl-N$^2$-Isobutyryl-L-2'-deoxyguanosine (S-8).** To a stirred solution of **S-7** (490 mg, 0.694 mmol) in dry $CH_2Cl_2$ (14 mL) was added cyclohexene (2.1 mL) and 1.0 M $SO_2Cl_2$ in $CH_2Cl_2$ (1.39 mL, 1.39 mmol) at 0 °C. After stirred at 0 °C for 1 hour, the volatiles were removed under reduced pressure. To a solution of the residue in dry DMF (14 mL) was added $NaN_3$ (271 mg, 4.61 mmol) at room temperature. After stirred at room temperature for 2 h, the reaction mixture was dispersed in distilled water (200 mL) and extracted with EtOAc (2 x 200 mL). The combined organic layer was dried over $Na_2SO_4$, filtered and concentrated under reduced pressure. To a solution of the residue win acetonitrile (14 mL) was added 2M HCl (7 mL) at 0 °C. The reaction mixture was stirred for 30 min. The solution was diluted with EtOAc (100 mL) and washed with saturated $NaHCO_3$. The organic layer was washed with water (2 x 50 mL) and the organic layers were dried over $Na_2SO_4$, filtered and concentrated under reduced pressure. The resulting residue was purified by flash column chromatography (1:2 to 0:1, Hexanes–EtOAc) to afford **S-8** (163 mg, 40%) as colorless oil. $R_f$ 0.32 (1:4 hexanes–EtOAc); $^1$H NMR (400 MHz,

$CD_3OD$) δ 8.55 (s, 1H,H-8), 7.56–7.43 (m, 4H, ArH), 7.42–7.34 (m, 4H, ArH), 7.31–7.22 (m, 2H, ArH), 6.48 (app t, 1H, $J_{1',2'a} = J_{1',2'b}$ = 6.5 Hz, H-1'), 4.87–4.80 (m, 3H, H-3', C*H*$_2$N$_3$), 4.16–4.11 (m, 1H, H-4'), 3.82 (dd, 1H, $J_{5'a,4'}$ = 3.9 Hz, $J_{gem}$ = 12.1 Hz, H-5'a), 3.76 (dd, 1H, $J_{5'b,4'}$ = 4.5 Hz, $J_{gem}$ = 12.1 Hz, H-5'b), 2.94 (app dt, 1H, $J_{2'b,1} = J_{2'a,3'}$ = 6.5 Hz, $J_{gem}$ = 13.8 Hz, H-2'a), 2.79 (sep, 1H, $J$ = 7.0 Hz, COC*H*($CH_3$)$_2$), 2.63 (ddd, 1H, $J_{2'a,1}$= 6.5 Hz, $J_{2'a,3'}$ = 4.3 Hz, $J_{gem}$ = 13.8 Hz, H-2'b); $^{13}C$ NMR (100 MHz, $CD_3OD$) δ 179.29, 156.08, 154.73, 152.81, 151.39, 144.79, 142.38, 129.44, 127.53, 121.49, 86.48, 85.42, 82.30, 79.07, 62.18, 39.57, 38.13, 36.08, 18.83; HRMS (ESI) m/z $[M + H]^+$ calcd for $C_{28}H_{30}N_9O_6$ 588.2219; Found 588.2177

**3'-*O*-azidomethyl-L-2'-deoxyguanosine triphosphate (S-9). S-8** (40 mg, 0.146 mmol) with proton sponge (37.59 mg, 0.175 mmol) was dried in a vacuum desiccator over $P_2O_5$ overnight. To a solution of **S-8** in trimethyl phosphate (0.49 mL) was added $POCl_3$ (21 uL, 0.219 mmol) dropwisely at 0 °C. The mixture was stirred at 0 °C for 1 hour and then was added a well-vortexed mixture of tributylammonium pyrophosphate (304 mg) and tributylamine (0.27 ml, 2.31 mmol) in anhydrous DMF (1.2 mL). The mixture was stirred for 20 min at room temperature and 0.1M triethylammonium bicarbonate buffer (TEAB buffer, pH 8.0, 0.1 M, 4 ml) was then added and the mixture was stirred for 3 hours at room temperature. The mixture was then added concentrated ammonium hydroxide (15 mL) and stirred overnight at room temperature. The resulting mixture was concentrated, and the residue was diluted with HPLC water (10.0 mL). The mixture was extracted with $CH_2Cl_2$ (3 x 10.0 mL) and the aqueous layer was concentrated under reduced pressure to 5.0 mL. The residue was subjected to preparative C18 HPLC (0%B to 90%B over 50 min, mobile phase A: 0.1M TEAB, B: ACN). The fractions with products were collected and concentrated under reduced pressure. The residue was purified with semi-prep HPLC column on an ion-exchange column (Agilent PL-SAX, 10 μm, 1000Å), mobile phase: A, 15% acetonitrile in water; B, 15% acetonitrile in 1M TEAB buffer. Elution was performed in gradient condition (0%B to 80%B over 50 min). The fractions with products were collected, and concentrated under reduced pressure, and then washed 3 times with HPLC grade water (5 mL). After NMR analysis, the pure product was lyophilized to afford **S-9** (2.7 mg, 4.5%) as a foam type solid.$^1H$ NMR (400 MHz, $D_2O$): δ 8.09 (s, 1H), 6.21 (m, 1H, H-1'), 4.83 (d, 1H, $J$ = 11.5 Hz, C*H*$_2$N$_3$), 4.77 (d, 1H, $J$ = 11.5 Hz, C*H*$_2$N$_3$), 4.64–4.61 (m, 1H), 4.37–4.31 (m, 1H), 4.16–4.03 (m, 2H, H-5'a, H-5'b), 2.81–2.72 (m, 1H, H-2'a), 2.61-2.50 (m, 1H, H-2'b); $^{31}P$ NMR (121.4 MHz, $D_2O$): δ –10.8 (bs, 1P), -11.5 (d,

1P, *J* = 18.4 Hz), -23.2 (bs, 1P). HRMS (ESI) m/z calcd for [M] $C_{11}H_{17}N_8O_{13}P_3$ 562.0128; Found $[M – H]^-$ 561.0072.

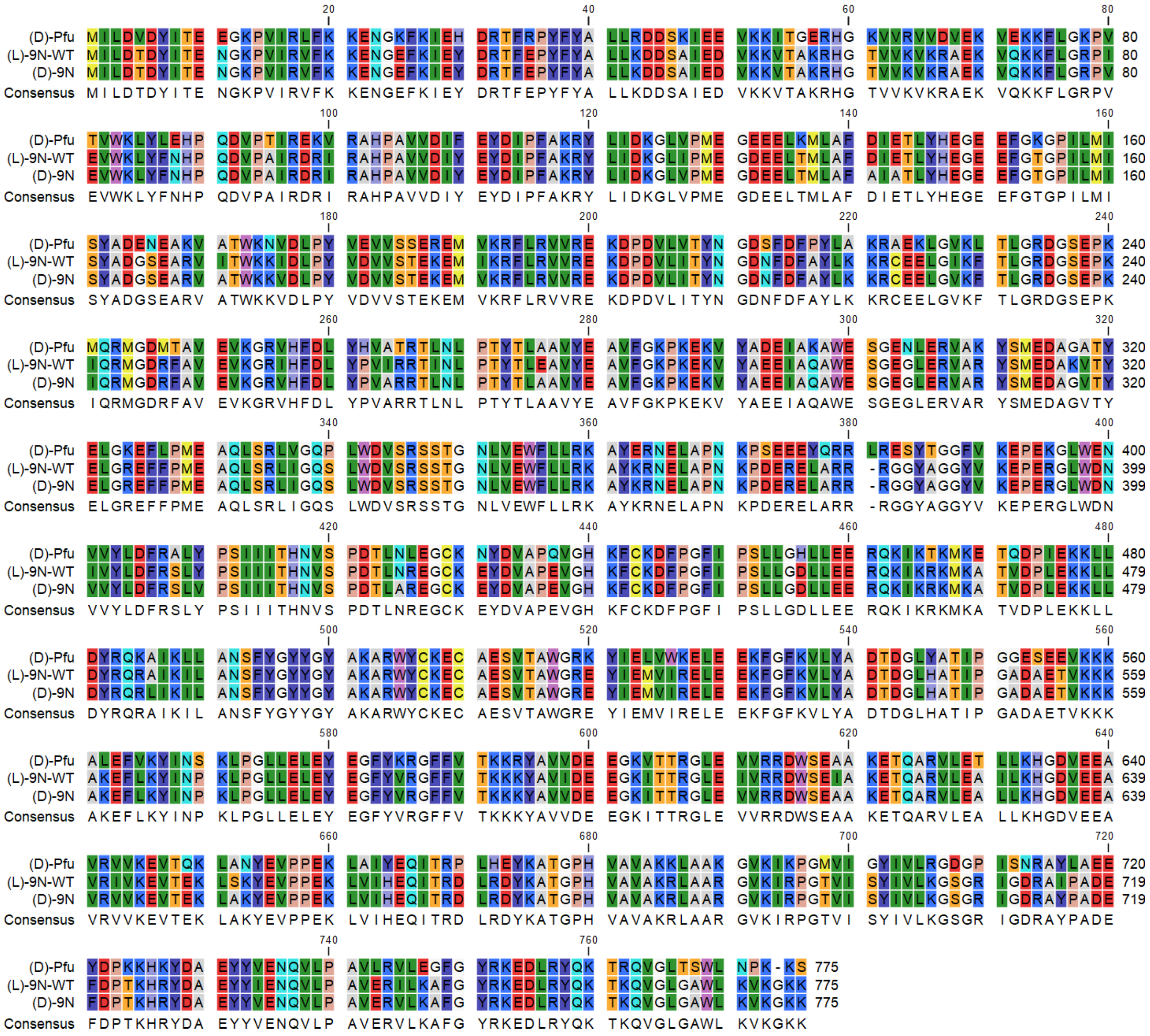

**Supplementary Figure 1.** Amino acid sequence alignment of mirror image Pfu, natural 9°N wild-type and mirror image 9°N containing mutations at D142A, E144A, Y409V and A485L for function and E277A and N425A for native chemical ligation sites.

**(a) D-9°N-N-1:**

**HHHHHH**MILDTDYITENGKPVIRVFKKENGEFKIEYDRTFEPYFYALLKDDSAIEDVKKVT-$NHNH_2$

**(b)**

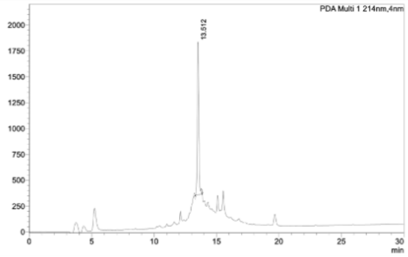

**(c)**

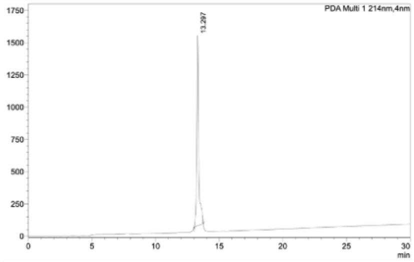

**(d)**

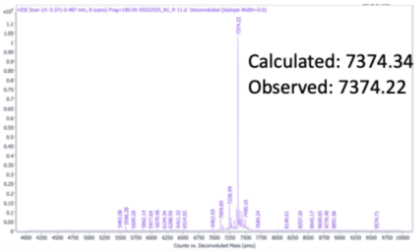

**Supplementary Figure 2.** (a) Sequence of D-9°N-N-1; (b) Analytical HPLC chromatogram of D-9°N-N-1 crude (λ=214 nm); (c) Analytical HPLC chromatogram of the purified D-9°N-N-1 (λ=214 nm); (d) Deconvoluted MS (LC-QTOF) spectrum of purified D-9°N-N-1. Analytical Method: Column: Welch XB-C4, 4.6x250mm, 5µm. Buffers: A: 0.1%TFA/$H_2O$, B: 0.1%TFA/$CH_3CN$. Gradient: 20 to 70% of Buffer B over 30min; Purification Method: Buffers: A: 0.1%TFA/$H_2O$, B: 0.1%TFA/$CH_3CN$. Gradient: 20 to 70% of Buffer B over 50min. Column: Agilent Polaris C18-A, 21.2x250mm, 5um, 180Å.

**(a) D-9°N-N-2:**

Thz-KRHGTVVKVKRAEKVQKKFLGRPVEVWKLYFNHPQDVPAIRDRIR-$NHNH_2$

**(b)**

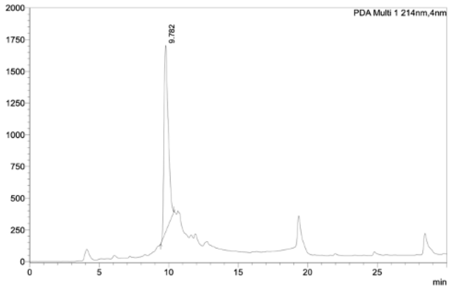

**(c)**

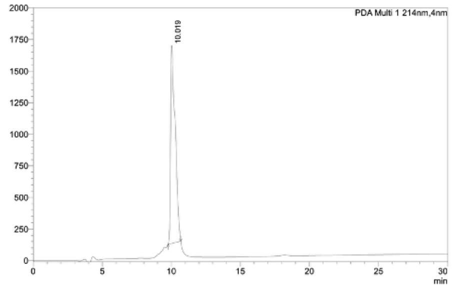

**(d)**

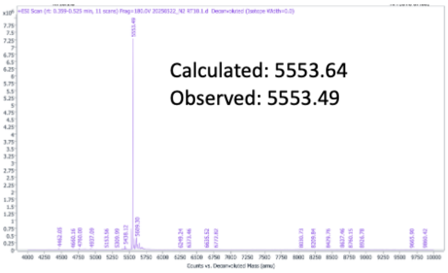

**Supplementary Figure 3.** (a) Sequence of D-9°N-N-2; (b) Analytical HPLC chromatogram of D-9°N-N-2 crude (λ=214 nm); (c) Analytical HPLC chromatogram of the purified D-9°N-N-2 (λ=214 nm); (d) Deconvoluted MS (LC-QTOF) spectrum of purified D-9°N-N-2. Analytical Method: Column: Welch XB-C4, 4.6x250mm, 5µm. Buffers: A: 0.1%TFA/$H_2O$, B: 0.1%TFA/$CH_3CN$. Gradient: 20 to 70% of Buffer B over 30min; Purification Method: Buffers: A: 0.1%TFA/$H_2O$, B: 0.1%TFA/$CH_3CN$. Gradient: 20 to 70% of Buffer B over 50min. Column: Agilent Polaris C18-A, 21.2x250mm, 5um, 180Å.

**(a) D-9°N-N-3:**

CHPAVVDIYEYDIPFAKRYLIDKGLVPMEGDEELTMLAFAIATLYHEGEEFGTGPILMISY-$NHNH_2$

**(b)**

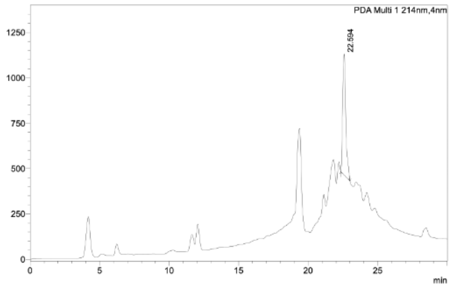

**(c)**

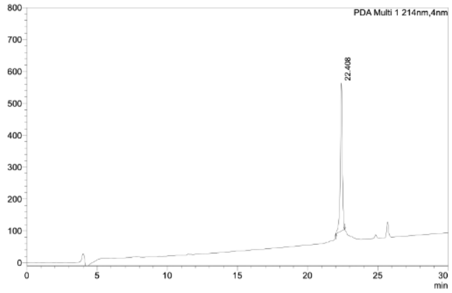

**(d)**

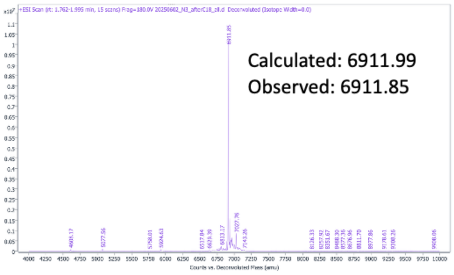

**Supplementary Figure 4.** (a) Sequence of D-9°N-N-3; (b) Analytical HPLC chromatogram of D-9°N-N-3 crude (λ=214 nm); (c) Analytical HPLC chromatogram of the purified D-9°N-N-3 (λ=214 nm); (d) Deconvoluted MS (LC-QTOF) spectrum of purified D-9°N-N-3. Analytical Method: Column: Welch XB-C4, 4.6x250mm, 5µm. Buffers: A: 0.1%TFA/$H_2O$, B: 0.1%TFA/$CH_3CN$. Gradient: 20 to 70% of Buffer B over 30min; Purification Method: Buffers: A: 0.1%TFA/$H_2O$, B: 0.1%TFA/$CH_3CN$. Gradient: 30 to 70% of Buffer B over 50min. Column: Agilent Polaris C18-A, 21.2x250mm, 5um, 180Å.

**(a) D-9°N-N-4:**

Thz-DGSEARVATWKKVDLPYVDVVSTEKEMVKRFLRVVREKDPDVLITYNGDNFDF-$NHNH_2$

(b) (c) (d)

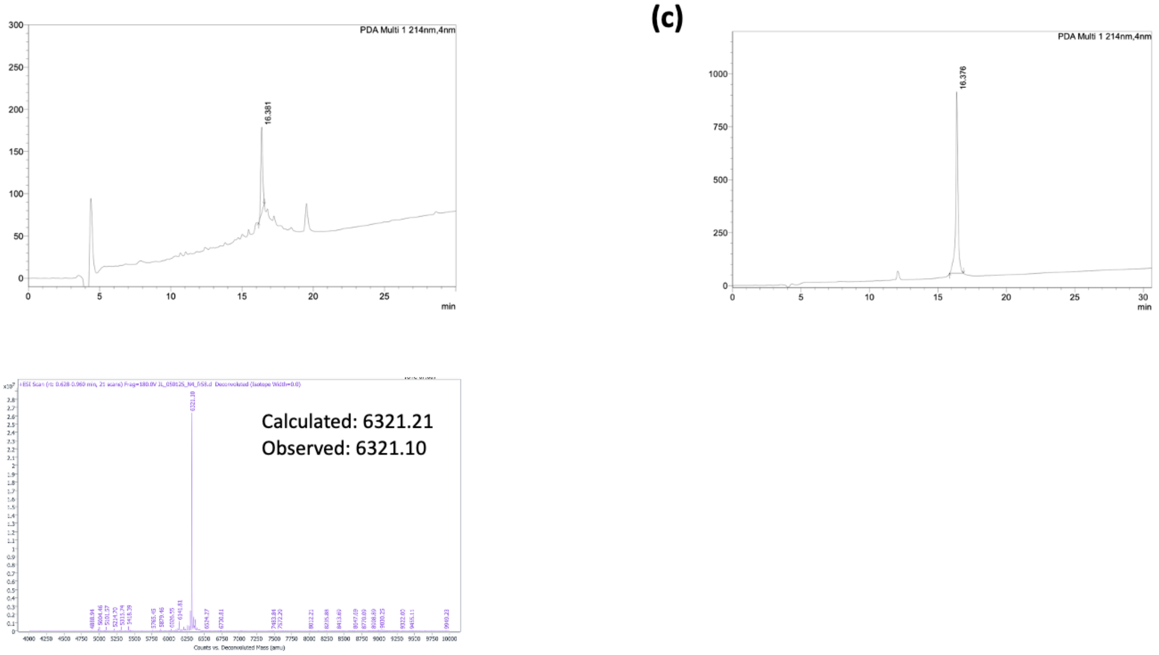

**Supplementary Figure 5.** (a) Sequence of D-9°N-N-4; (b) Analytical HPLC chromatogram of D-9°N-N-4 crude (λ=214 nm); (c) Analytical HPLC chromatogram of the purified D-9°N-N-4 (λ=214 nm); (d) Deconvoluted MS (LC-QTOF) spectrum of purified D-9°N-N-4. Analytical Method: Column: Welch XB-C4, 4.6x250mm, 5μm. Buffers: A: 0.1%TFA/$H_2O$, B: 0.1%TFA/$CH_3CN$. Gradient: 20 to 70% of Buffer B over 30min; Purification Method: Buffers: A: 0.1%TFA/$H_2O$, B: 0.1%TFA/$CH_3CN$. Gradient: 30 to 50% of Buffer B over 40min. Column: Agilent Polaris C18-A, 21.2x250mm, 5um, 180Å.

**(a) D-9°N-N-5:**

CYLKKRC(Acm)EELGVKFTLGRDGSEPKIQRMGDRFAVEVKGRVHFDLYPVARRTLNLPTYTL-$NHNH_2$

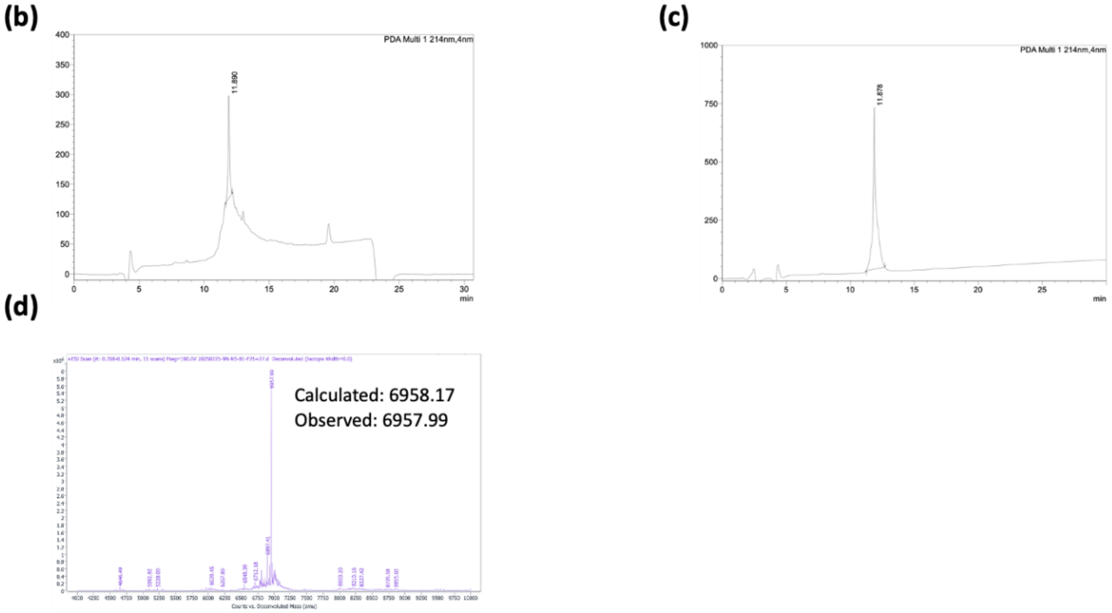

**Supplementary Figure 6.** (a) Sequence of D-9°N-N-5; (b) Analytical HPLC chromatogram of D-9°N-N-5 crude (λ=214 nm); (c) Analytical HPLC chromatogram of the purified D-9°N-N-5 (λ=214 nm); (d) Deconvoluted MS (LC-QTOF) spectrum of purified D-9°N-N-5. Analytical Method: Column: Welch XB-C4, 4.6x250mm, 5µm. Buffers: A: 0.1%TFA/$H_2O$, B: 0.1%TFA/$CH_3CN$. Gradient: 20 to 70% of Buffer B over 30min; Purification Method: Buffers: A: 0.1%TFA/$H_2O$, B: 0.1%TFA/$CH_3CN$. Gradient: 20 to 70% of Buffer B over 50min. Column: Agilent Polaris C18-A, 21.2x250mm, 5um, 180Å.

**(a) D-9°N-N-6:**

Thz-AVYEAVFGKPKEKVYAEEIAQAWESGEGLERVARYSMED-$NHNH_2$

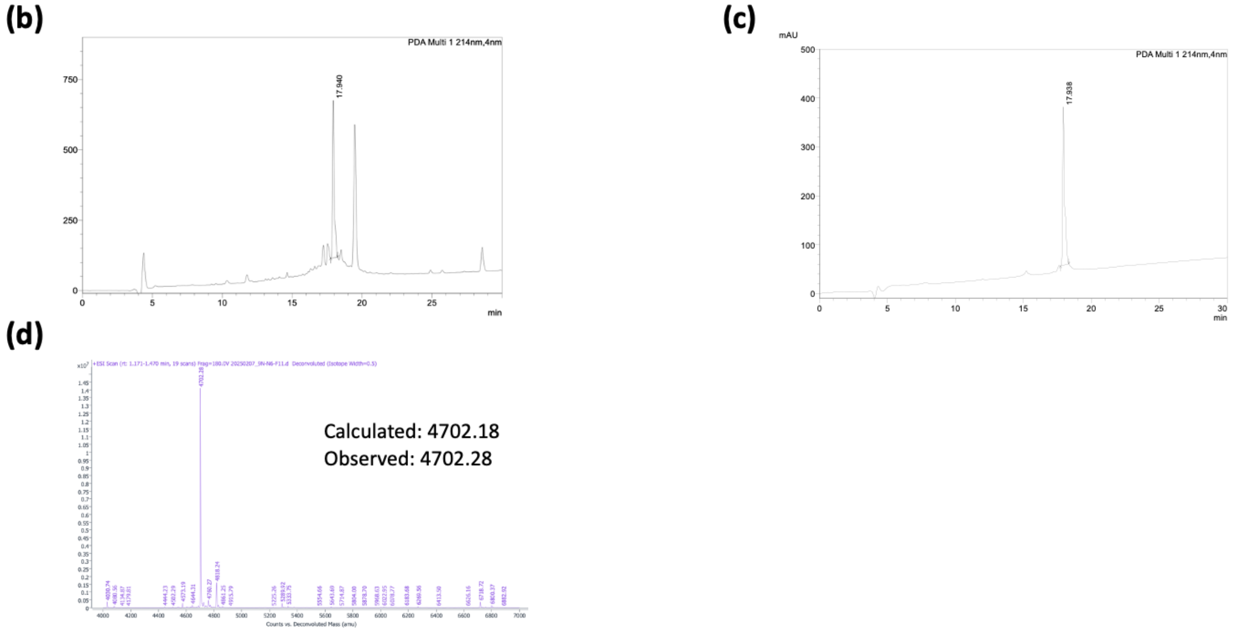

**Supplementary Figure 7.** (a) Sequence of D-9°N-N-6; (b) Analytical HPLC chromatogram of D-9°N-N-6 crude (λ=214 nm); (c) Analytical HPLC chromatogram of the purified D-9°N-N-6 (λ=214 nm); (d) Deconvoluted MS (LC-QTOF) spectrum of purified D-9°N-N-6. Analytical Method: Column: Welch XB-C4, 4.6x250mm, 5µm. Buffers: A: 0.1%TFA/$H_2O$, B: 0.1%TFA/$CH_3CN$. Gradient: 20 to 70% of Buffer B over 30min; Purification Method: Buffers: A: 0.1%TFA/$H_2O$, B: 0.1%TFA/$CH_3CN$. Gradient: 30 to 70% of Buffer B over 40min. Column: Agilent Polaris C18-A, 21.2x250mm, 5um, 180Å.

**(a) D-9°N-N-7:**

Thz-GVTYELGREFFPMEAQLSRLIGQS LWDVSRSSTGNLVEWFLLRKAYKRNEL-$NHNH_2$

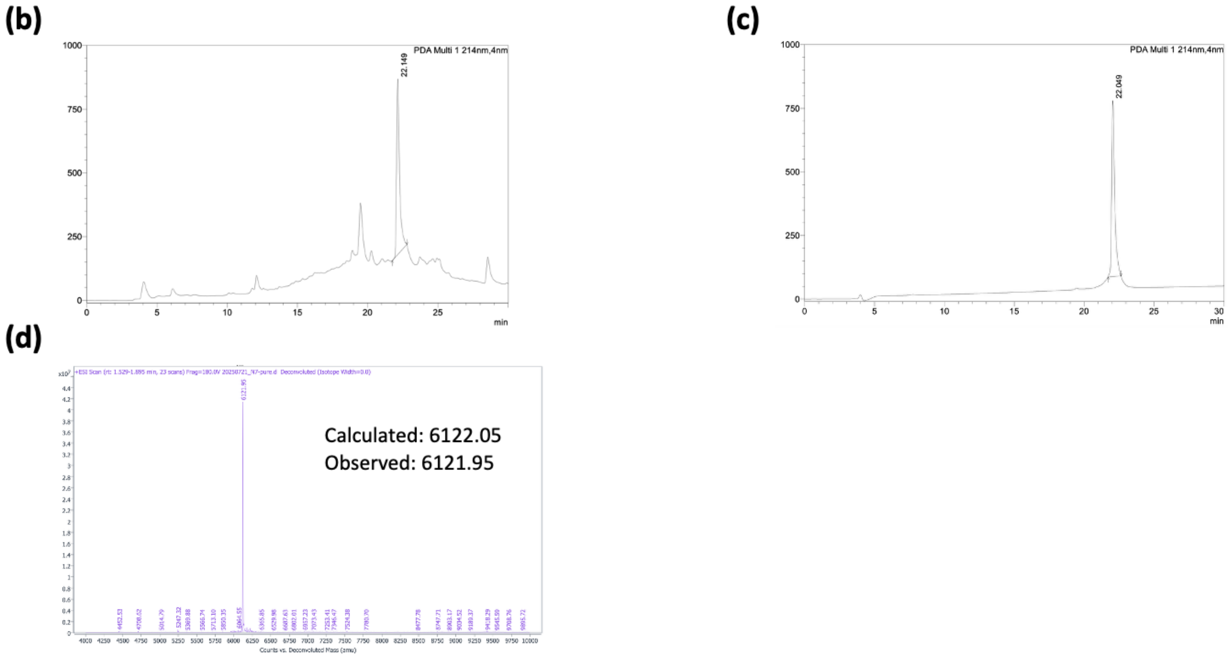

**Supplementary Figure 8.** (a) Sequence of D-9°N-N-7; (b) Analytical HPLC chromatogram of D-9°N-N-7 crude (λ=214 nm); (c) Analytical HPLC chromatogram of the purified D-9°N-N-7 (λ=214 nm); (d) Deconvoluted MS (LC-QTOF) spectrum of purified D-9°N-N-7. Analytical Method: Column: Welch XB-C4, 4.6x250mm, 5µm. Buffers: A: 0.1%TFA/$H_2O$, B: 0.1%TFA/$CH_3CN$. Gradient: 20 to 70% of Buffer B over 30min; Purification Method: Buffers: A: 0.1%TFA/$H_2O$, B: 0.1%TFA/$CH_3CN$. Gradient: 30 to 70% of Buffer B over 50min. Column: Agilent Polaris C18-A, 21.2x250mm, 5um, 180Å.

**(a) D-9°N-N-8:**

CPNKPDERELARRRGGYAGGYV KEPERGLWDNVVYLDFRSLVPSIIITHNVSPDTL-$NHNH_2$

**(b)**

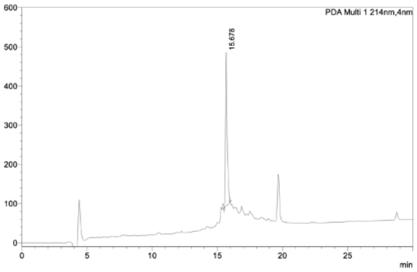

**(c)**

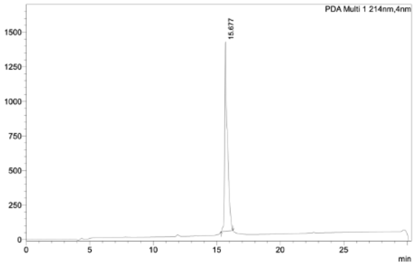

**(d)**

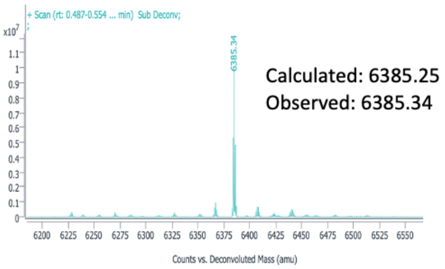

**Supplementary Figure 9.** (a) Sequence of D-9°N-N-8; (b) Analytical HPLC chromatogram of D-9°N-N-8 crude (λ=214 nm); (c) Analytical HPLC chromatogram of the purified D-9°N-N-8 (λ=214 nm); (d) Deconvoluted MS (LC-QTOF) spectrum of purified D-9°N-N-8. Analytical Method: Column: Welch XB-C4, 4.6x250mm, 5μm. Buffers: A: 0.1%TFA/$H_2O$, B: 0.1%TFA/$CH_3CN$. Gradient: 20 to 70% of Buffer B over 30min; Purification Method: Buffers: A: 0.1%TFA/$H_2O$, B: 0.1%TFA/$CH_3CN$. Gradient: 20 to 70% of Buffer B over 50min. Column: Agilent Polaris C18-A, 21.2x250mm, 5um, 180Å.

**(a) D-9°N-N-9:**

CREGCKEYDVAPEVGHKFCKDFPGFI PSLLGDLLEERQKIKRK-OH

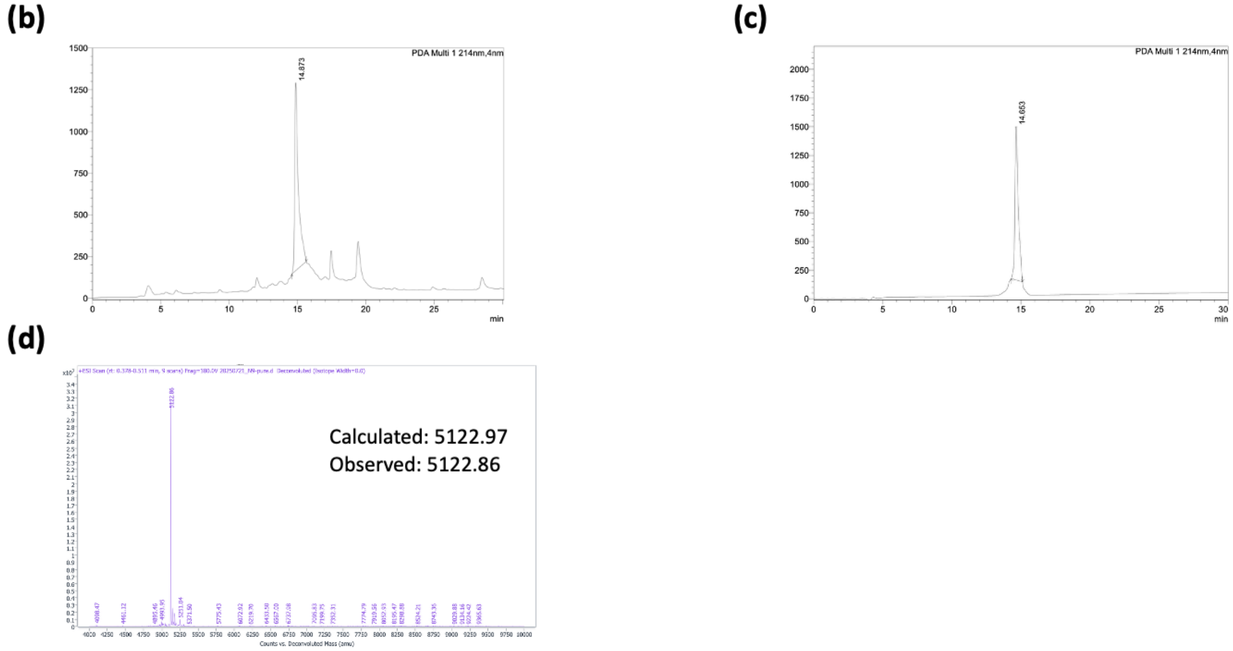

**Supplementary Figure 10.** (a) Sequence of D-9°N-N-9; (b) Analytical HPLC chromatogram of D-9°N-N-9 crude (λ=214 nm); (c) Analytical HPLC chromatogram of the purified D-9°N-N-9 (λ=214 nm); (d) Deconvoluted MS (LC-QTOF) spectrum of purified D-9°N-N-8. Analytical Method: Column: Welch XB-C4, 4.6x250mm, 5μm. Buffers: A: 0.1%TFA/$H_2O$, B: 0.1%TFA/$CH_3CN$. Gradient: 20 to 70% of Buffer B over 30min; Purification Method: Buffers: A: 0.1%TFA/$H_2O$, B: 0.1%TFA/$CH_3CN$. Gradient: 25 to 70% of Buffer B over 45min. Column: Agilent Polaris C18-A, 21.2x250mm, 5um, 180Å.

**(a) D-9°N-N-10:**

CKRHGTVVKVKRAEKVQKKFLGRPVEVWKLYFNHPQDVPAIRDRIRCHPAVVDIYEYDIPFAKRYLIDKGLVPMEGDE
ELTMLAFAIATLYHEGEEFGTGPILMISY-$NHNH_2$

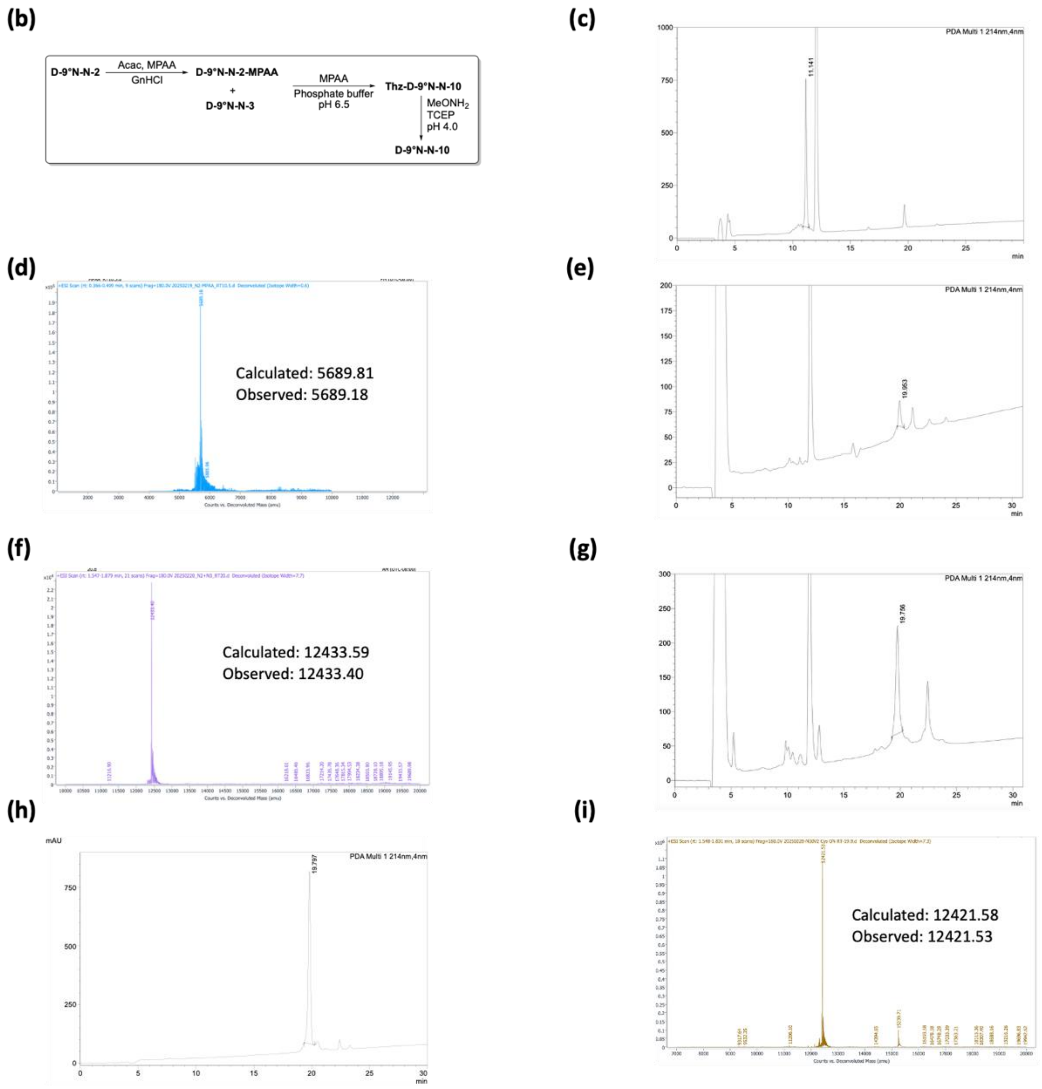

**Supplementary Figure 11.** (a) Peptide sequence of D-9°N-N-10; (b) Reaction Scheme to obtain D-9°N-N-10; (c) Analytical HPLC chromatogram of D-9°N-N-2 thioesterification crude (λ=214 nm); (d) Deconvoluted MS (LC-QTOF) spectrum of D-9°N-N-2-MPAA from collection of analytical HPLC ; (e) Analytical HPLC chromatogram of Thz-D-9°N-N-10 crude (λ=214 nm); (f) Deconvoluted MS (LC-QTOF) spectrum of Thz-D-9°N-N-10 from collection of analytical HPLC; (g) Analytical HPLC chromatogram of D-9°N-N-10 crude (λ=214 nm); (h) Analytical HPLC chromatogram of purified D-9°N-N-10 (λ=214 nm); (i) Deconvoluted MS (LC-QTOF) spectrum of purified D-9°N-N-10; Analytical Method: Column: Welch XB-C4, 4.6x250mm, 5μm. Buffers:

A: 0.1%TFA/$H_2O$, B: 0.1%TFA/$CH_3CN$. Gradient: 20 to 70% of Buffer B over 50min; Purification Method: Buffers: A: 0.1%TFA/$H_2O$, B: 0.1%TFA/$CH_3CN$. Gradient: 20 to 70% of Buffer B over 50min. Column: Welch Ultisil XB-C4 21.2x250mm, 5um, 300Å

**(a) D-9°N-N-11:**

**HHHHHH**MILDTDYITENGKPVIRVFKKENGEFKIEYDRTFEPYFYALLKDDSAIEDVKKVTCKRHGTVVKVKRAEKVQ
KKFLGRPVEVWKLYFNHPQDVPAIRDRIRCHPAVVDIYEYDIPFAKRYLIDKGLVPMEGDEELTMLAFAIATLYHEGEEF
GTGPILMISY-NHNH$_2$

**(b)**

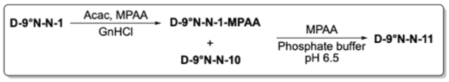

**(c)**

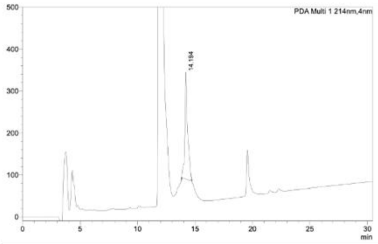

**(d)**

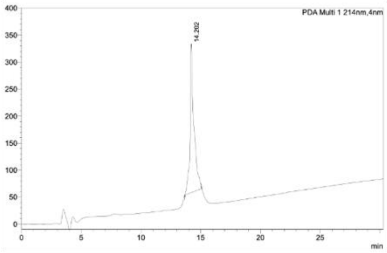

**(e)**

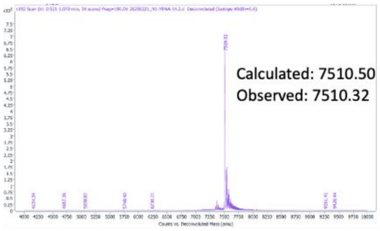

**(f)**

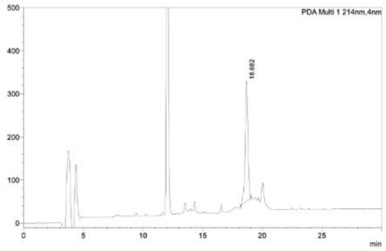

**(g)**

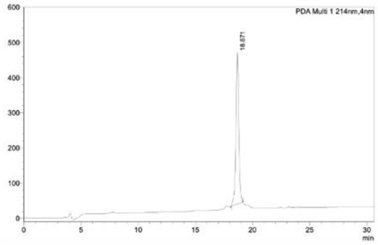

**(h)**

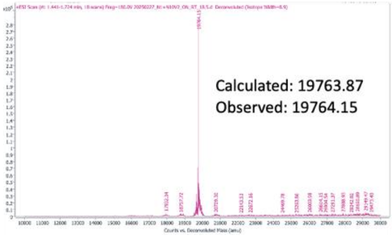

**Supplementary Figure 12.** (a) Peptide sequence of D-9°N-N-11; (b) Reaction Scheme to obtain D-9°N-N-11; (c) Analytical HPLC chromatogram of D-9°N-N-1 thioesterification crude (λ=214 nm); (d) Analytical HPLC chromatogram of purified D-9°N-N-1-MPAA (λ=214 nm); (e) Deconvoluted MS (LC-QTOF) spectrum of purified D-9°N-N-1-MPAA; (f) Analytical HPLC chromatogram of D-9°N-N-11 crude (λ=214 nm); (g) Analytical HPLC chromatogram of purified D-9°N-N-11 (λ=214 nm); (H) Deconvoluted MS (LC-QTOF) spectrum of purified D-9°N-N-11; Analytical Method: Column: Welch XB-C4, 4.6x250mm, 5μm. Buffers: A: 0.1%TFA/$H_2O$, B: 0.1%TFA/$CH_3CN$. Gradient: 20 to 70% of Buffer B over 50min; Purification Method: Buffers: A:

0.1%TFA/$H_2O$, B: 0.1%TFA/$CH_3CN$. Gradient: 20 to 70% of Buffer B over 50min. Column: Welch Ultisil XB-C4 21.2x250mm, 5um, 300Å

**(a) D-9°N-N-12:**

CDGSEARVATWKKVDLPYVDVVSTEKEMVKRFLRVVREKDPDVLITYNGDNFDFCYLKKRC(Acm)EELGVKFTLGRD GSEPKIQRMGDRFAVEVKGRVHFDLYPVARRTLNLPTYTL-$NHNH_2$

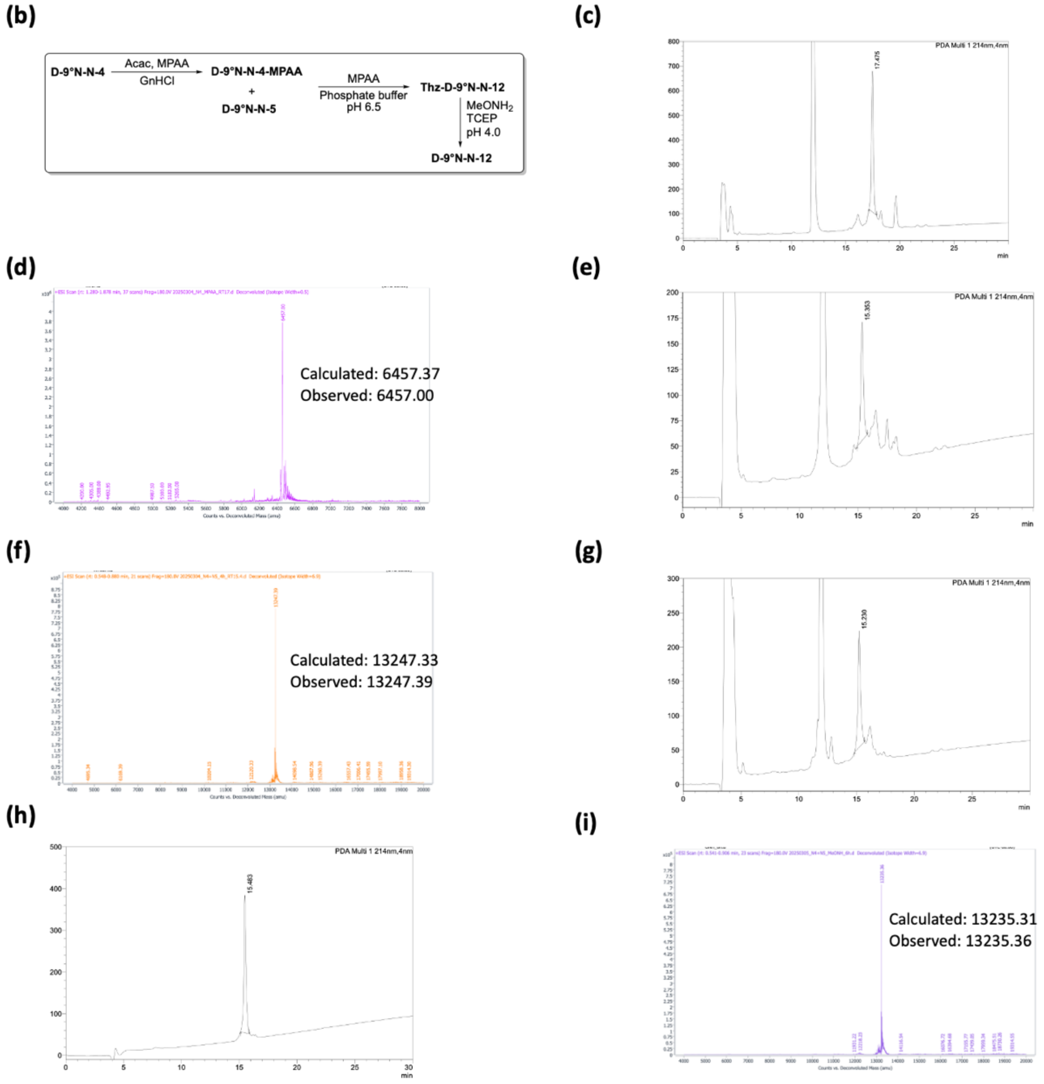

**Supplementary Figure 13.** (a) Peptide sequence of D-9°N-N-12; (b) Reaction Scheme to obtain D-9°N-N-12; (c) Analytical HPLC chromatogram of D-9°N-N-4 thioesterification crude (λ=214 nm); (d) Deconvoluted MS (LC-QTOF) spectrum of D-9°N-N-4-MPAA from collection of analytical HPLC ; (e) Analytical HPLC chromatogram of Thz-D-9°N-N-12 crude (λ=214 nm); (f) Deconvoluted MS (LC-QTOF) spectrum of Thz-D-9°N-N-12 from collection of analytical HPLC; (g) Analytical HPLC chromatogram of D-9°N-N-12 crude (λ=214 nm); (h) Analytical HPLC chromatogram of purified D-9°N-N-12 (λ=214 nm); (i) Deconvoluted MS (LC-QTOF) spectrum of purified D-9°N-N-12; Analytical Method: Column: Welch XB-C4, 4.6x250mm, 5µm. Buffers:

A: 0.1%TFA/$H_2O$, B: 0.1%TFA/$CH_3CN$. Gradient: 20 to 70% of Buffer B over 50min; Purification Method: Buffers: A: 0.1%TFA/$H_2O$, B: 0.1%TFA/$CH_3CN$. Gradient: 20 to 70% of Buffer B over 50min. Column: Welch Ultisil XB-C4 21.2x250mm, 5um, 300Å

**(a) D-9°N-N-13:**

Thz-CGVTYELGREFFPMEAQLSRLIGQSLWDVSRSSTGNLVEWFLLRKAYKRNELCPNKPDERELARRRGGYAGGYVKEPE RGLWDNVVYLDFRSLVPSIIITHNVSPDTL-$NHNH_2$

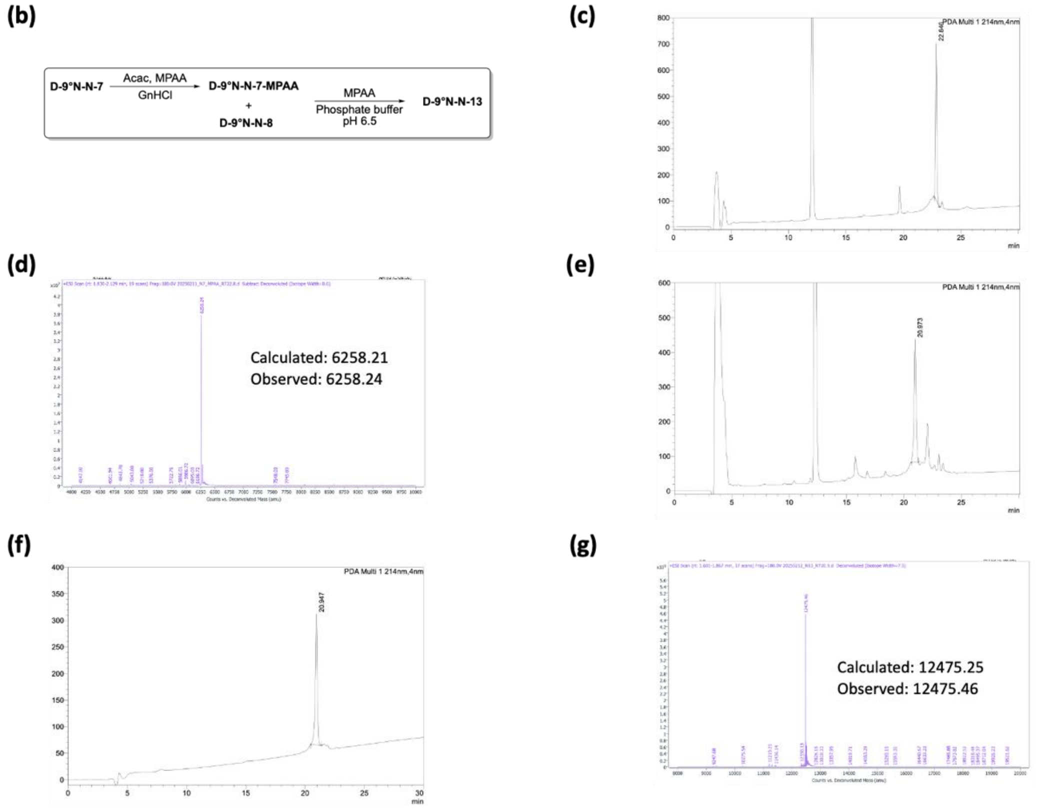

**Supplementary Figure 14.** (a) Peptide sequence of D-9°N-N-13; (b) Reaction Scheme to obtain D-9°N-N-13; (c) Analytical HPLC chromatogram of D-9°N-N-7 thioesterification crude (λ=214 nm); (d) Deconvoluted MS (LC-QTOF) spectrum of D-9°N-N-7-MPAA from collection of analytical HPLC ; (e) Analytical HPLC chromatogram of D-9°N-N-13 crude (λ=214 nm); (f) Analytical HPLC chromatogram of purified D-9°N-N-13 (λ=214 nm); (g) Deconvoluted MS (LC-QTOF) spectrum of purified D-9°N-N-13; Analytical Method: Column: Welch XB-C4, 4.6x250mm, 5µm. Buffers: A: 0.1%TFA/$H_2O$, B: 0.1%TFA/$CH_3CN$. Gradient: 20 to 70% of Buffer B over 50min; Purification Method: Buffers: A: 0.1%TFA/$H_2O$, B: 0.1%TFA/$CH_3CN$. Gradient: 20 to 70% of Buffer B over 50min. Column: Welch Ultisil XB-C4 21.2x250mm, 5um, 300Å

**(a) D-9°N-N-14:**

CGVTYELGREFFPMEAQLSRLIGQSLWDVSRSSTGNLVEWFLLRKAYKRNELCPNKPDERELARRRGGYAGGYVKEPERGL
WDNVVYLDFRSLVPSIIITHNVSPDTLCREGC(Acm)KEYDVAPEVGHKFC(Acm)KDFPGFIPSLLGDLLEERQKIKRK-OH

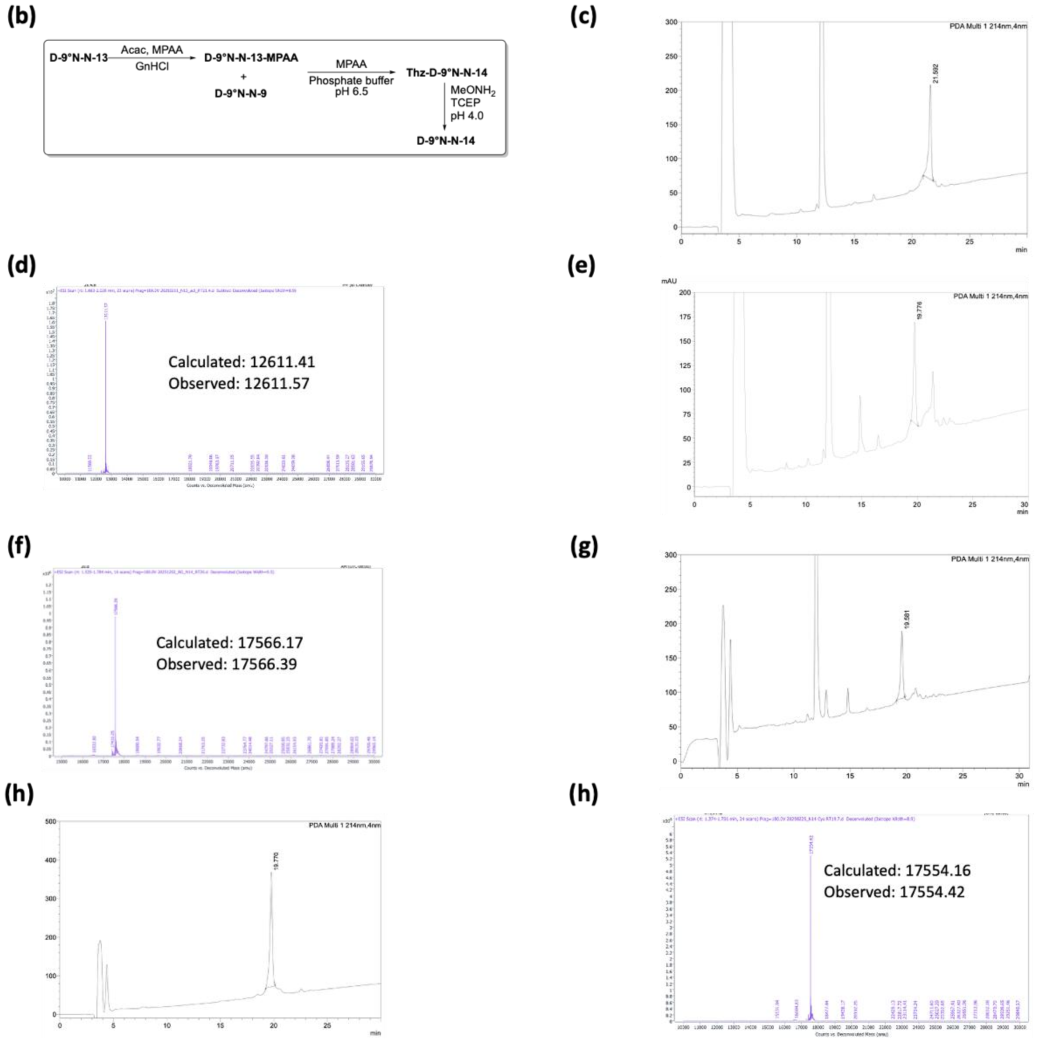

**Supplementary Figure 15.** (a) Peptide sequence of D-9°N-N-14; (b) Reaction Scheme to obtain D-9°N-N-14; (c) Analytical HPLC chromatogram of D-9°N-N-13 thioesterification crude (λ=214 nm); (d) Deconvoluted MS (LC-QTOF) spectrum of D-9°N-N-13-MPAA from collection of analytical HPLC ; (e) Analytical HPLC chromatogram of Thz-D-9°N-N-14 crude (λ=214 nm); (f) Deconvoluted MS (LC-QTOF) spectrum of Thz-D-9°N-N-14 from collection of analytical HPLC; (g) Analytical HPLC chromatogram of D-9°N-N-14 crude (λ=214 nm); (h) Analytical HPLC chromatogram of purified D-9°N-N-14 (λ=214 nm); (i) Deconvoluted MS (LC-QTOF) spectrum of purified D-9°N-N-14; Analytical Method: Column: Welch XB-C4, 4.6x250mm, 5μm. Buffers: A: 0.1%TFA/$H_2O$, B: 0.1%TFA/$CH_3CN$. Gradient: 20 to 70% of Buffer B over 50min; Purification

Method: Buffers: A: 0.1%TFA/$H_2O$, B: 0.1%TFA/$CH_3CN$. Gradient: 20 to 70% of Buffer B over 50min. Column: Welch Ultisil XB-C4 21.2x250mm, 5um, 300Å

**(a) D-9°N-N-15:**

**HHHHHH**MILDTDYITENGKPVIRVFKKENGEFKIEYDRTFEPYFYALLKDDSAIEDVKKVTCKRHGTVVKVKRAEKVQKKFL
GRPVEVWKLYFNHPQDVPAIRDRIRCHPAVVDIYEYDIPFAKRYLIDKGLVPMEGDEELTMLAFAIATLYHEGEEFGTGPILMI
SYCDGSEARVATWKKVDLPYVDVVSTEKEMVKRFLRVVREKDPDVLITYNGDNFDFCYLKKRC(Acm)EELGVKFTLGRDG
SEPKIQRMGDRFAVEVKGRVHFDLYPVARRTLNLPTYTL-NHNH$_2$

(b)

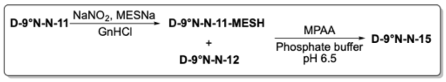

(c)

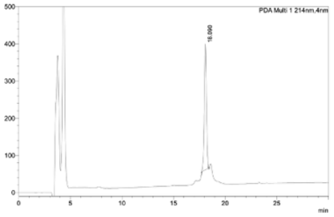

(d)

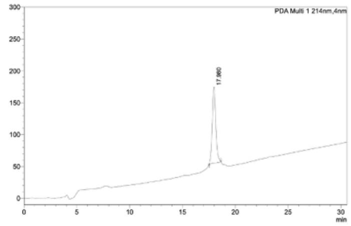

(e)

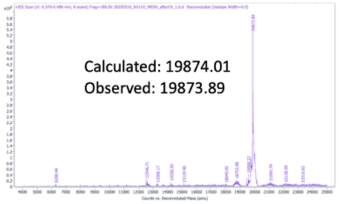

(f)

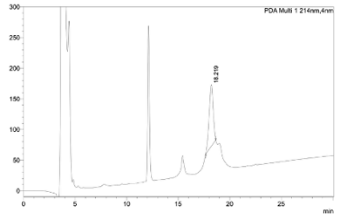

(g)

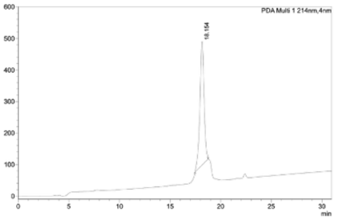

(h)

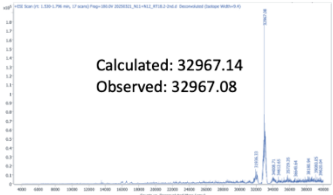

**Supplementary Figure 16.** (a) Peptide sequence of D-9°N-N-15; (b) Reaction Scheme to obtain D-9°N-N-15; (c) Analytical HPLC chromatogram of D-9°N-N-11 thioesterification crude (λ=214 nm); (d) Analytical HPLC chromatogram of purified D-9°N-N-11-MESH (λ=214 nm); (e) Deconvoluted MS (LC-QTOF) spectrum of purified D-9°N-N-11-MESH; (f) Analytical HPLC chromatogram of D-9°N-N-15 crude (λ=214 nm); (g) Analytical HPLC chromatogram of purified D-9°N-N-15 (λ=214 nm); (h) Deconvoluted MS (LC-QTOF) spectrum of purified D-9°N-N-15; Analytical Method: Column: Welch XB-C4, 4.6x250mm, 5µm. Buffers: A: 0.1%TFA/$H_2O$, B: 0.1%TFA/$CH_3CN$. Gradient: 20 to 70% of Buffer B over 50min; Purification Method: Buffers: A:

0.1%TFA/$H_2O$, B: 0.1%TFA/$CH_3CN$. Gradient: 20 to 70% of Buffer B over 50min. Column: Welch Ultisil XB-C4 21.2x250mm, 5um, 300Å

**(a) D-9°N-N-16:**

**HHHHHH**MILDTDYITENGKPVIRVFKKENGEFKIEYDRTFEPYFYALLKDDSAIEDVKKVTAKRHGTVVKVKRAEKVQKKFL
GRPVEVWKLYFNHPQDVPAIRDRIRAHPAVVDIYEYDIPFAKRYLIDKGLVPMEGDEELTMLAFAIATLYHEGEEFGTGPILMI
SYADGSEARVATWKKVDLPYVDVVSTEKEMVKRFLRVVREKDPDVLITYNGDNFDFAYLKKRC(Acm)EELGVKFTLGRDG
SEPKIQRMGDRFAVEVKGRVHFDLYPVARRTLNLPTYTL-$NHNH_2$

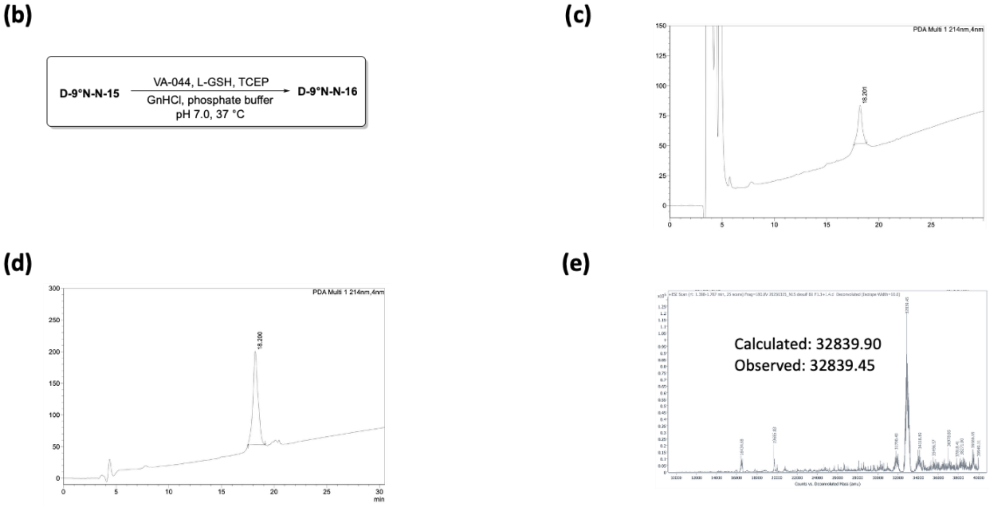

**Supplementary Figure 17.** (a) Peptide sequence of D-9°N-N-16; (b) Reaction Scheme to obtain D-9°N-N-16; (c) Analytical HPLC chromatogram of D-9°N-N-16 crude (λ=214 nm); (d) Analytical HPLC chromatogram of purified D-9°N-N-16 (λ=214 nm); (e) Deconvoluted MS (LC-QTOF) spectrum of purified D-9°N-N-16; Analytical Method: Column: Welch XB-C4, 4.6x250mm, 5µm. Buffers: A: 0.1%TFA/$H_2O$, B: 0.1%TFA/$CH_3CN$. Gradient: 20 to 70% of Buffer B over 50min; Purification Method: Buffers: A: 0.1%TFA/$H_2O$, B: 0.1%TFA/$CH_3CN$. Gradient: 30 to 70% of Buffer B over 40 min. Column: Welch Ultisil XB-C4 21.2x250mm, 5um, 300Å

**(a) D-9°N-N-17:**

CAVYEAVFGKPKEKVYAEEIAQAWESGEGLERVARYSMEDAGVTYELGREFFPMEAQLSRLIGQSLWDVSRSSTGNLVEW
FLLRKAYKRNELAPNKPDERELARRRGGYAGGYVKEPERGLWDNVVYLDFRSLVPSIIITHNVSPDTLAREGC(Acm)KEYDV
APEVGHKFC(Acm)KDFPGFIPSLLGDLLEERQKIKRK-OH

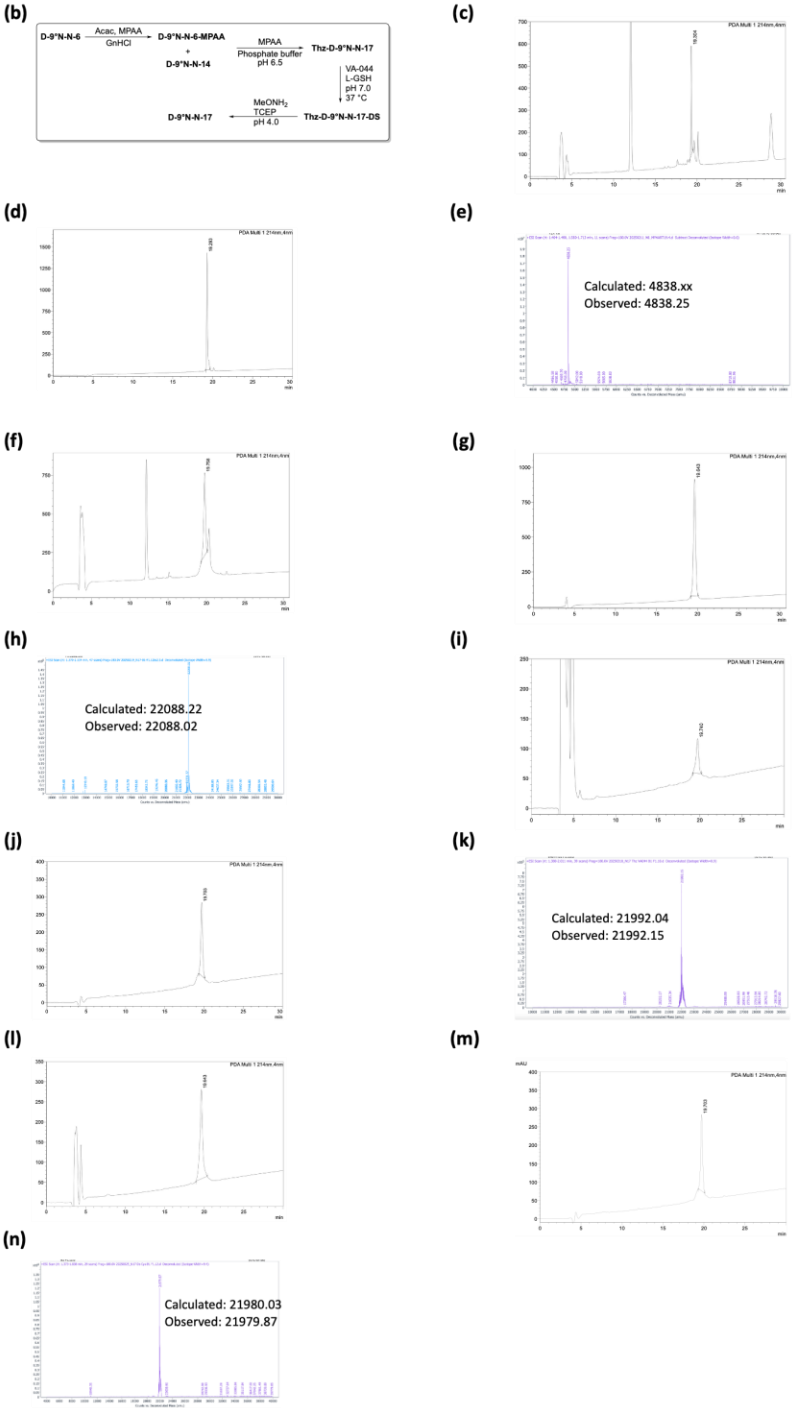

**Supplementary Figure 18.** (a) Peptide sequence of D-9°N-N-17; (b) Reaction Scheme to obtain D-9°N-N-17; (c) Analytical HPLC chromatogram of D-9°N-N-6 thioesterification crude (λ=214

nm); (d) Analytical HPLC chromatogram of purified D-9°N-N-6-MPAA (λ=214 nm); (e) Deconvoluted MS (LC-QTOF) spectrum of purified D-9°N-N-6-MPAA; (f) Analytical HPLC chromatogram of Thz-D-9°N-N-17 crude (λ=214 nm); (g) Analytical HPLC chromatogram of purified Thz-D-9°N-N-17 (λ=214 nm); (h) Deconvoluted MS (LC-QTOF) spectrum of purified Thz-D-9°N-N-17; (i) Analytical HPLC chromatogram of Thz-D-9°N-N-17-DS crude (λ=214 nm); (j) Analytical HPLC chromatogram of purified Thz-D-9°N-N-17-DS (λ=214 nm); (k) Deconvoluted MS (LC-QTOF) spectrum of purified Thz-D-9°N-N-17-DS; (l) Analytical HPLC chromatogram of D-9°N-N-17 crude (λ=214 nm); (m) Analytical HPLC chromatogram of purified D-9°N-N-17 (λ=214 nm); (n) Deconvoluted MS (LC-QTOF) spectrum of purified D-9°N-N-17; Analytical Method: Column: Welch XB-C4, 4.6x250mm, 5μm. Buffers: A: 0.1%TFA/$H_2O$, B: 0.1%TFA/$CH_3CN$. Gradient: 20 to 70% of Buffer B over 50min; Purification Method: Buffers: A: 0.1%TFA/$H_2O$, B: 0.1%TFA/$CH_3CN$. Gradient: 20 to 70% of Buffer B over 50min. Column: Welch Ultisil XB-C4 21.2x250mm, 5um, 300Å

**(a) D-9°N-N-18:**

**HHHHHH**MILDTDYITENGKPVIRVFKKENGEFKIEYDRTFEPYFYALLKDDSAIEDVKKVTAKRHGTVVKVKRAEKVQKKFL
GRPVEVWKLYFNHPQDVPAIRDRIRAHPAVVDIYEYDIPFAKRYLIDKGLVPMEGDEELTMLAFAIATLYHEGEEFGTGPILMI
SYADGSEARVATWKKVDLPYVDVVSTEKEMVKRFLRVVREKDPDVLITYNGDNFDFAYLKKRC(Acm)EELGVKFTLGRDG
SEPKIQRMGDRFAVEVKGRVHFDLYPVARRTLNLPTYTLCAVYEAVFGKPKEKVYAEEIAQAWESGEGLERVARYSMEDAG
VTYELGREFFPMEAQLSRLIGQSLWDVSRSSTGNLVEWFLLRKAYKRNELAPNKPDERELARRRGGYAGGYVKEPERGLW
DNVVYLDFRSLVPSIIITHNVSPDTLAREGC(Acm)KEYDVAPEVGHKFC(Acm)KDFPGFIPSLLGDLLEERQKIKRK-OH

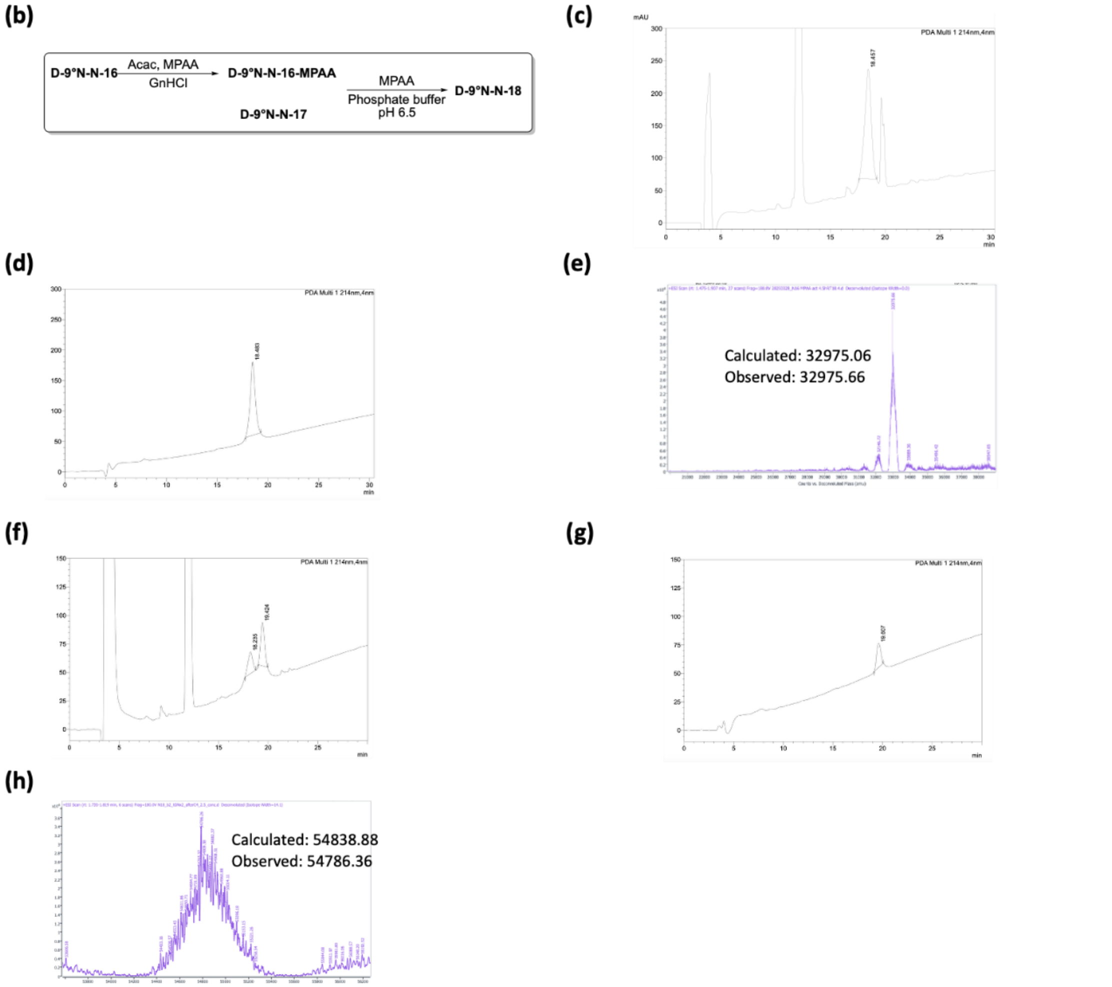

**Supplementary Figure 19.** (a) Peptide sequence of D-9°N-N-18; (b) Reaction Scheme to obtain D-9°N-N-18; (c) Analytical HPLC chromatogram of D-9°N-N-16 thioesterification crude (λ=214 nm); (d) Analytical HPLC chromatogram of purified D-9°N-N-16-MPAA (λ=214 nm); (e) Deconvoluted MS (LC-QTOF) spectrum of purified D-9°N-N-16-MPAA; (f) Analytical HPLC chromatogram of D-9°N-N-18 crude (λ=214 nm); (g) Analytical HPLC chromatogram of purified D-9°N-N-18 (λ=214 nm); (h) Deconvoluted MS (LC-QTOF) spectrum of purified D-9°N-N-18; Analytical Method: Column: Welch XB-C4, 4.6x250mm, 5μm. Buffers: A: 0.1%TFA/$H_2O$, B: 0.1%TFA/$CH_3CN$. Gradient: 20 to 70% of Buffer B over 50min; Purification Method: Buffers: A:

0.1%TFA/$H_2O$, B: 0.1%TFA/$CH_3CN$. Gradient: 30 to 70% of Buffer B over 80 min. Column: Welch Ultisil XB-C4 21.2x250mm, 5um, 300Å

**(a) D-9°N-N-19:**

**HHHHHH**MILDTDYITENGKPVIRVFKKENGEFKIEYDRTFEPYFYALLKDDSAIEDVKKVTAKRHGTVVKVKRAEKVQKKFL
GRPVEVWKLYFNHPQDVPAIRDRIRAHPAVVDIYEYDIPFAKRYLIDKGLVPMEGDEELTMLAFAIATLYHEGEEFGTGPILMI
SYADGSEARVATWKKVDLPYVDVVSTEKEMVKRFLRVVREKDPDVLITYNGDNFDFAYLKKRC(Acm)EELGVKFTLGRDG
SEPKIQRMGDRFAVEVKGRVHFDLYPVARRTLNLPTYTLAAVYEAVFGKPKEKVYAEEIAQAWESGEGLERVARYSMEDAG
VTYELGREFFPMEAQLSRLIGQSLWDVSRSSTGNLVEWFLLRKAYKRNELAPNKPDERELARRRGGYAGGYVKEPERGLW
DNVVYLDFRSLVPSIIITHNVSPDTLAREGC(Acm)KEYDVAPEVGHKFC(Acm)KDFPGFIPSLLGDLLEERQKIKRK-OH

**(b)**

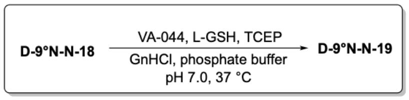

**(c)**

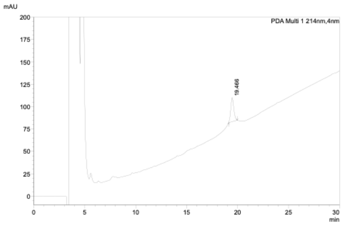

**(d)**

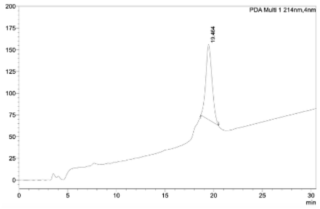

**(e)**

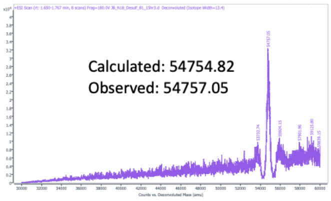

**Supplementary Figure 20.** (a) Peptide sequence of D-9°N-N-19; (b) Reaction Scheme to obtain D-9°N-N-19; (c) Analytical HPLC chromatogram of D-9°N-N-19 crude (λ=214 nm); (d) Analytical HPLC chromatogram of purified D-9°N-N-19 (λ=214 nm); (e) Deconvoluted MS (LC-QTOF) spectrum of purified D-9°N-N-19; Analytical Method: Column: Welch XB-C4, 4.6x250mm, 5µm. Buffers: A: 0.1%TFA/$H_2O$, B: 0.1%TFA/$CH_3CN$. Gradient: 20 to 70% of Buffer B over 50min; Purification Method: Buffers: A: 0.1%TFA/$H_2O$, B: 0.1%TFA/$CH_3CN$. Gradient: 30 to 70% of Buffer B over 30 min. Column: Welch Ultisil XB-C4 21.2x250mm, 5um, 300Å

**(a) D-9°N-N-20:**

**HHHHHH**MILDTDYITENGKPVIRVFKKENGEFKIEYDRTFEPYFYALLKDDSAIEDVKKVTAKRHGTVVKVKRAEKVQKKFL
GRPVEVWKLYFNHPQDVPAIRDRIRAHPAVVDIYEYDIPFAKRYLIDKGLVPMEGDEELTMLAFAIATLYHEGEEFGTGPILMI
SYADGSEARVATWKKVDLPYVDVVSTEKEMVKRFLRVVREKDPDVLITYNGDNFDFAYLKKRCEELGVKFTLGRDGSEPKI
QRMGDRFAVEVKGRVHFDLYPVARRTLNLPTYTLAAVYEAVFGKPKEKVYAEEIAQAWESGEGLERVARYSMEDAGVTYE
LGREFFPMEAQLSRLIGQSLWDVSRSSTGNLVEWFLLRKAYKRNELAPNKPDERELARRRGGYAGGYVKEPERGLWDNVV
YLDFRSLVPSIIITHNVSPDTLAREGCKEYDVAPEVGHKFCKDFPGFIPSLLGDLLEERQKIKRK-OH

**(b)**

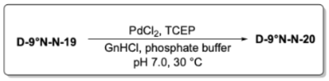

**(c)**

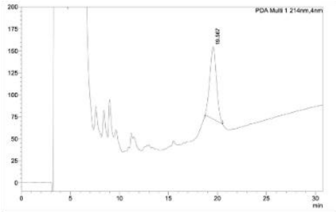

**(d)**

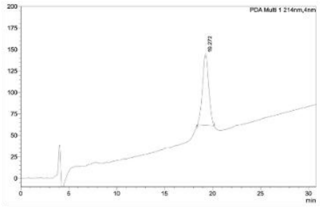

**(e)**

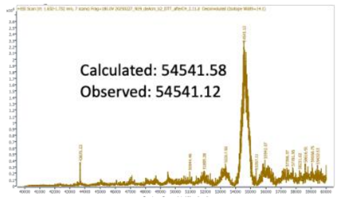

**Supplementary Figure 21.** (a) Peptide sequence of D-9°N-N-20; (b) Reaction Scheme to obtain D-9°N-N-20; (c) Analytical HPLC chromatogram of D-9°N-N-20 crude (λ=214 nm); (d) Analytical HPLC chromatogram of purified D-9°N-N-20 (λ=214 nm); (e) Deconvoluted MS (LC-QTOF) spectrum of purified D-9°N-N-20; Analytical Method: Column: Welch XB-C4, 4.6x250mm, 5µm. Buffers: A: 0.1%TFA/$H_2O$, B: 0.1%TFA/$CH_3CN$. Gradient: 20 to 70% of Buffer B over 50min; Purification Method: Buffers: A: 0.1%TFA/$H_2O$, B: 0.1%TFA/$CH_3CN$. Gradient: 30 to 70% of Buffer B over 30 min. Column: Welch Ultisil XB-C4 21.2x250mm, 5um, 300Å

**(a) D-9°N-C-1:**
MKATVDPLEKKLLDYRQRLIKILANSFYGYYGY-$NHNH_2$

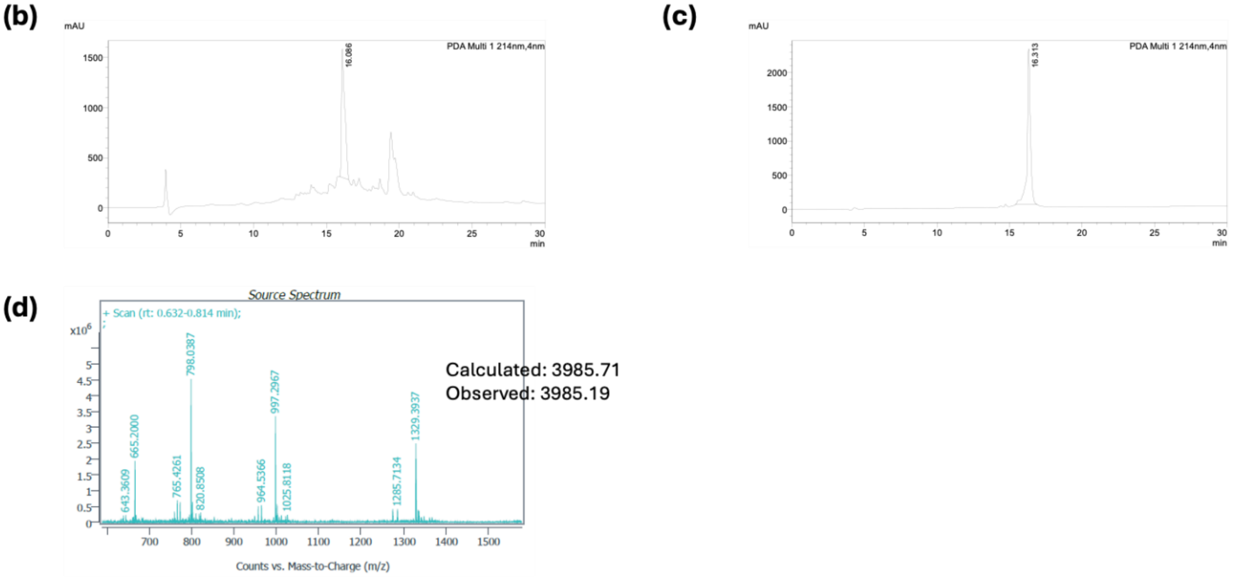

**Supplementary Figure 22.** (a) Sequence of D-9°N-C-1; (b) Analytical HPLC chromatogram of D-9°N-C-1 crude (λ=214 nm); (c) Analytical HPLC chromatogram of the purified D-9°N-C-1 (λ=214 nm); (d) Deconvoluted MS (LC-QTOF) spectrum of purified D-9°N-C-1. Analytical Method: Column: Welch XB-C4, 4.6x250mm, 5µm. Buffers: A: 0.1%TFA/$H_2O$, B: 0.1%TFA/$CH_3CN$. Gradient: 20 to 70% of Buffer B over 30min; Purification Method: Buffers: A: 0.1%TFA/$H_2O$, B: 0.1%TFA/$CH_3CN$. Gradient: 20 to 70% of Buffer B over 50min. Column: Agilent Polaris C18-A, 21.2x250mm, 5um, 180Å.

**(a) D-9°N-C-2:**
CKARWYC(Acm)KEC(Acm)AESVTAWGREYIEMVIRELEEKFGFKVLY-$NHNH_2$

**(b)**

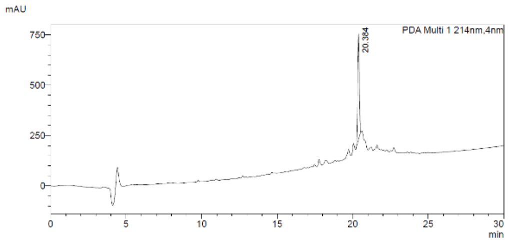

**(c)**

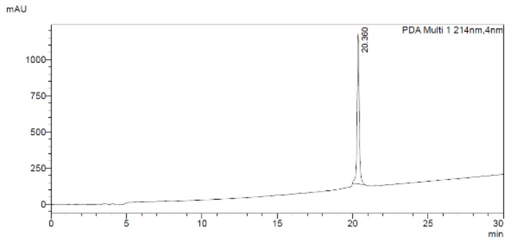

**(d)**

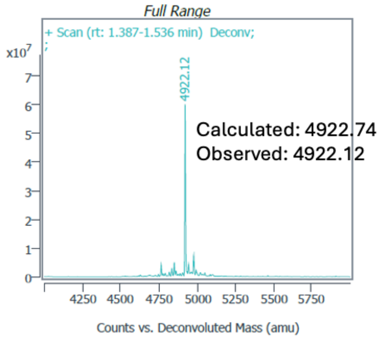

**Supplementary Figure 23.** (a) Sequence of D-9°N-C-2; (b) Analytical HPLC chromatogram of D-9°N-C-2 crude (λ=214 nm); (c) Analytical HPLC chromatogram of the purified D-9°N-C-2 (λ=214 nm); (d) Deconvoluted MS (LC-QTOF) spectrum of purified D-9°N-C-2. Analytical Method: Column: Welch XB-C4, 4.6x250mm, 5µm. Buffers: A: 0.1%TFA/$H_2O$, B: 0.1%TFA/$CH_3CN$. Gradient: 20 to 70% of Buffer B over 30min; Purification Method: Buffers: A: 0.1%TFA/$H_2O$, B: 0.1%TFA/$CH_3CN$. Gradient: 30 to 70% of Buffer B over 40min. Column: Agilent Polaris C18-A, 21.2x250mm, 5um, 180Å.

**(a) D-9°N-C-3:**

CDTDGLHATIPGADAETVKKKAKEFLKYINPKLPGLLELEYEGFYVRGFFVTKKY-NHNH$_2$

**(b)**

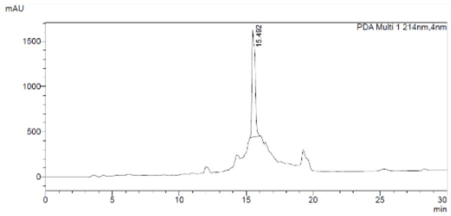

**(c)**

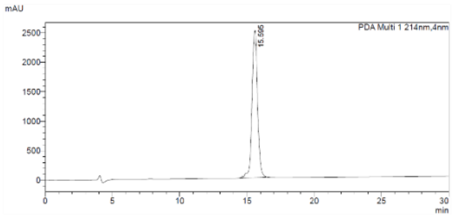

**(d)**

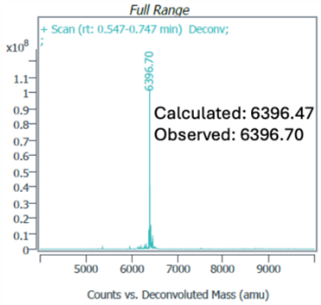

**Supplementary Figure 24.** (a) Sequence of D-9°N-C-3; (b) Analytical HPLC chromatogram of D-9°N-C-3 crude (λ=214 nm); (c) Analytical HPLC chromatogram of the purified D-9°N-C-3 (λ=214 nm); (d) Deconvoluted MS (LC-QTOF) spectrum of purified D-9°N-C-3. Analytical Method: Column: Welch XB-C4, 4.6x250mm, 5µm. Buffers: A: 0.1%TFA/$H_2O$, B: 0.1%TFA/$CH_3CN$. Gradient: 20 to 70% of Buffer B over 30min; Purification Method: Buffers: A: 0.1%TFA/$H_2O$, B: 0.1%TFA/$CH_3CN$. Gradient: 20 to 70% of Buffer B over 50min. Column: Agilent Polaris C18-A, 21.2x250mm, 5um, 180Å.

**(a) D-9°N-C-4:**

(Thz)VVDEEGKITTRGLEVVRRDWSEAAKETQARVLEALLKHGDVEEAVRVVKEVTEKL-$NHNH_2$

**(b)**

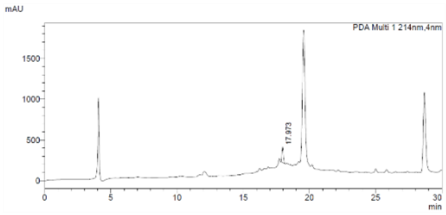

**(c)**

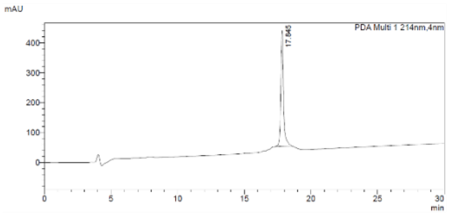

**(d)**

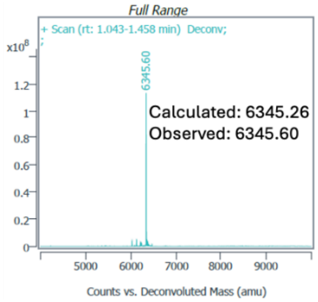

**Supplementary Figure 25.** (a) Sequence of D-9°N-C-4; (b) Analytical HPLC chromatogram of D-9°N-C-4 crude (λ=214 nm); (c) Analytical HPLC chromatogram of the purified D-9°N-C-4 (λ=214 nm); (d) Deconvoluted MS (LC-QTOF) spectrum of purified D-9°N-C-4. Analytical Method: Column: Welch XB-C4, 4.6x250mm, 5µm. Buffers: A: 0.1%TFA/$H_2O$, B: 0.1%TFA/$CH_3CN$. Gradient: 20 to 70% of Buffer B over 30min; Purification Method: Buffers: A: 0.1%TFA/$H_2O$, B: 0.1%TFA/$CH_3CN$. Gradient: 30 to 70% of Buffer B over 40min. Column: Agilent Polaris C18-A, 21.2x250mm, 5um, 180Å.

**(a) D-9°N-C-5:**

CKYEVPPEKLVIHEQITRDLRDYKATGPHVAVAKRLAARGVKIRPGTVISYIVLKGSGRIGDR-$NHNH_2$

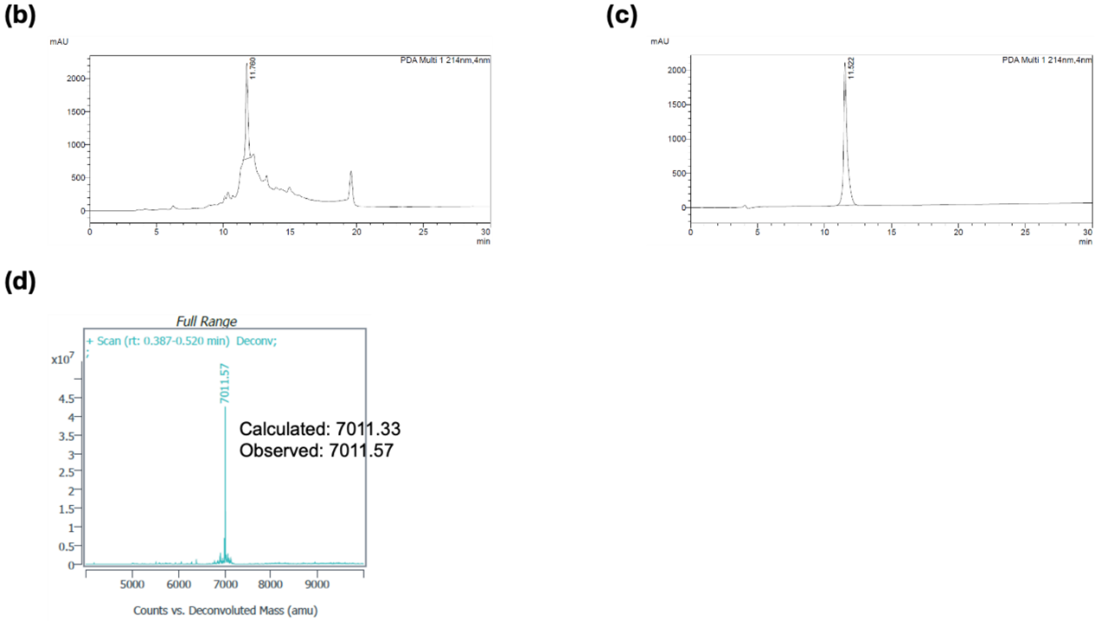

**Supplementary Figure 26.** (a) Sequence of D-9°N-C-5; (b) Analytical HPLC chromatogram of D-9°N-C-5 crude (λ=214 nm); (c) Analytical HPLC chromatogram of the purified D-9°N-C-5 (λ=214 nm); (d) Deconvoluted MS (LC-QTOF) spectrum of purified D-9°N-C-5. Analytical Method: Column: Welch XB-C4, 4.6x250mm, 5µm. Buffers: A: 0.1%TFA/$H_2O$, B: 0.1%TFA/$CH_3CN$. Gradient: 20 to 70% of Buffer B over 30min; Purification Method: Buffers: A: 0.1%TFA/$H_2O$, B: 0.1%TFA/$CH_3CN$. Gradient: 20 to 50% of Buffer B over 30min. Column: Agilent Polaris C18-A, 21.2x250mm, 5um, 180Å.

**(a) D-9°N-C-6:**

CYPADEFDPTKHRYDAEYYVENQVLPAVERVLKAFGYRKEDLRYQKTKQVGLGAWLKVKGKK –OH

**(b)**

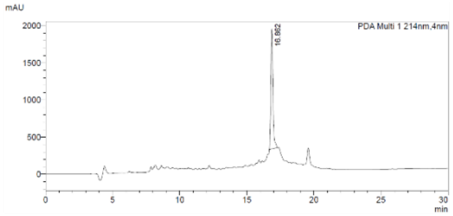

**(c)**

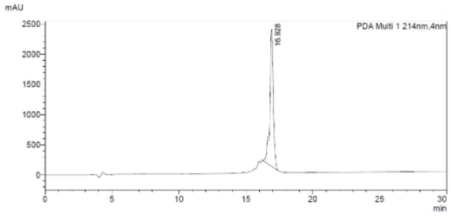

**(d)**

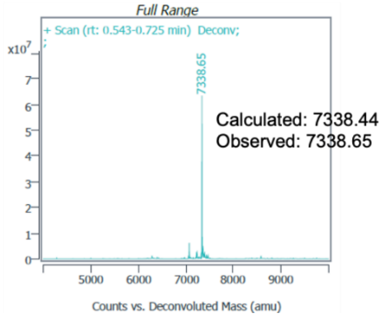

**Supplementary Figure 27.** (a) Sequence of D-9°N-C-6; (b) Analytical HPLC chromatogram of D-9°N-C-6 crude (λ=214 nm); (c) Analytical HPLC chromatogram of the purified D-9°N-C-6 (λ=214 nm); (d) Deconvoluted MS (LC-QTOF) spectrum of purified D-9°N-C-6. Analytical Method: Column: Welch XB-C4, 4.6x250mm, 5μm. Buffers: A: 0.1%TFA/$H_2O$, B: 0.1%TFA/$CH_3CN$. Gradient: 20 to 70% of Buffer B over 30min; Purification Method: Buffers: A: 0.1%TFA/$H_2O$, B: 0.1%TFA/$CH_3CN$. Gradient: 30 to 70% of Buffer B over 40min. Column: Agilent Polaris C18-A, 21.2x250mm, 5um, 180Å.

**(a) D-9°N-C-7:**

MKATVDPLEKKLLDYRQRLIKILANSFYGYYGYCKARWYC(Acm)KEC(Acm)AESVTAWGREYI EMVIRELE EKFGFKVLY-NHNH2

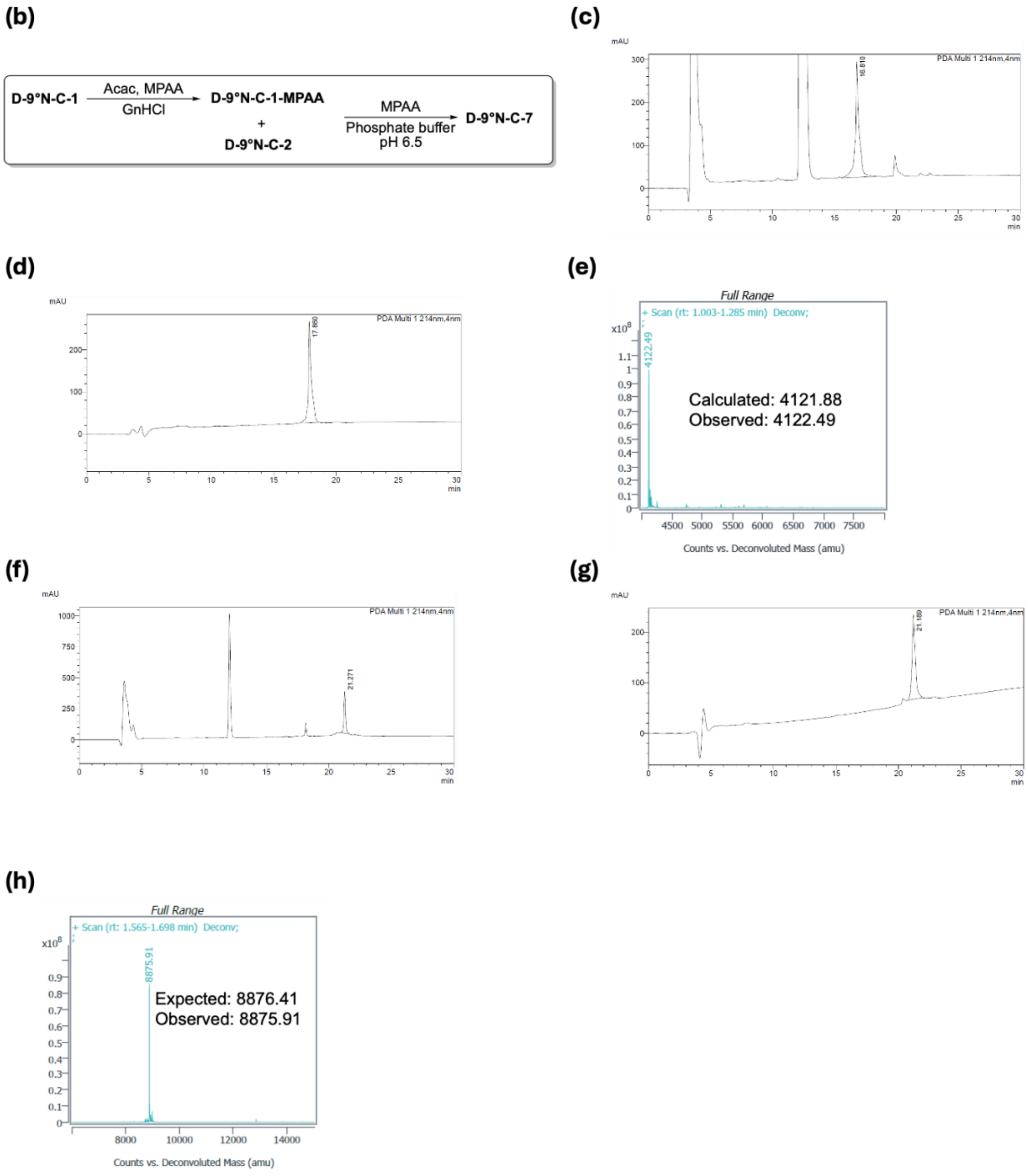

**Supplementary Figure 28.** (a) Peptide sequence of D-9°N-C-7; (b) Reaction Scheme to obtain D-9°N-C-7; (c) Analytical HPLC chromatogram of D-9°N-C-1 thioesterification crude (λ=214 nm); (d) Analytical HPLC chromatogram of purified D-9°N-C-1-MPAA (λ=214 nm); (e) Deconvoluted MS (LC-QTOF) spectrum of D-9°N-C-1-MPAA ; (f) Analytical HPLC chromatogram of D-9°N-C-7 crude (λ=214 nm); (g) Analytical HPLC chromatogram of purified

D-9°N-C-7 (λ=214 nm); (h) Deconvoluted MS (LC-QTOF) spectrum of purified D-9°N-C-7; Analytical Method: Column: Welch XB-C4, 4.6x250mm, 5μm. Buffers: A: 0.1%TFA/$H_2O$, B: 0.1%TFA/$CH_3CN$. Gradient: 20 to 70% of Buffer B over 50min; Purification Method: Buffers: A: 0.1%TFA/$H_2O$, B: 0.1%TFA/$CH_3CN$. Gradient: 30 to 70% of Buffer B over 40min. Column: Welch Ultisil XB-C4 21.2x250mm, 5um, 300Å

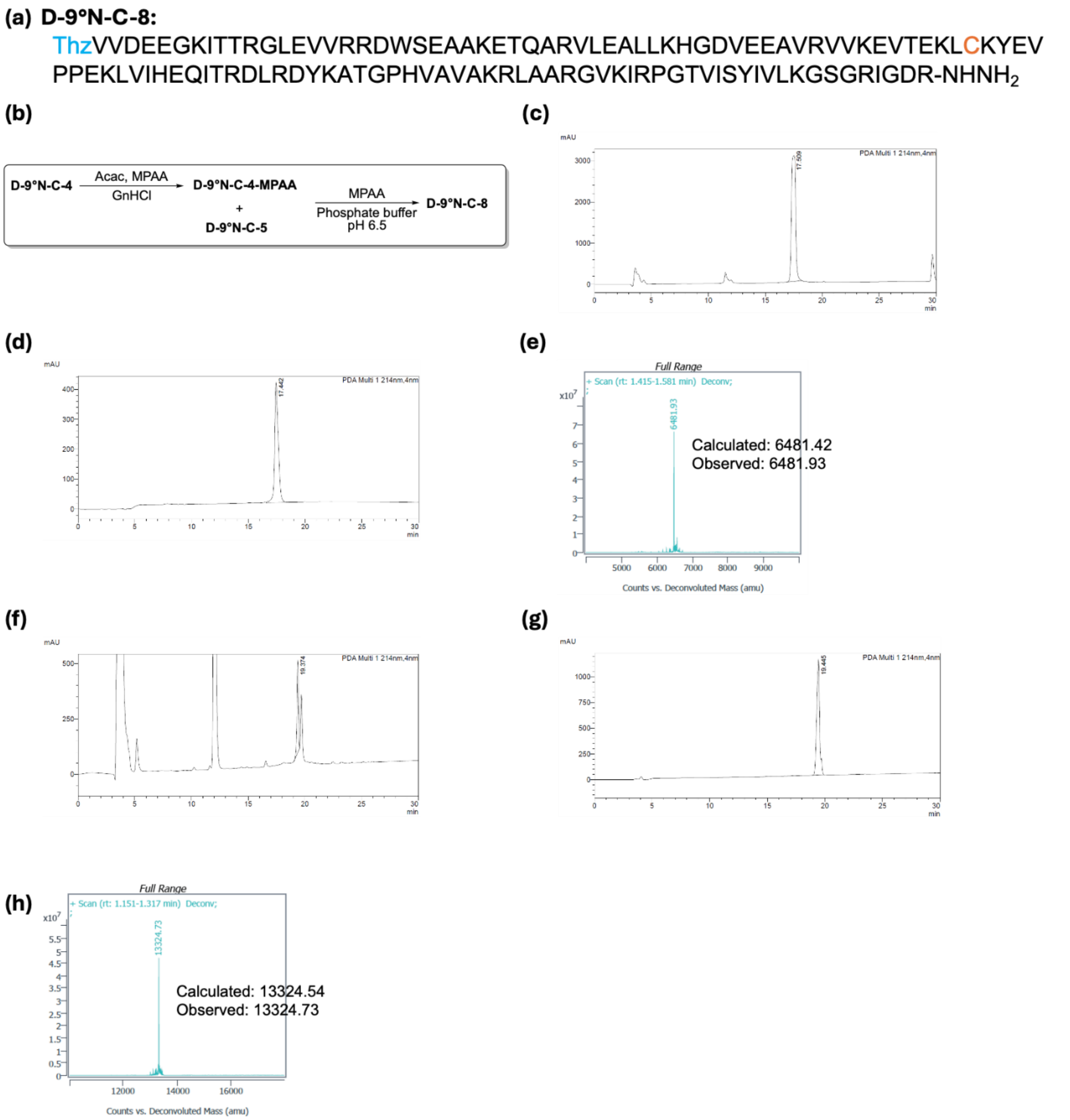

**Supplementary Figure 29.** (a) Peptide sequence of D-9°N-C-8; (b) Reaction Scheme to obtain D-9°N-C-8; (c) Analytical HPLC chromatogram of D-9°N-C-4 thioesterification crude (λ=214 nm); (d) Analytical HPLC chromatogram of purified D-9°N-C-4-MPAA (λ=214 nm); (e) Deconvoluted MS (LC-QTOF) spectrum of D-9°N-C-4-MPAA ; (f) Analytical HPLC chromatogram of D-9°N-C-8 crude (λ=214 nm); (g) Analytical HPLC chromatogram of purified D-9°N-C-8 (λ=214 nm); (h) Deconvoluted MS (LC-QTOF) spectrum of purified D-9°N-C-8; Analytical Method: Column: Welch XB-C4, 4.6x250mm, 5μm. Buffers: A: 0.1%TFA/$H_2O$, B:

0.1%TFA/$CH_3CN$. Gradient: 20 to 70% of Buffer B over 50min; Purification Method: Buffers: A: 0.1%TFA/$H_2O$, B: 0.1%TFA/$CH_3CN$. Gradient: 30 to 70% of Buffer B over 40min. Column: Welch Ultisil XB-C4 21.2x250mm, 5um, 300Å

**(a) D-9°N-C-9:**

MKATVDPLEKKLLDYRQRLIKILANSFYGYYGYCKARWYC(Acm)KEC(Acm)AESVTAWGREYI
EMVIRELEEKFGFKVLYCDTDGLHATIPGADAETVKKKAKEFLKYINPKLPGLLELEYEGFYVRG
FFVTKKKY-$NHNH_2$

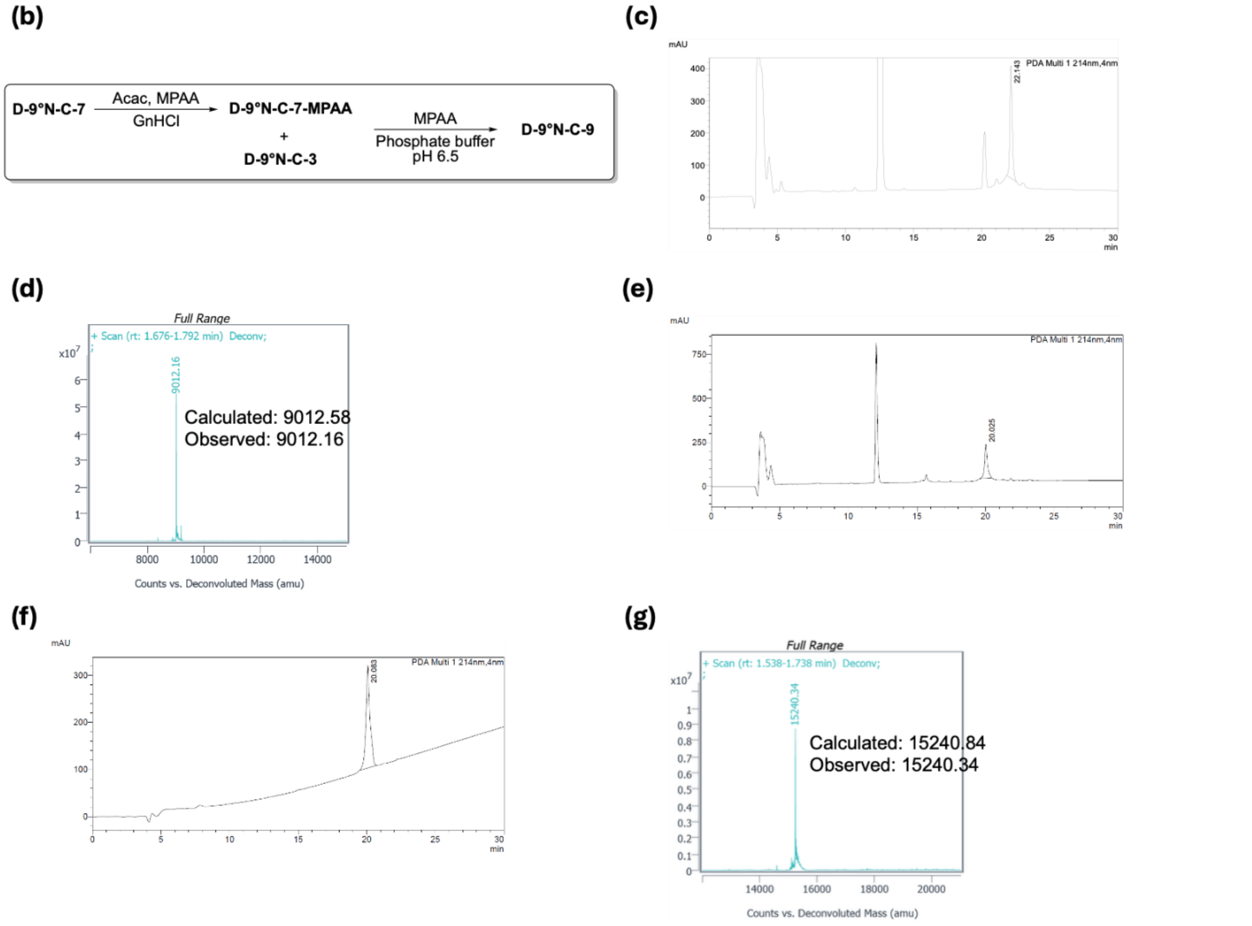

**Supplementary Figure 30.** (a) Peptide sequence of D-9°N-C-9; (b) Reaction Scheme to obtain D-9°N-C-9; (c) Analytical HPLC chromatogram of D-9°N-C-7 thioesterification crude (λ=214 nm); (d) Deconvoluted MS (LC-QTOF) spectrum of D-9°N-C-7-MPAA from fraction collection of analytical HPLC; (e) Analytical HPLC chromatogram of D-9°N-C-9 crude (λ=214 nm); (f) Analytical HPLC chromatogram of purified D-9°N-C-9 (λ=214 nm); (g) Deconvoluted MS (LC-QTOF) spectrum of purified D-9°N-C-9; Analytical Method: Column: Welch XB-C4, 4.6x250mm, 5µm. Buffers: A: 0.1%TFA/$H_2O$, B: 0.1%TFA/$CH_3CN$. Gradient: 20 to 70% of Buffer B over 50min; Purification Method: Buffers: A: 0.1%TFA/$H_2O$, B: 0.1%TFA/$CH_3CN$. Gradient: 30 to 70% of Buffer B over 40min. Column: Welch Ultisil XB-C4 21.2x250mm, 5um, 300Å

(a) **D-9°N-C-10:**

CVVDEEGKITTRGLEVVRRDWSEAAKETQARVLEALLKHGDVEEAVRVVKEVTEKLCKYEVPP
EKLVIHEQITRDLRDYKATGPHVAVAKRLAARGVKIRPGTVISYIVLKGSGRIGDRCYPADEFDPT
KHRYDAEYYVENQVLPAVERVLKAFGYRKEDLRYQKTKQVGLGAWLKVKGKK–OH

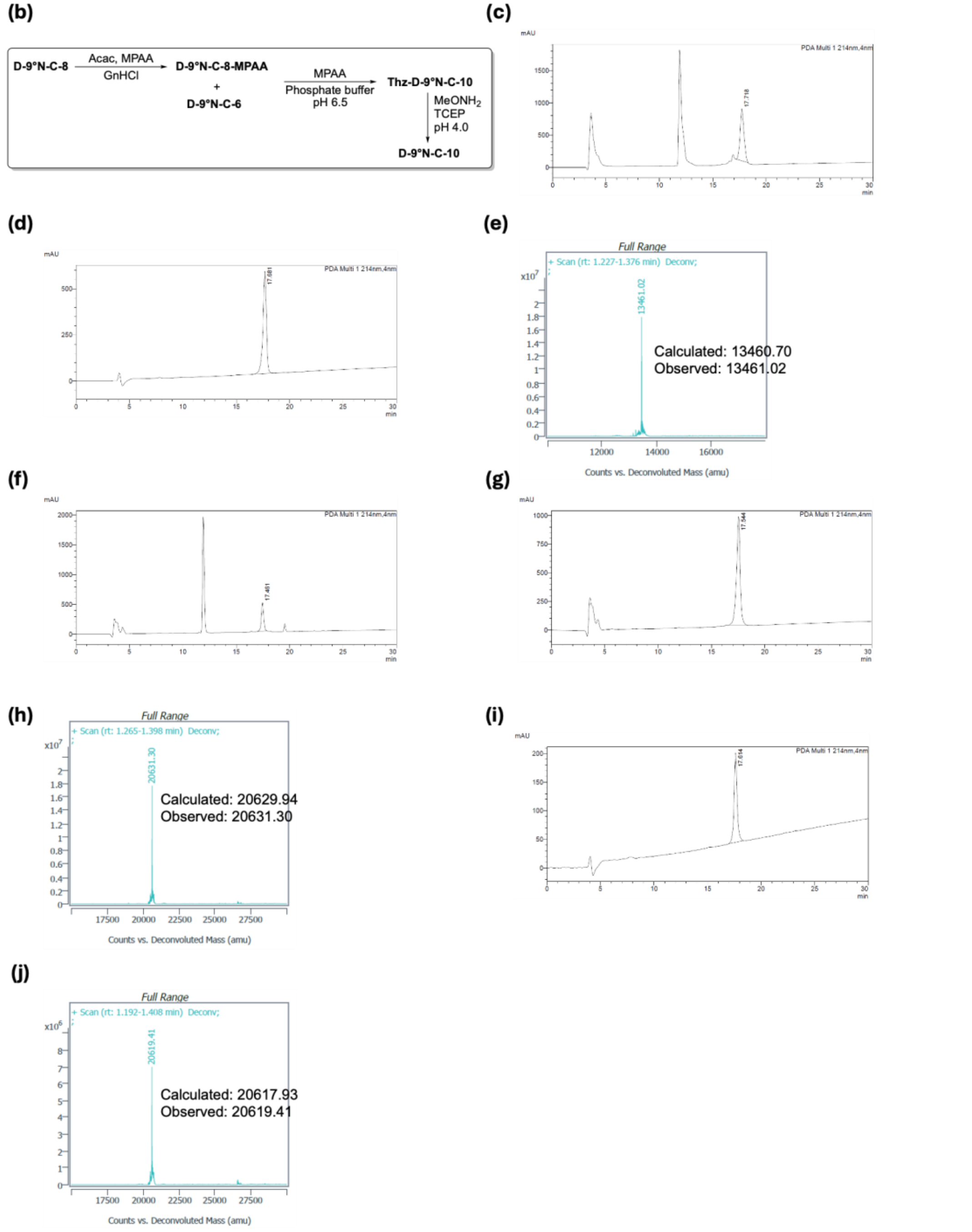

**Supplementary Figure 31.** (a) Peptide sequence of D-9°N-C-10; (b) Reaction Scheme to obtain D-9°N-C-10; (c) Analytical HPLC chromatogram of D-9°N-C-8 thioesterification crude (λ=214 nm); (d) Analytical HPLC chromatogram of purified D-9°N-C-8-MPAA (λ=214 nm); (e)

Deconvoluted MS (LC-QTOF) spectrum of purified D-9°N-C-8-MPAA; (f) Analytical HPLC chromatogram of Thz-D-9°N-C-10 crude (λ=214 nm); (g) Analytical HPLC chromatogram of purified Thz-D-9°N-C-10 (λ=214 nm); (h) Deconvoluted MS (LC-QTOF) spectrum of purified Thz-D-9°N-C-10; (g) Analytical HPLC chromatogram of purified D-9°N-C-10 (λ=214 nm); (h) Deconvoluted MS (LC-QTOF) spectrum of purified D-9°N-C-10; Analytical Method: Column: Welch XB-C4, 4.6x250mm, 5μm. Buffers: A: 0.1%TFA/$H_2O$, B: 0.1%TFA/$CH_3CN$. Gradient: 20 to 70% of Buffer B over 50min; Purification Method: Buffers: A: 0.1%TFA/$H_2O$, B: 0.1%TFA/$CH_3CN$. Gradient: 30 to 70% of Buffer B over 40min. Column: Welch Ultisil XB-C4 21.2x250mm, 5um, 300Å

**(a) D-9°N-C-11:**

MKATVDPLEKKLLDYRQRLIKILANSFYGYYGYCKARWYC(Acm)KEC(Acm)AESVTAWGREYIE
MVIRELEEKFGFKVLYCDTDGLHATIPGADAETVKKKAKEFLKYINPKLPGLLELEYEGFYVRGF
FVTKKKYCVVDEEGKITTRGLEVVRRDWSEAAKETQARVLEALLKHGDVEEAVRVVKEVTEKL
CKYEVPPEKLVIHEQITRDLRDYKATGPHVAVAKRLAARGVKIRPGTVISYIVLKGSGRIGDRCYP
ADEFDPTKHRYDAEYYVENQVLPAVERVLKAFGYRKEDLRYQKTKQVGLGAWLKVKGKK–OH

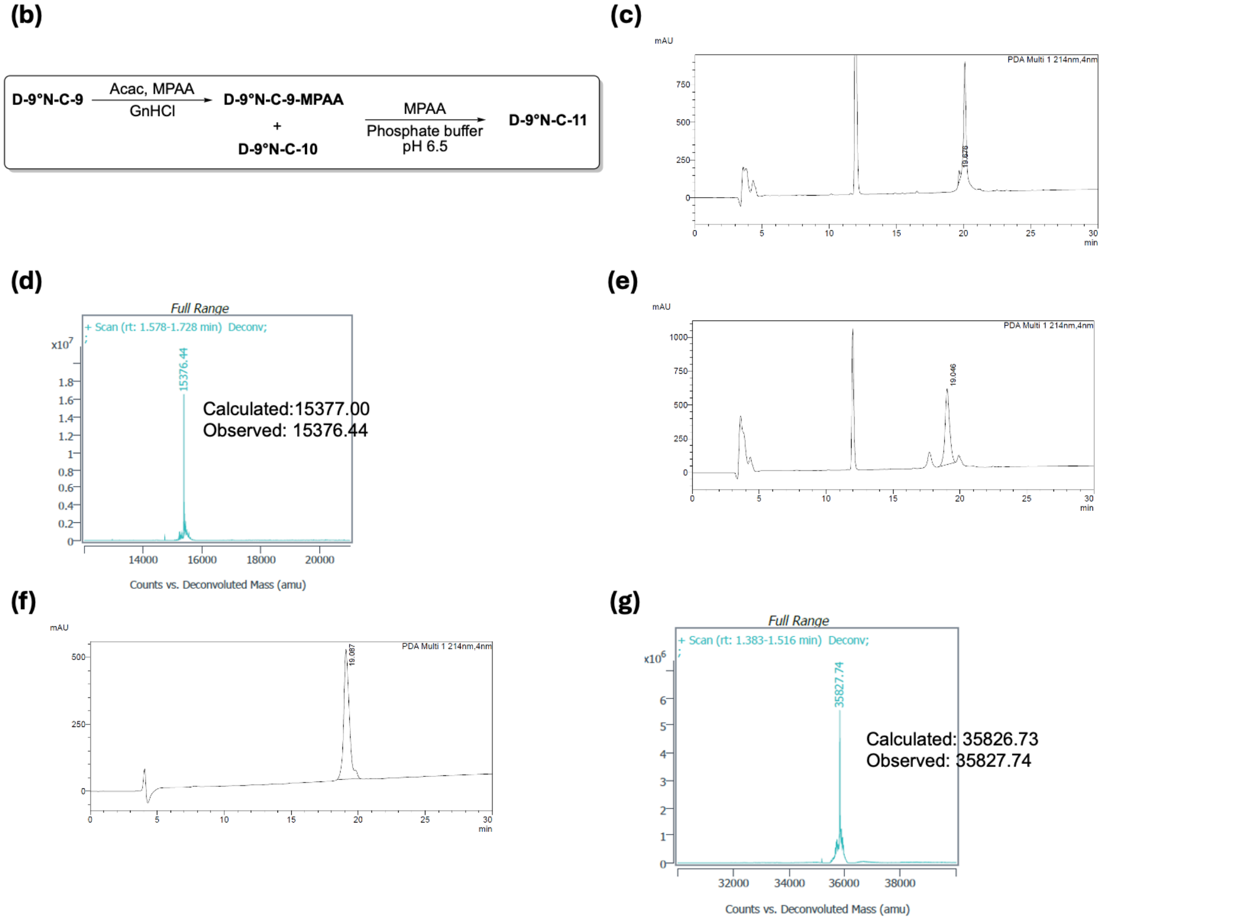

**Supplementary Figure 32.** (a) Peptide sequence of D-9°N-C-11; (b) Reaction Scheme to obtain D-9°N-C-11; (c) Analytical HPLC chromatogram of D-9°N-C-9 thioesterification crude (λ=214 nm); (d) Deconvoluted MS (LC-QTOF) spectrum of D-9°N-C-9-MPAA from fraction collection of analytical HPLC; (e) Analytical HPLC chromatogram of D-9°N-C-11 crude (λ=214 nm); (f) Analytical HPLC chromatogram of purified D-9°N-C-11 (λ=214 nm); (g) Deconvoluted MS (LC-QTOF) spectrum of purified D-9°N-C-11; Analytical Method: Column: Welch XB-C4, 4.6x250mm, 5μm. Buffers: A: 0.1%TFA/$H_2O$, B: 0.1%TFA/$CH_3CN$. Gradient: 20 to 70% of Buffer B over 50min; Purification Method: Buffers: A: 0.1%TFA/$H_2O$, B: 0.1%TFA/$CH_3CN$.

Gradient: 30 to 70% of Buffer B over 40min. Column: Welch Ultisil XB-C4 21.2x250mm, 5um, 300Å

**(a) D-9°N-C-12:**

MKATVDPLEKKLLDYRQRLIKILANSFYGYYGYAKARWYC(Acm)KEC(Acm)AESVTAWGREYIE
MVIRELEEKFGFKVLYADTDGLHATIPGADAETVKKKAKEFLKYINPKLPGLLELEYEGFYVRGFF
VTKKKYAVVDEEGKITTRGLEVVRRDWSEAAKETQARVLEALLKHGDVEEAVRVVKEVTEKLAK
YEVPPEKLVIHEQITRDLRDYKATGPHVAVAKRLAARGVKIRPGTVISYIVLKGSGRIGDRAYPAD
EFDPTKHRYDAEYYVENQVLPAVERVLKAFGYRKEDLRYQKTKQVGLGAWLKVKGKK–OH

**(b)**

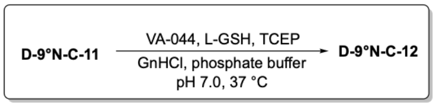

**(c)**

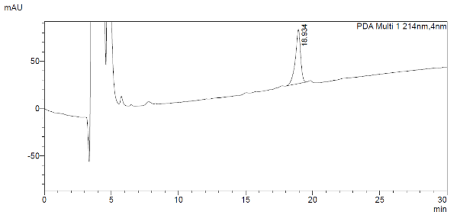

**(d)**

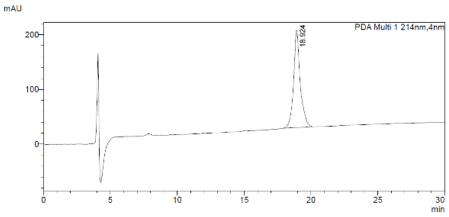

**(e)**

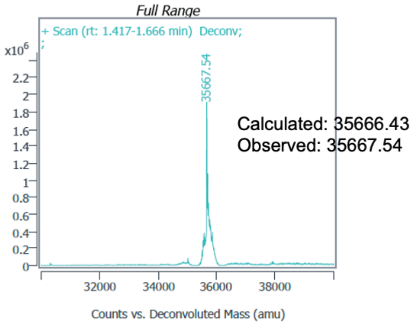

**Supplementary Figure 33.** (a) Peptide sequence of D-9°N-C-12; (b) Reaction Scheme to obtain D-9°N-C-12; (c) Analytical HPLC chromatogram of D-9°N-C-11 desulfurization crude (λ=214 nm); (d) Analytical HPLC chromatogram of purified D-9°N-C-12 (λ=214 nm); (e) Deconvoluted MS (LC-QTOF) spectrum of purified D-9°N-C-12; Analytical Method: Column: Welch XB-C4, 4.6x250mm, 5µm. Buffers: A: 0.1%TFA/$H_2O$, B: 0.1%TFA/$CH_3CN$. Gradient: 20 to 70% of Buffer B over 50min; Purification Method: Buffers: A: 0.1%TFA/$H_2O$, B: 0.1%TFA/$CH_3CN$. Gradient: 35 to 70% of Buffer B over 35min. Column: Welch Ultisil XB-C4 21.2x250mm, 5um, 300Å

**(a) D-9°N-C-13:**

MKATVDPLEKKLLDYRQRLIKILANSFYGYYGYAKARWYCKECAESVTAWGREYIEMVIRELEE
KFGFKVLYADTDGLHATIPGADAETVKKKAKEFLKYINPKLPGLLELEYEGFYVRGFFVTKKKYA
VVDEEGKITTRGLEVVRRDWSEAAKETQARVLEALLKHGDVEEAVRVVKEVTEKLAKYEVPPE
KLVIHEQITRDLRDYKATGPHVAVAKRLAARGVKIRPGTVISYIVLKGSGRIGDRAYPADEFDPTK
HRYDAEYYVENQVLPAVERVLKAFGYRKEDLRYQKTKQVGLGAWLKVKGKK–OH

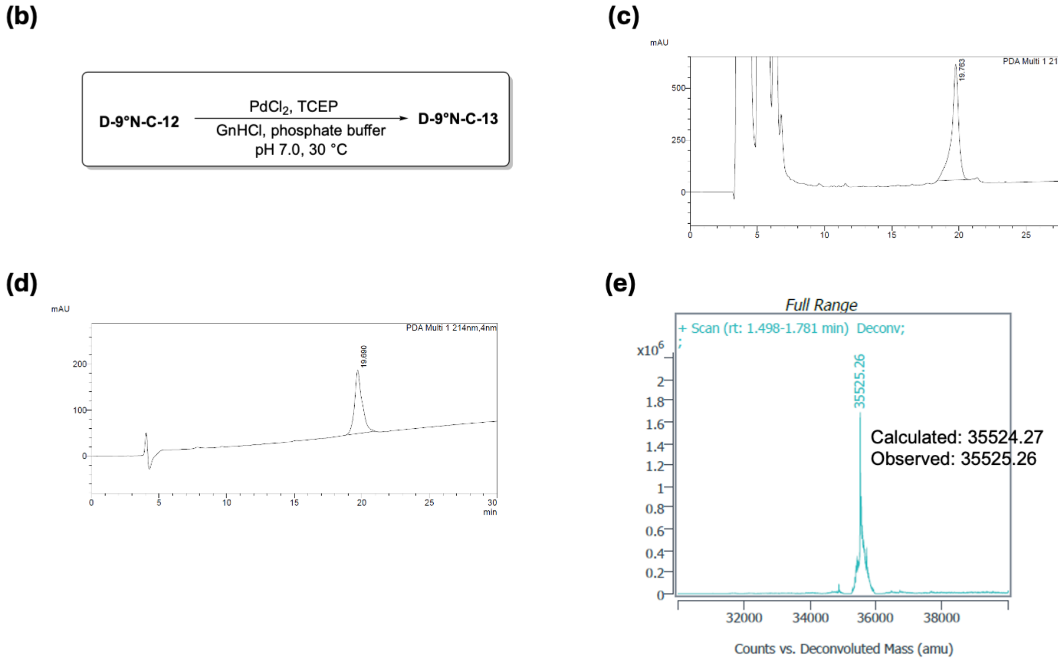

**Supplementary Figure 34.** (a) Peptide sequence of D-9°N-C-13; (b) Reaction Scheme to obtain D-9°N-C-13; (c) Analytical HPLC chromatogram of D-9°N-C-12 Acm deprotection crude (λ=214 nm); (d) Analytical HPLC chromatogram of purified D-9°N-C-13 (λ=214 nm); (e) Deconvoluted MS (LC-QTOF) spectrum of purified D-9°N-C-13; Analytical Method: Column: Welch XB-C4, 4.6x250mm, 5µm. Buffers: A: 0.1%TFA/$H_2O$, B: 0.1%TFA/$CH_3CN$. Gradient: 20 to 70% of Buffer B over 50min; Purification Method: Buffers: A: 0.1%TFA/$H_2O$, B: 0.1%TFA/$CH_3CN$. Gradient: 35 to 70% of Buffer B over 35min. Column: Welch Ultisil XB-C4 21.2x250mm, 5um, 300Å

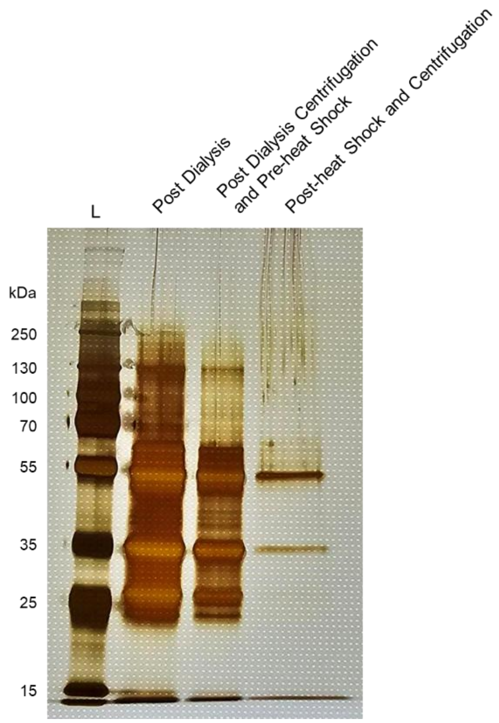

**Supplementary Figure 35.** SDS-PAGE analysis of refolding steps for (D)-9°N polymerase. Aliquots were removed from refolding steps and purification steps to monitor the presence of the N20 and C13. The following amounts were loaded onto the SDS-PAGE: post dialysis day 2 (10 μL), post-centrifugation to remove precipitation (10 μL), and post-heat shock and centrifugation (30 μL).

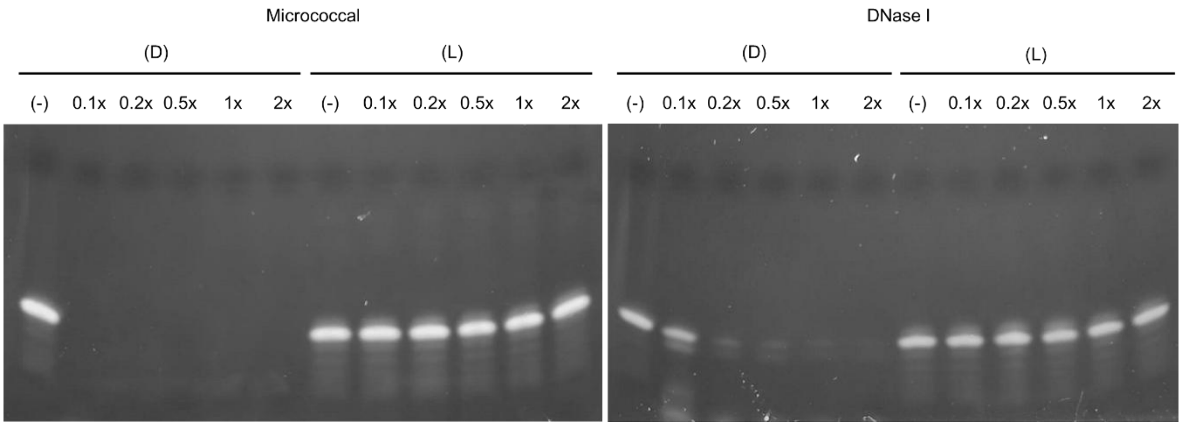

**Supplementary Figure 36.** Titration of nucleases in the presence of (D)- and (L)-form oligonucleotides. NEB defines micrococcal 1x as 12500 gel units/μL. NEB defines DNase I 1x activity as 2 units per reaction. NEB enzymes were diluted at 1:2, 1:5, and 1:10 for both micrococcal and DNase I. Micrococcal has strong DNA degradation ability up to a 10-fold dilution. DNase I has degradation ability up to a 5-fold dilution.

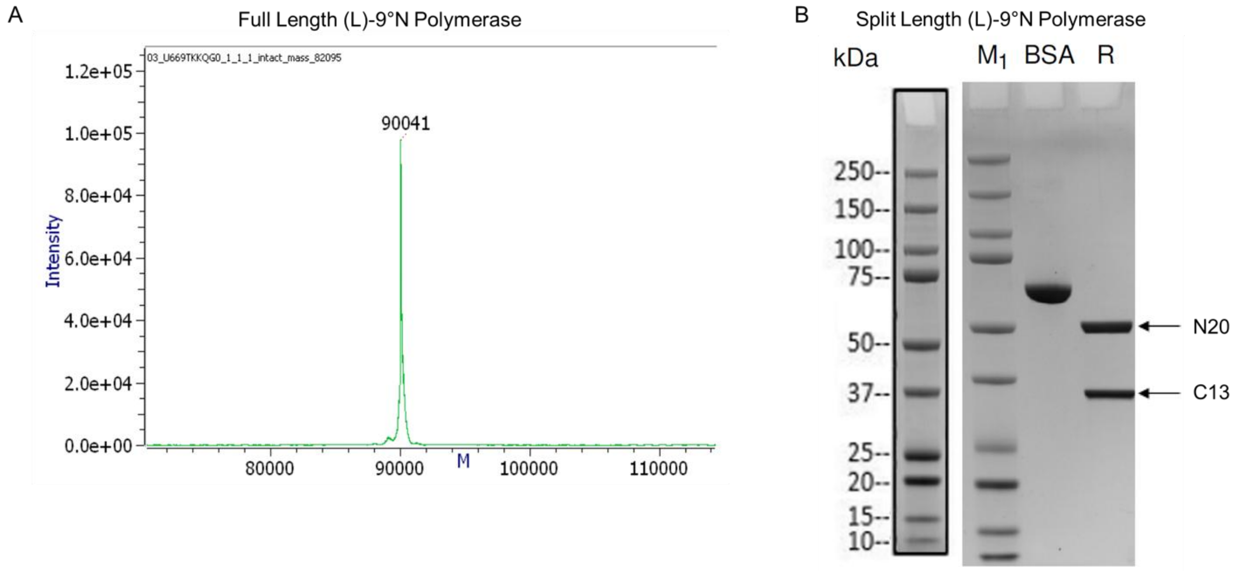

**Supplementary Figure 37.** Verification of recombinantly expressed full length and split (L)-9°N polymerase. LC-MS was performed on the Met-6xHis-tagged-(L)-9°N polymerase (observed m/z: 90041 Da, calculated m/z 90.06 kDa). SDS-PAGE analysis of split Met-6xHis-tagged-(L)-9°N N (calculated mw:54.55 kDa ) and C (calculated mw:35.53 kDa) fragments.

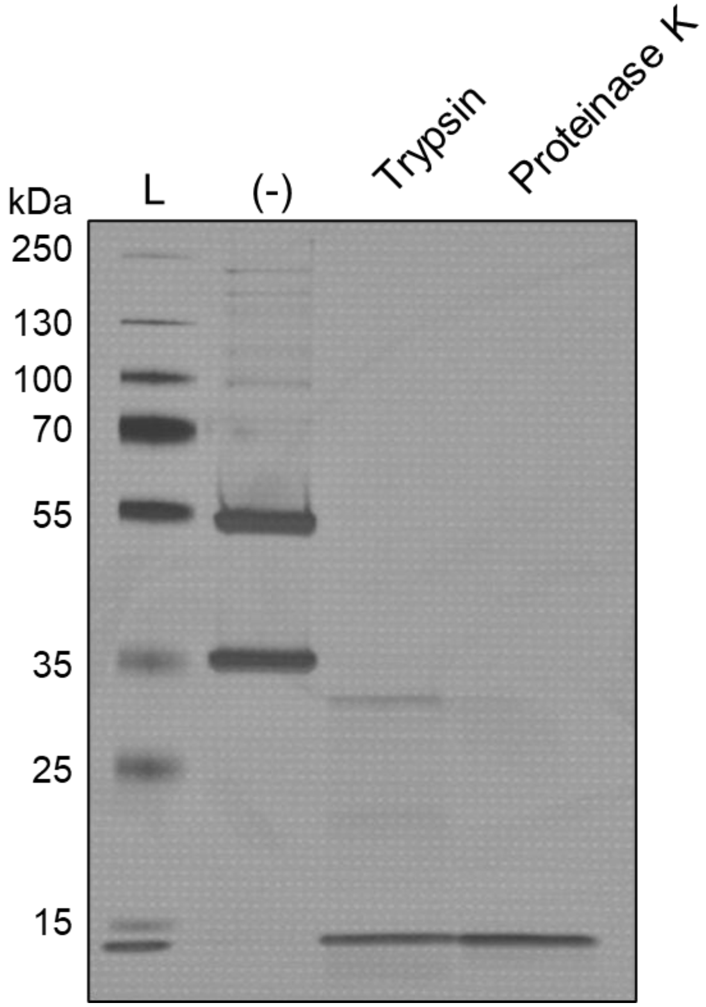

**Supplementary Figure 38**. Protease degradation of recombinantly expressed split (L)-9°N polymerase via SDS-PAGE analysis. Split polymerase was incubated with trypsin and proteinase K. Degradation is observed for the natural (L)-9°N polymerase.

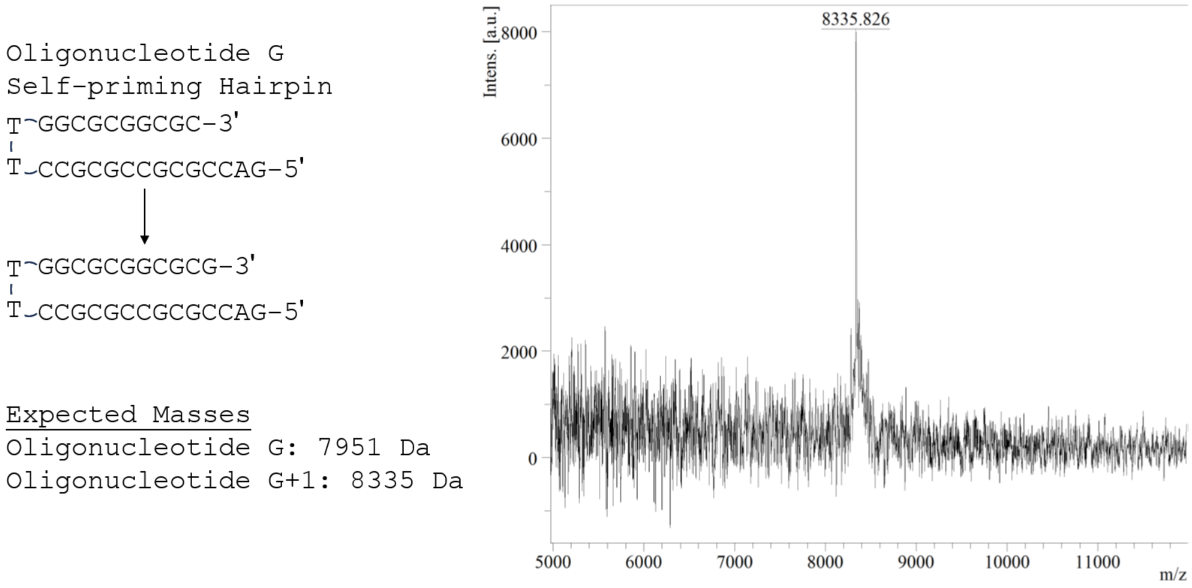

**Supplementary Figure 39.** Single nucleotide incorporation of (L)-3′-*O*-azidomethyl-dGTP across (L)-Oligonucleotide G self-priming template. Reaction was confirmed by MALDI with the expected mass of 8335 Da.

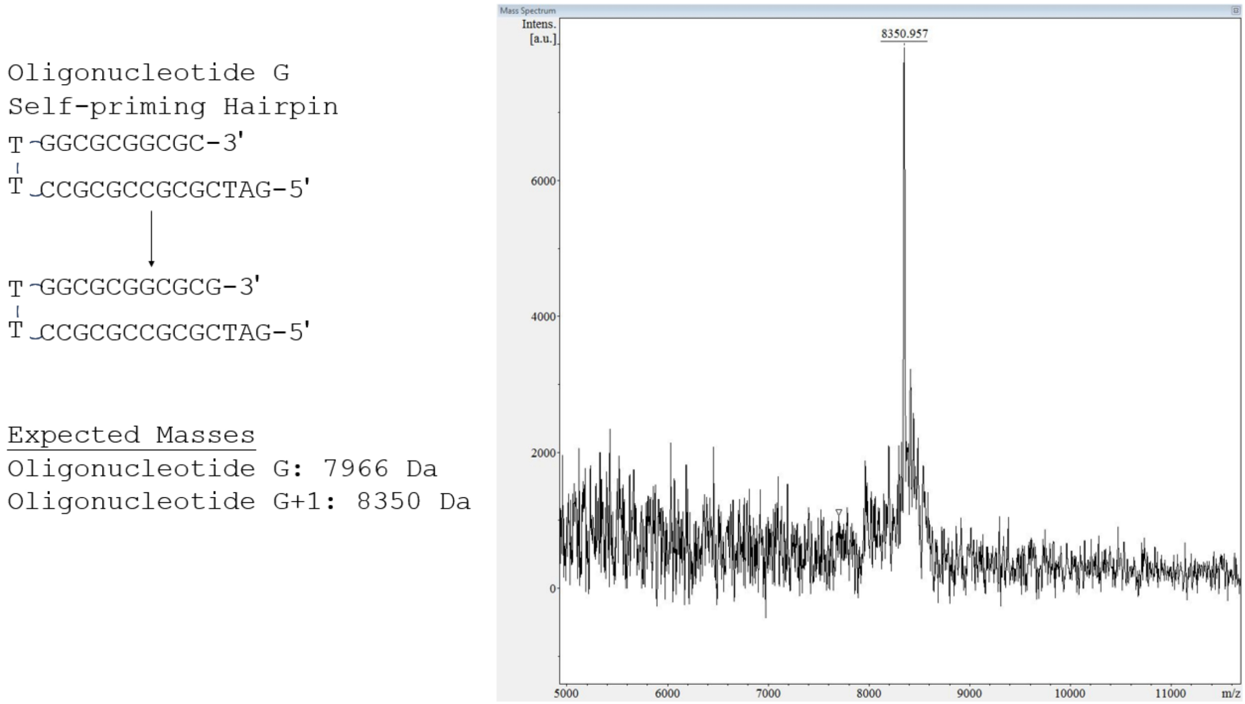

**Supplementary Figure 40.** Single nucleotide incorporation of (D)-3′-*O*-azidomethyl-dGTP across (D)-Oligonucleotide G self-priming template. Reaction was confirmed by MALDI with the expected mass of 8350 Da.

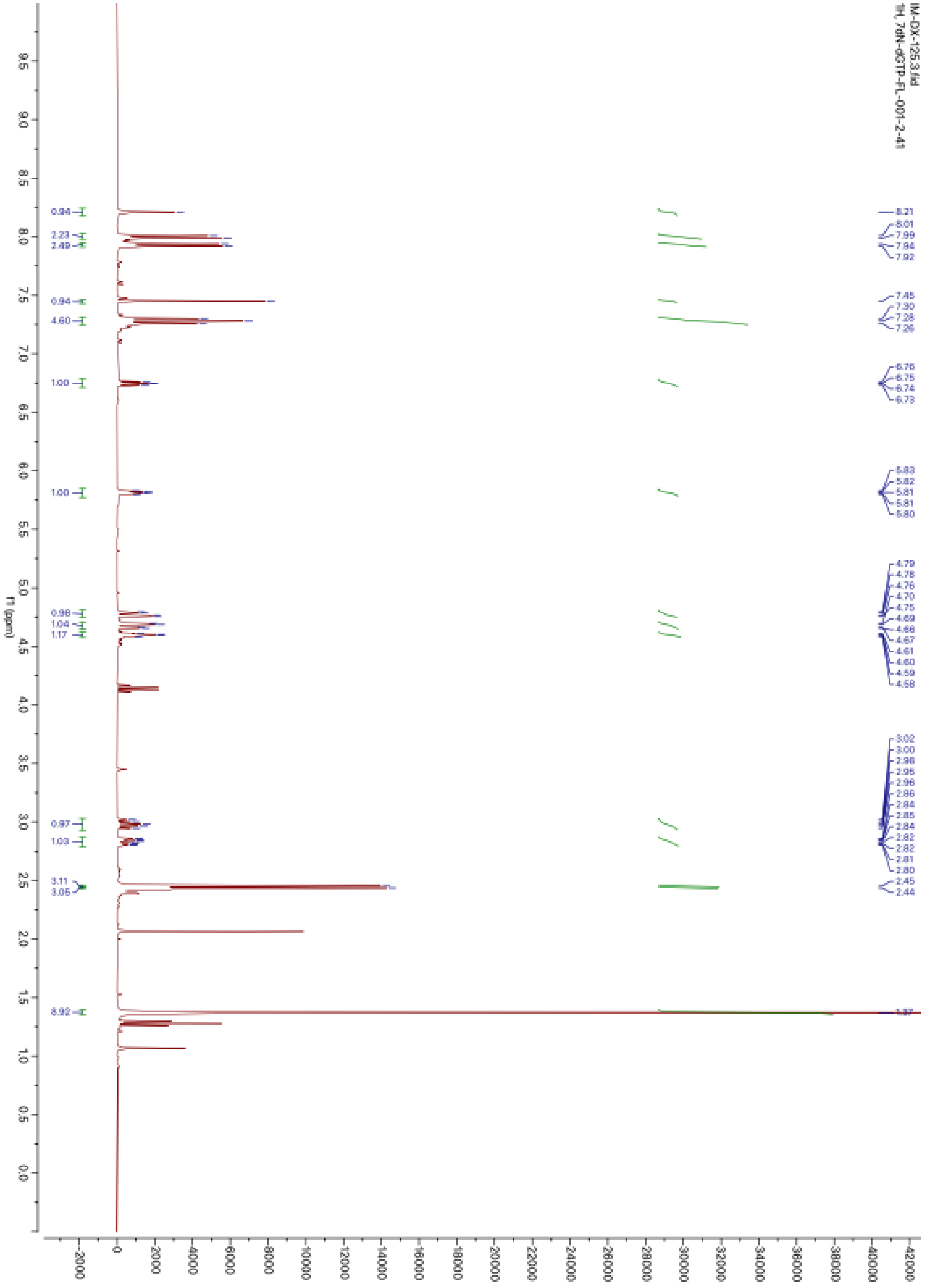

**Supplementary Figure 41.** $^{1}$H NMR of compound **4** in $CDCl_3$

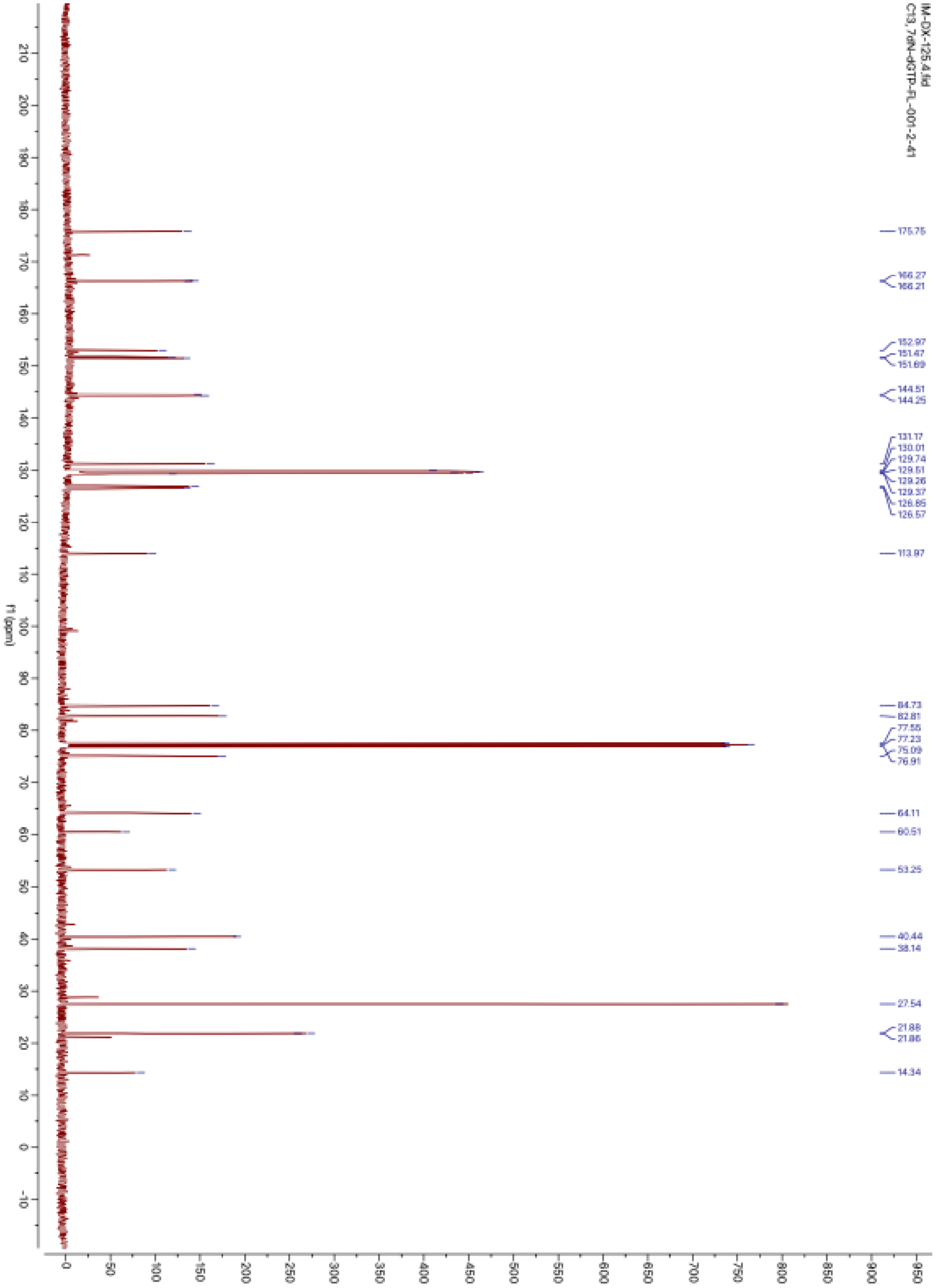

**Supplementary Figure 42.** $^{13}C$ NMR of compound **4** in $CDCl_3$

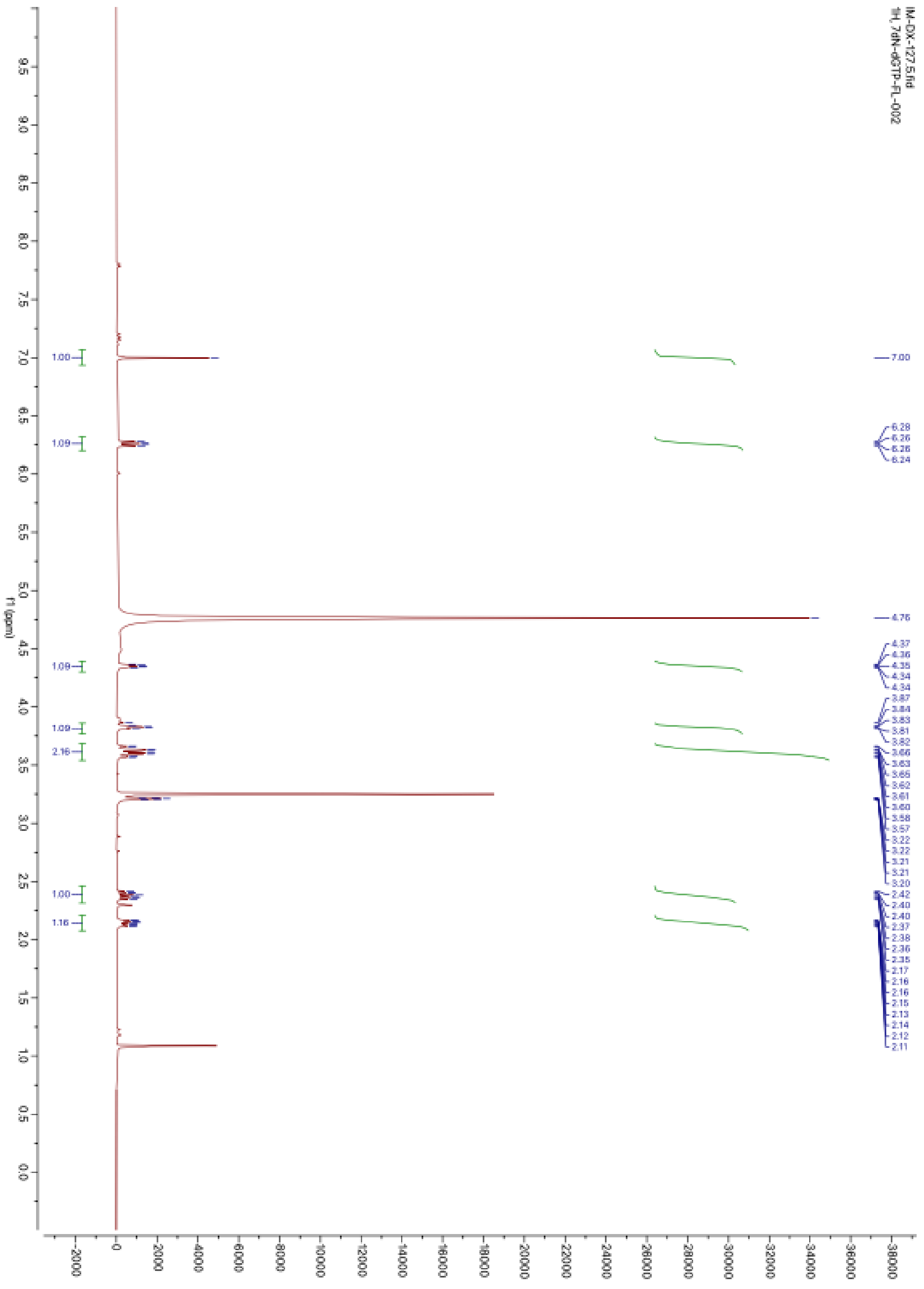

**Supplementary Figure 43.** $^{1}H$ NMR of compound **5** in $CD_3OD$

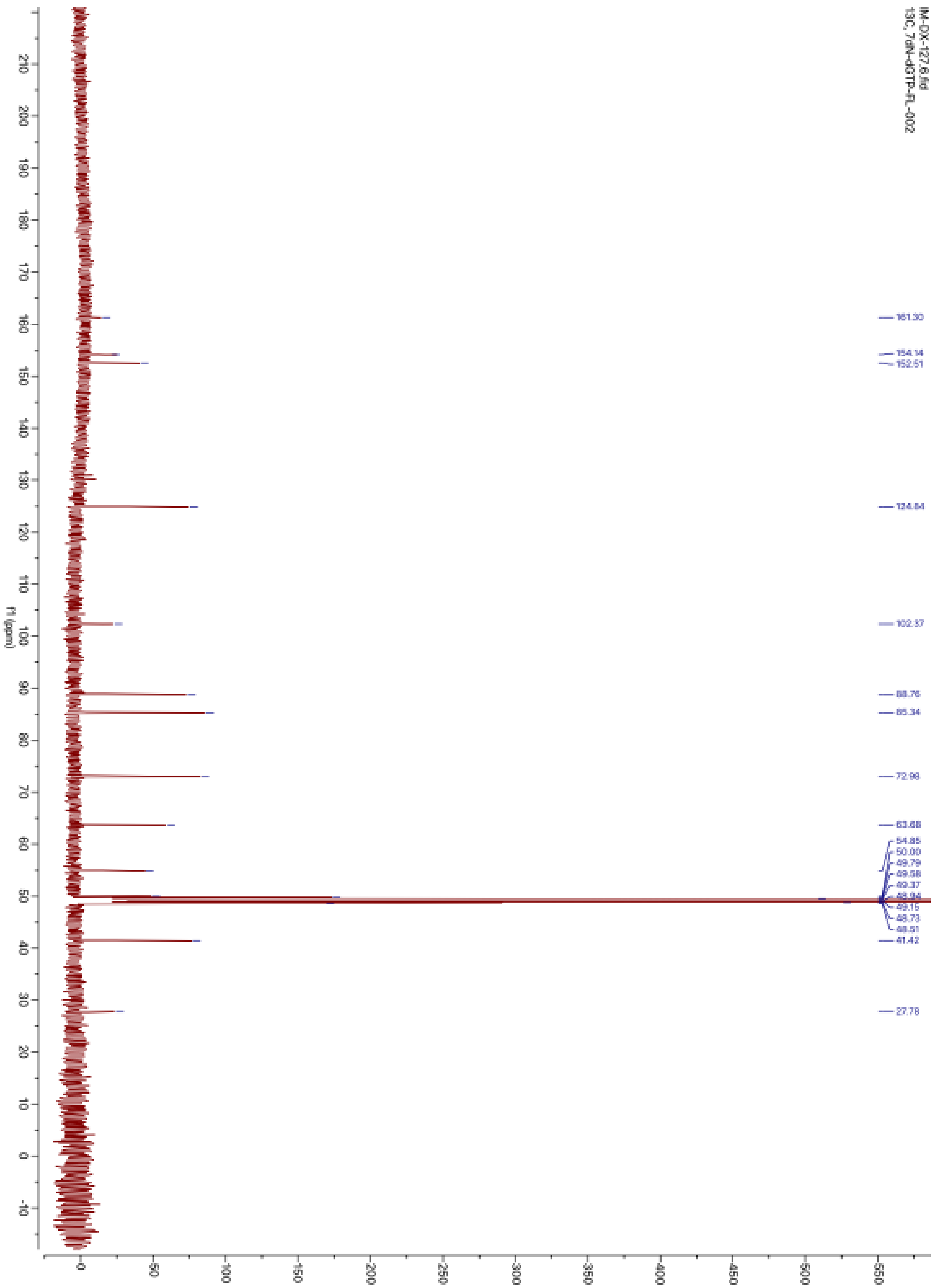

**Supplementary Figure 44.** $^{13}C$ NMR of compound **5** in $CD_3OD$

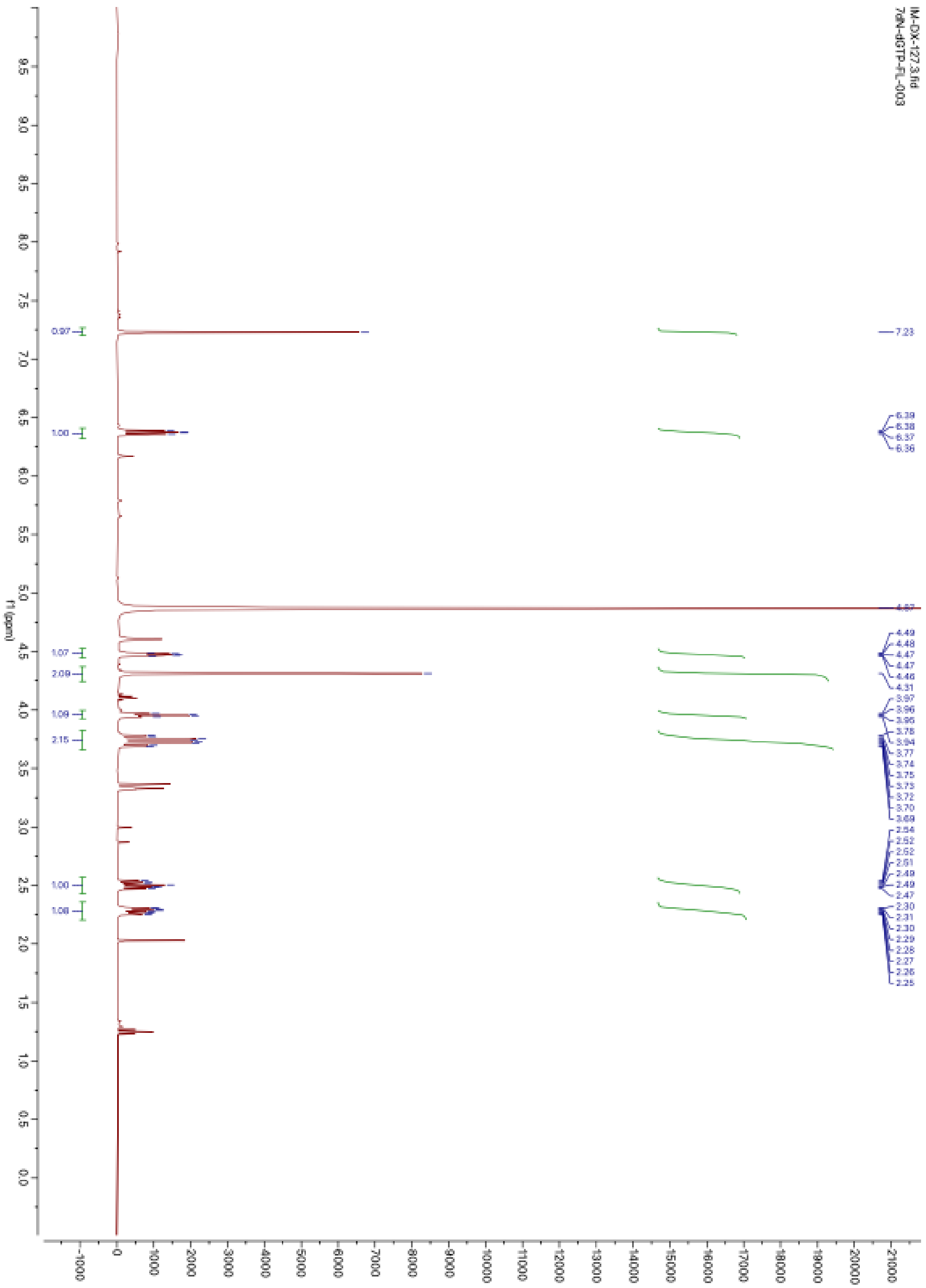

**Supplementary Figure 45.** $^{1}H$ NMR of compound **6** in $CD_3OD$

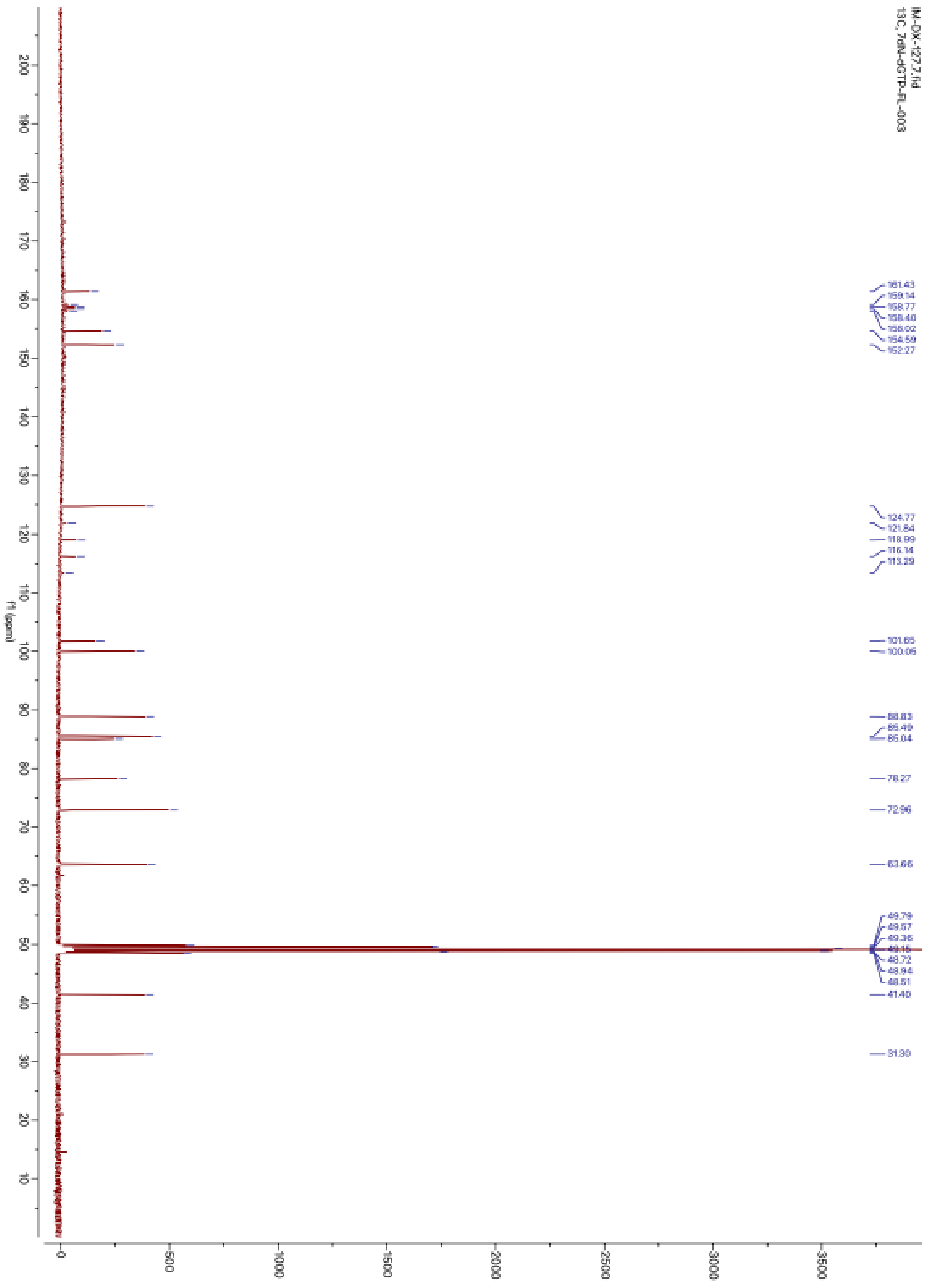

**Supplementary Figure 46.** $^{13}$C NMR of compound **6** in $CD_3OD$

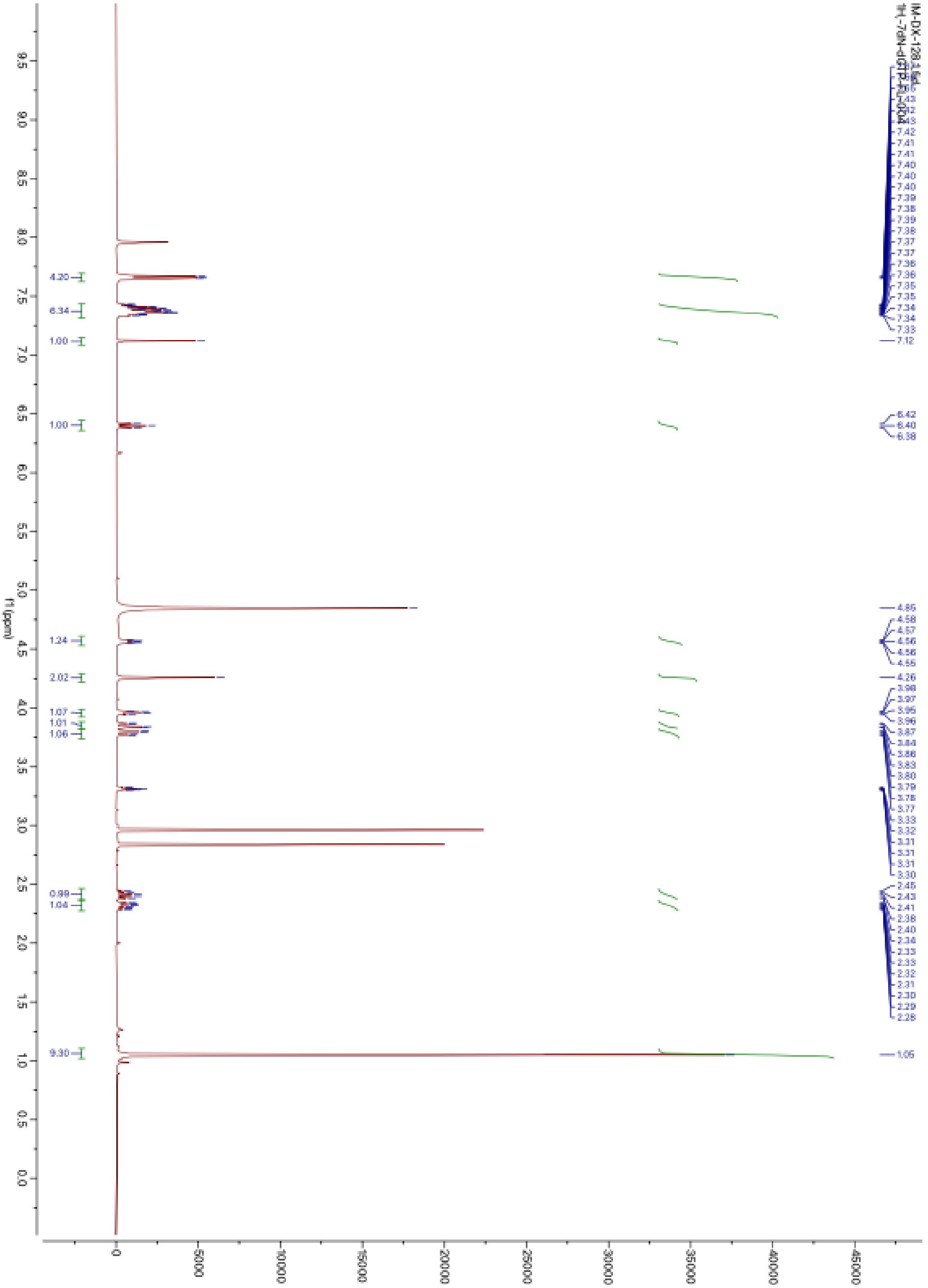

**Supplementary Figure 47.** $^{1}H$ NMR of compound **7** in $CD_3OD$

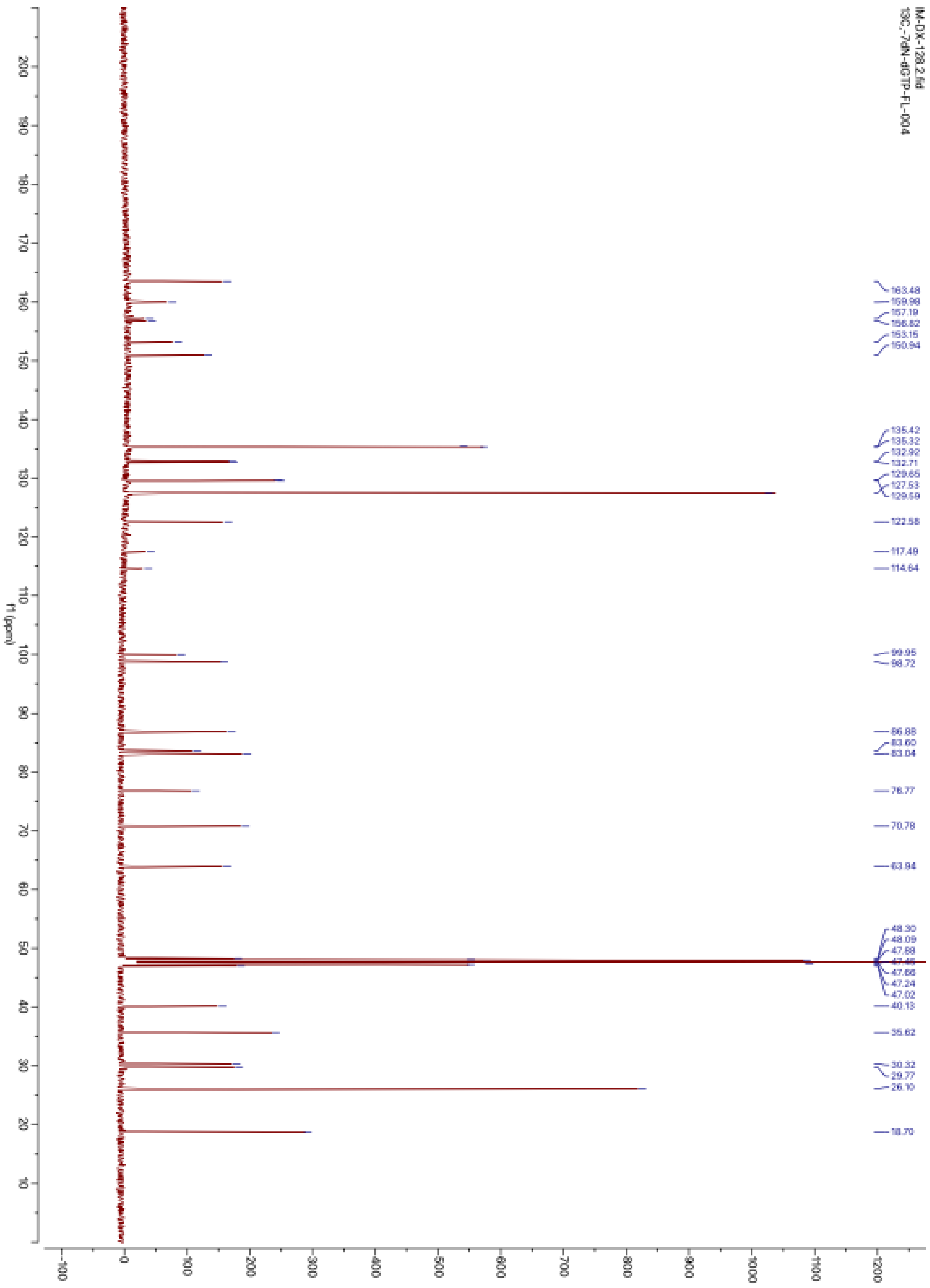

**Supplementary Figure 48.** $^{13}C$ NMR of compound **7** in $CD_3OD$

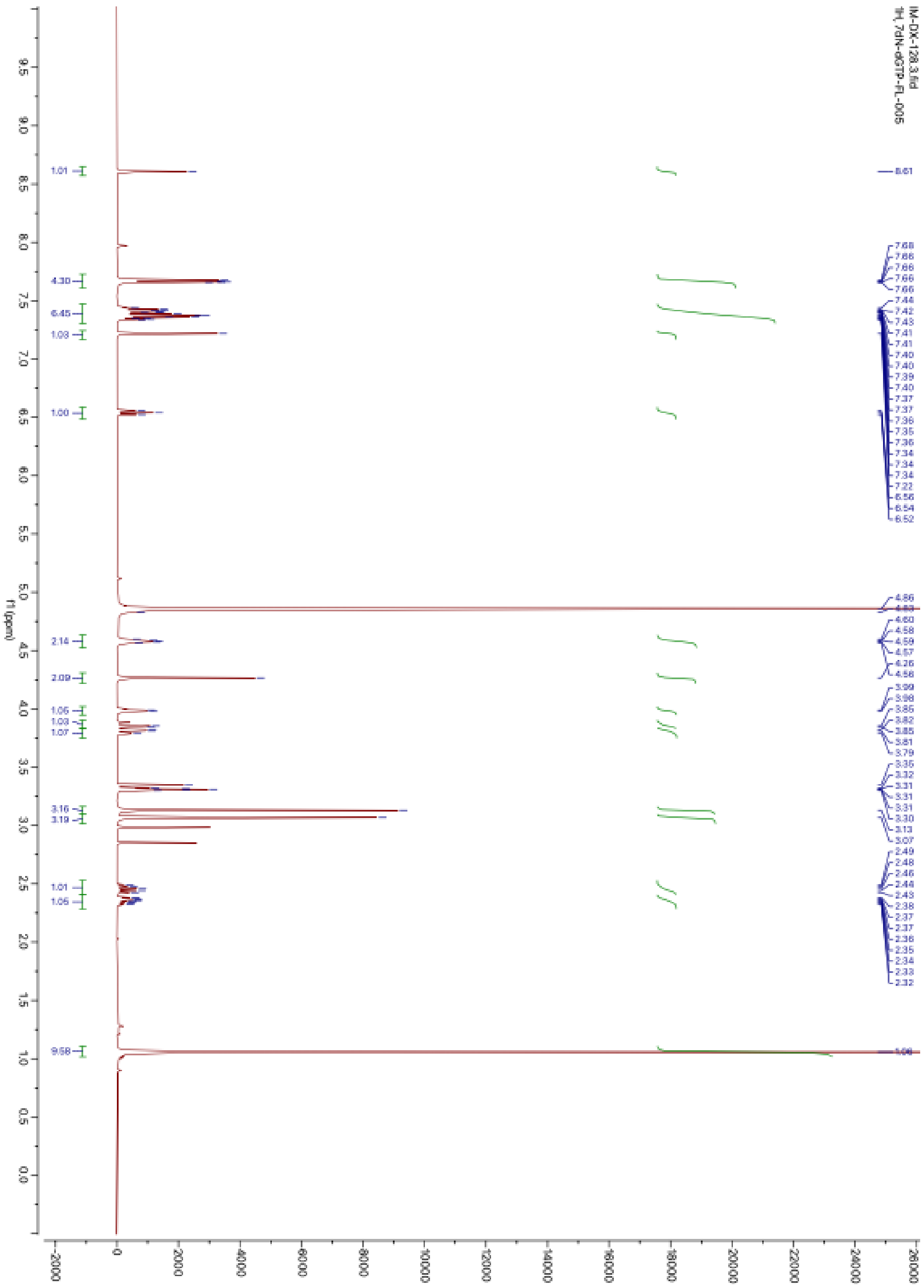

**Supplementary Figure 49.** $^{1}$H NMR of compound **8** in $CD_3OD$

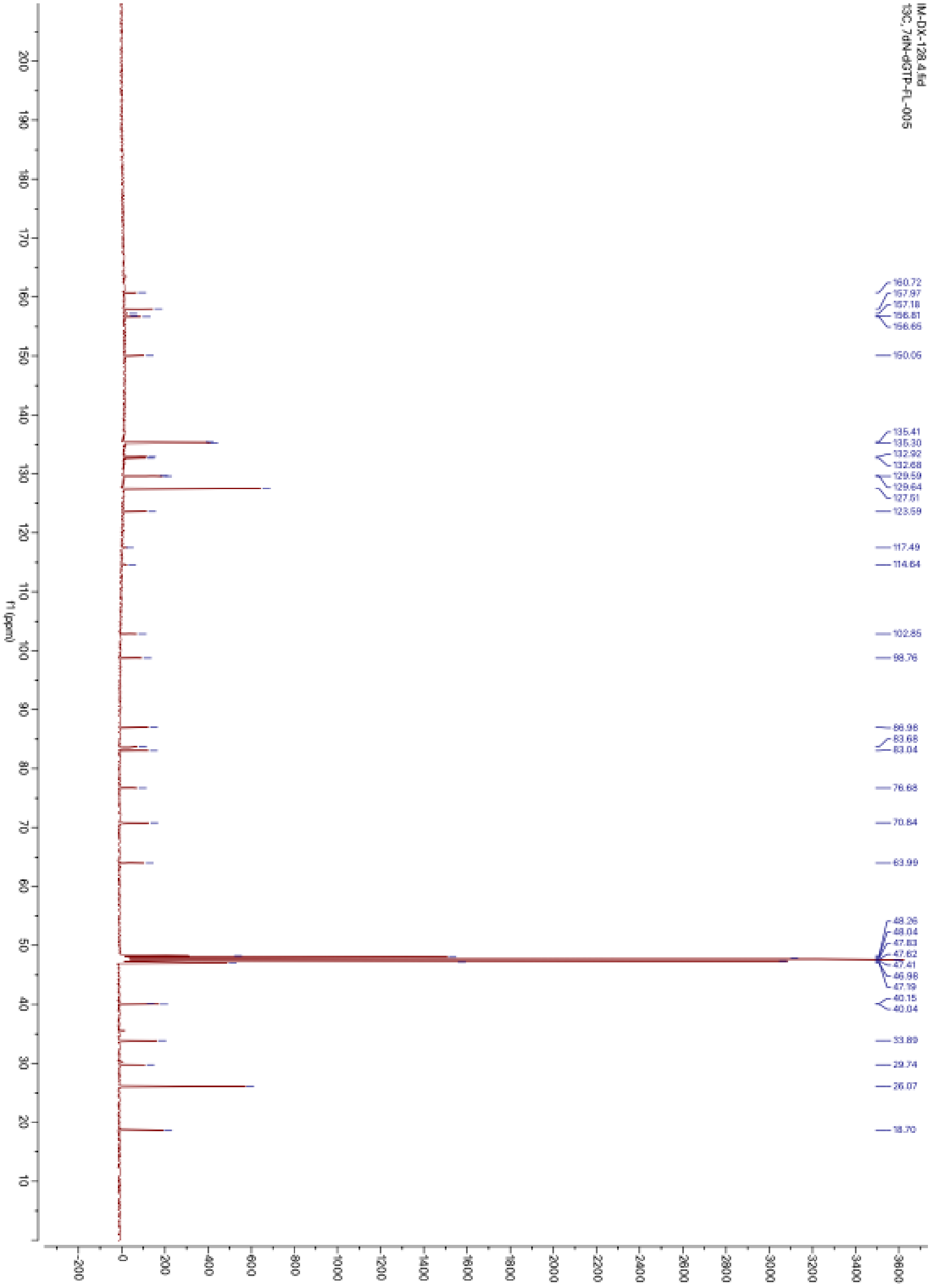

**Supplementary Figure 50.** $^{13}C$ NMR of compound **8** in $CD_3OD$

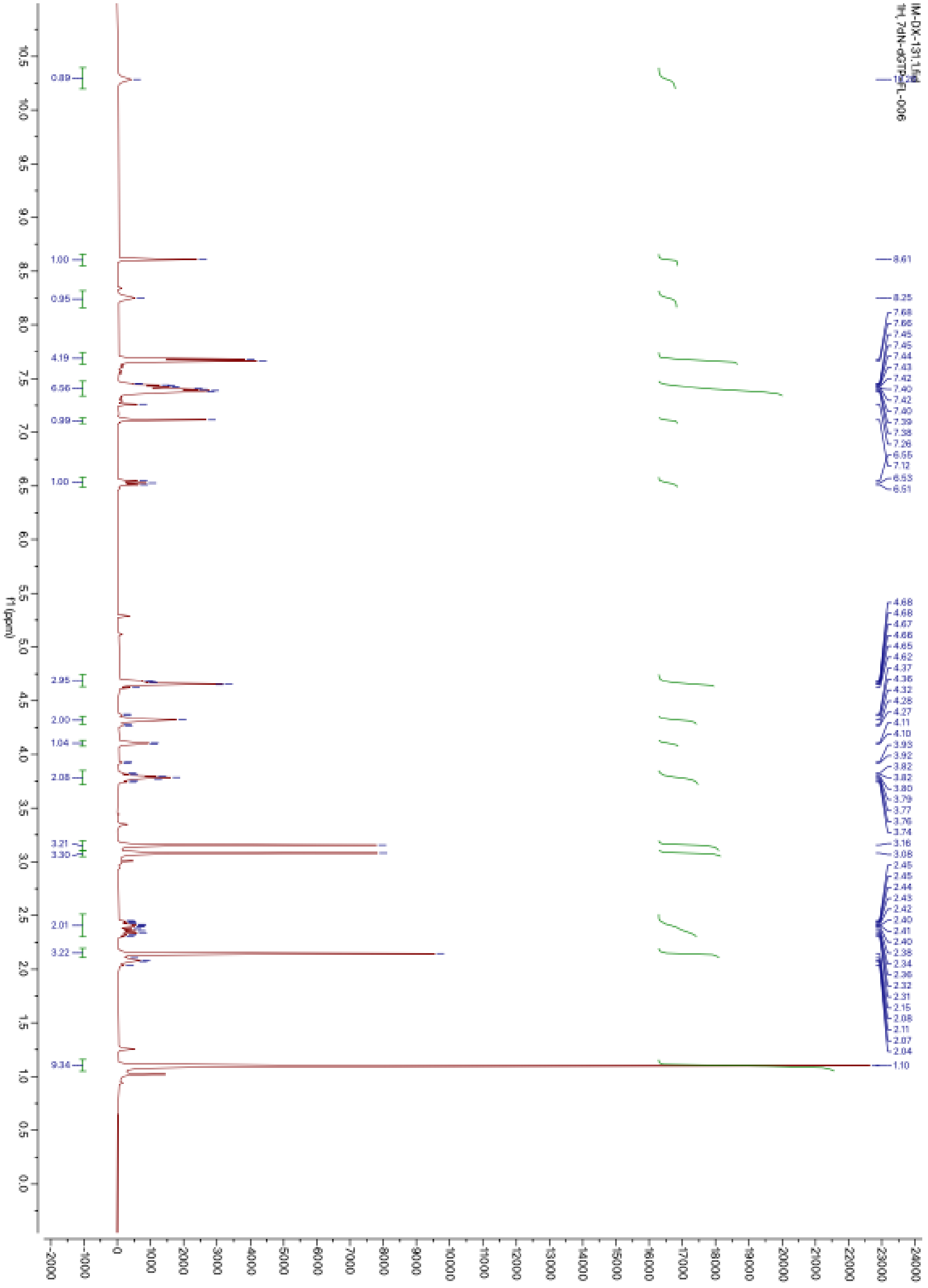

**Supplementary Figure 51.** $^{1}H$ NMR of compound **9** in $CDCl_3$

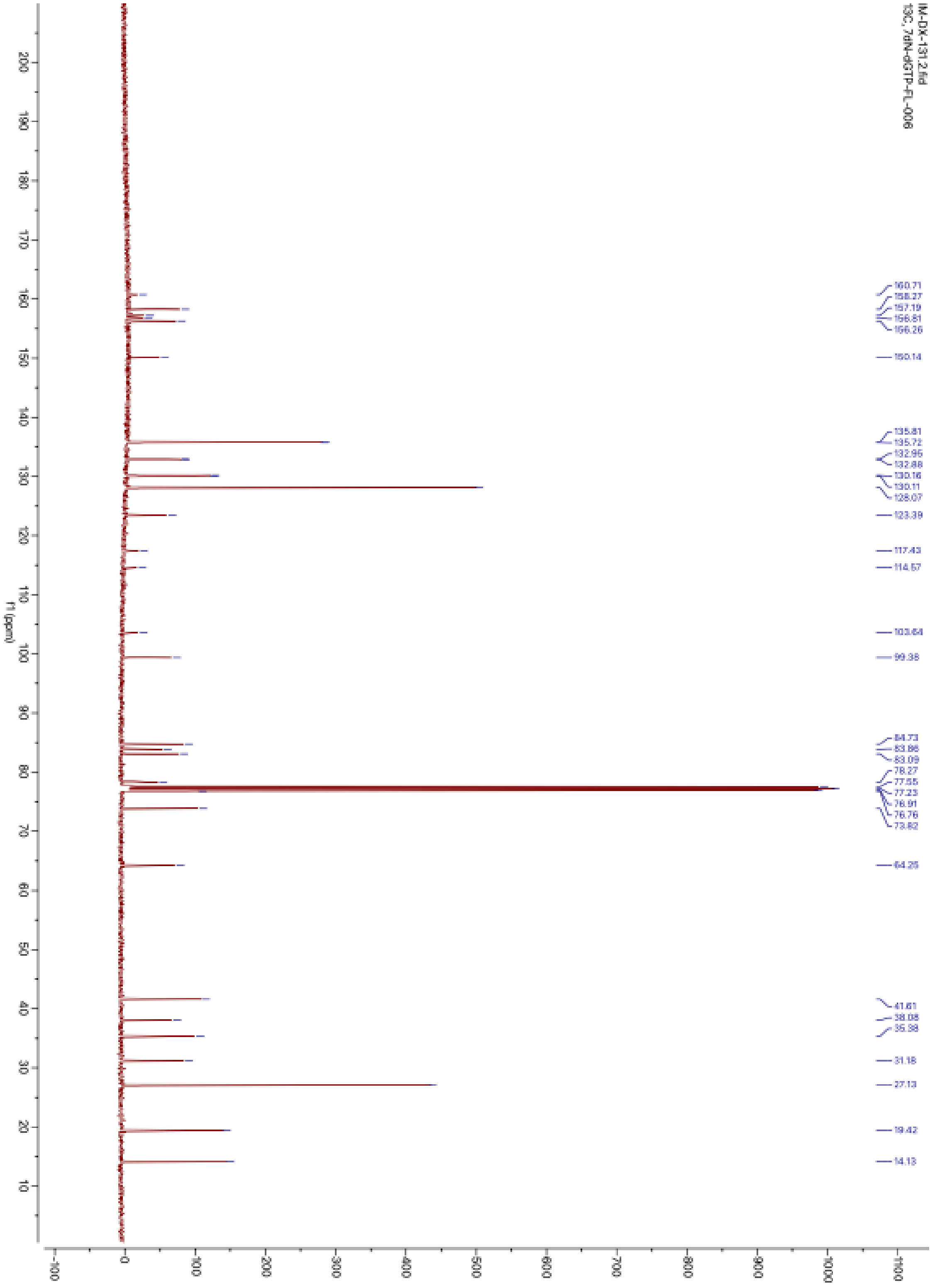

**Supplementary Figure 52.** $^{13}C$ NMR of compound **9** in $CDCl_3$

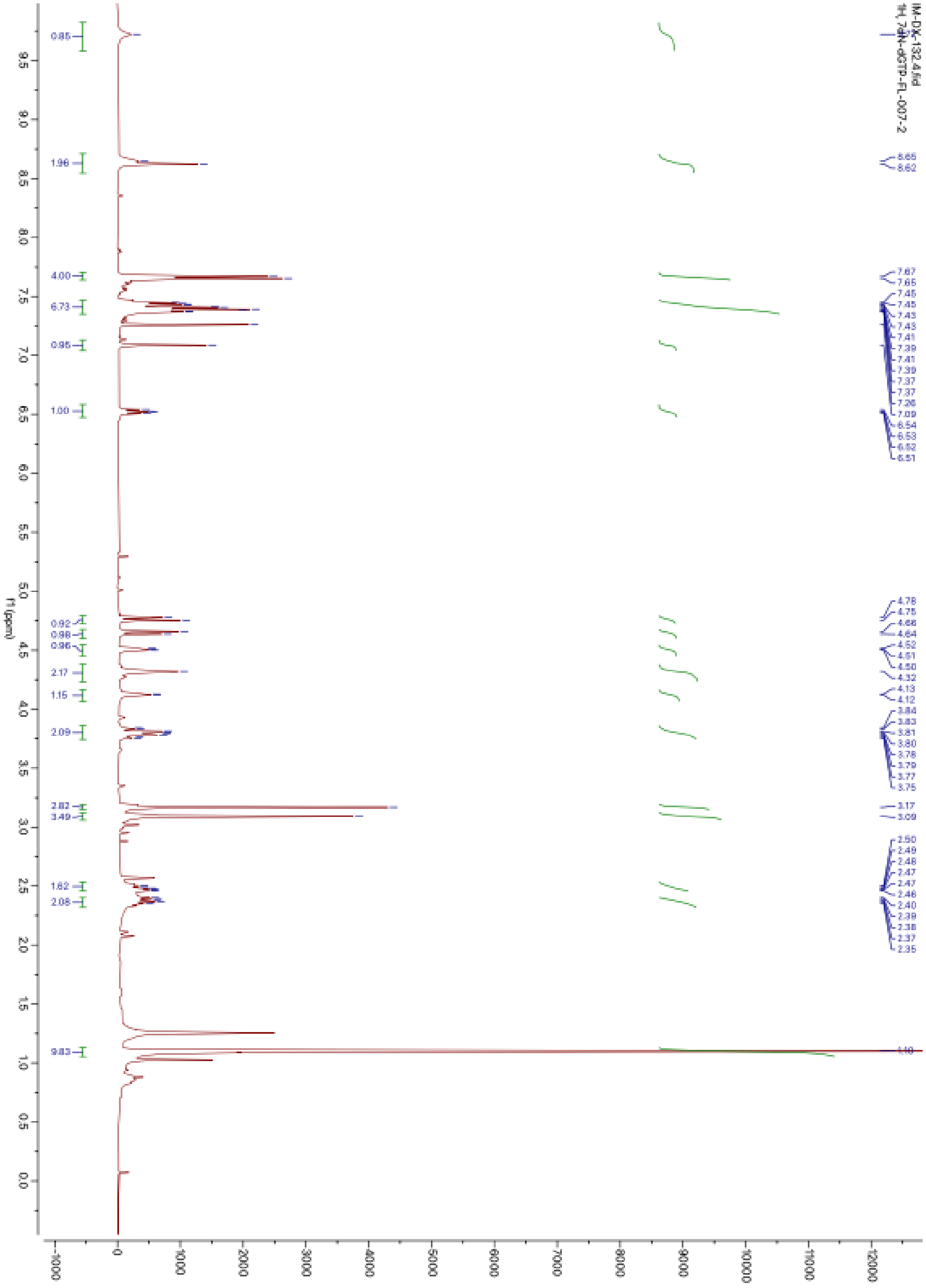

**Supplementary Figure 53.** $^{1}$H NMR of compound **10** in $CDCl_3$

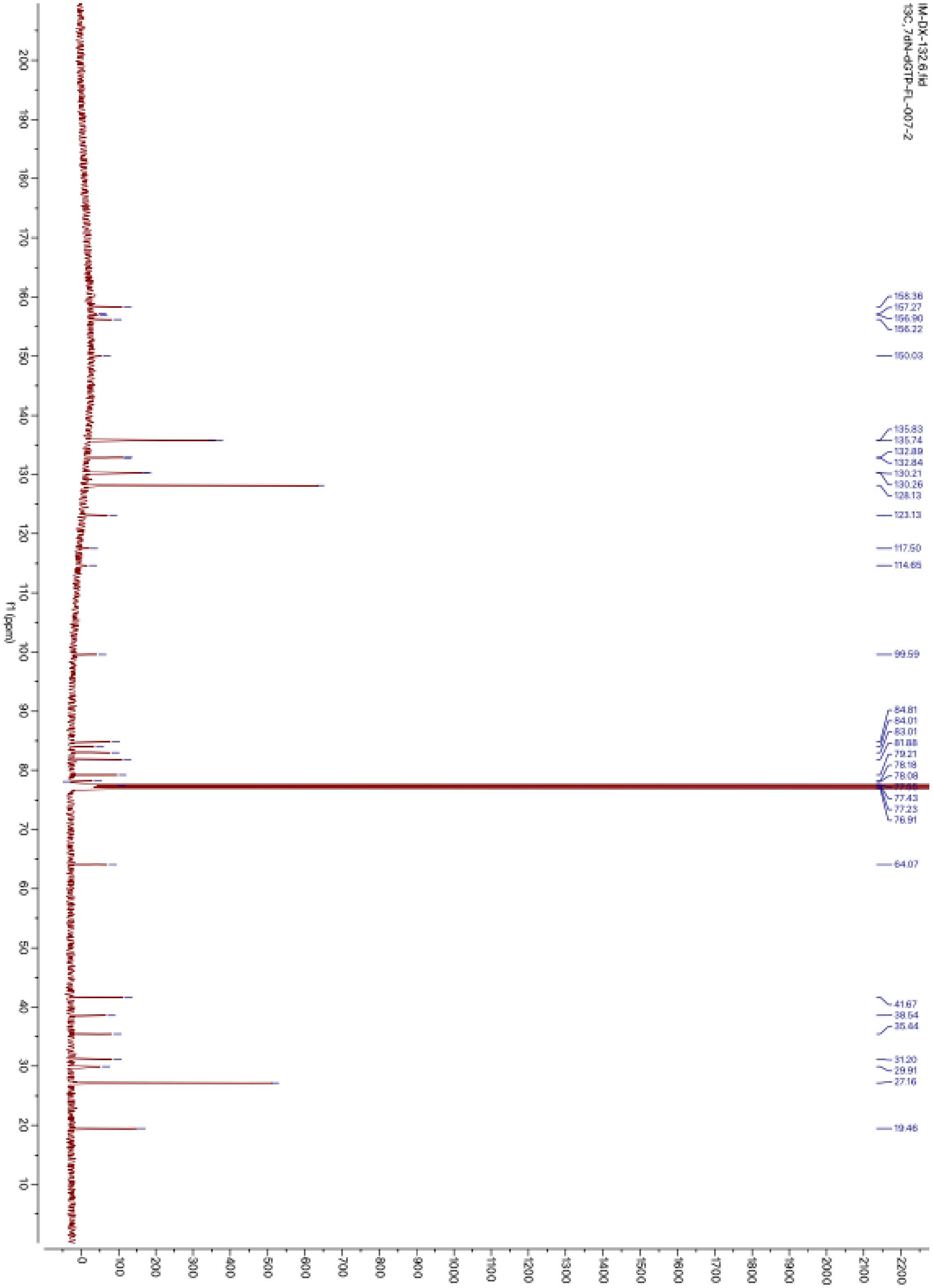

**Supplementary Figure 54.** $^{13}C$ NMR of compound **10** in $CDCl_3$

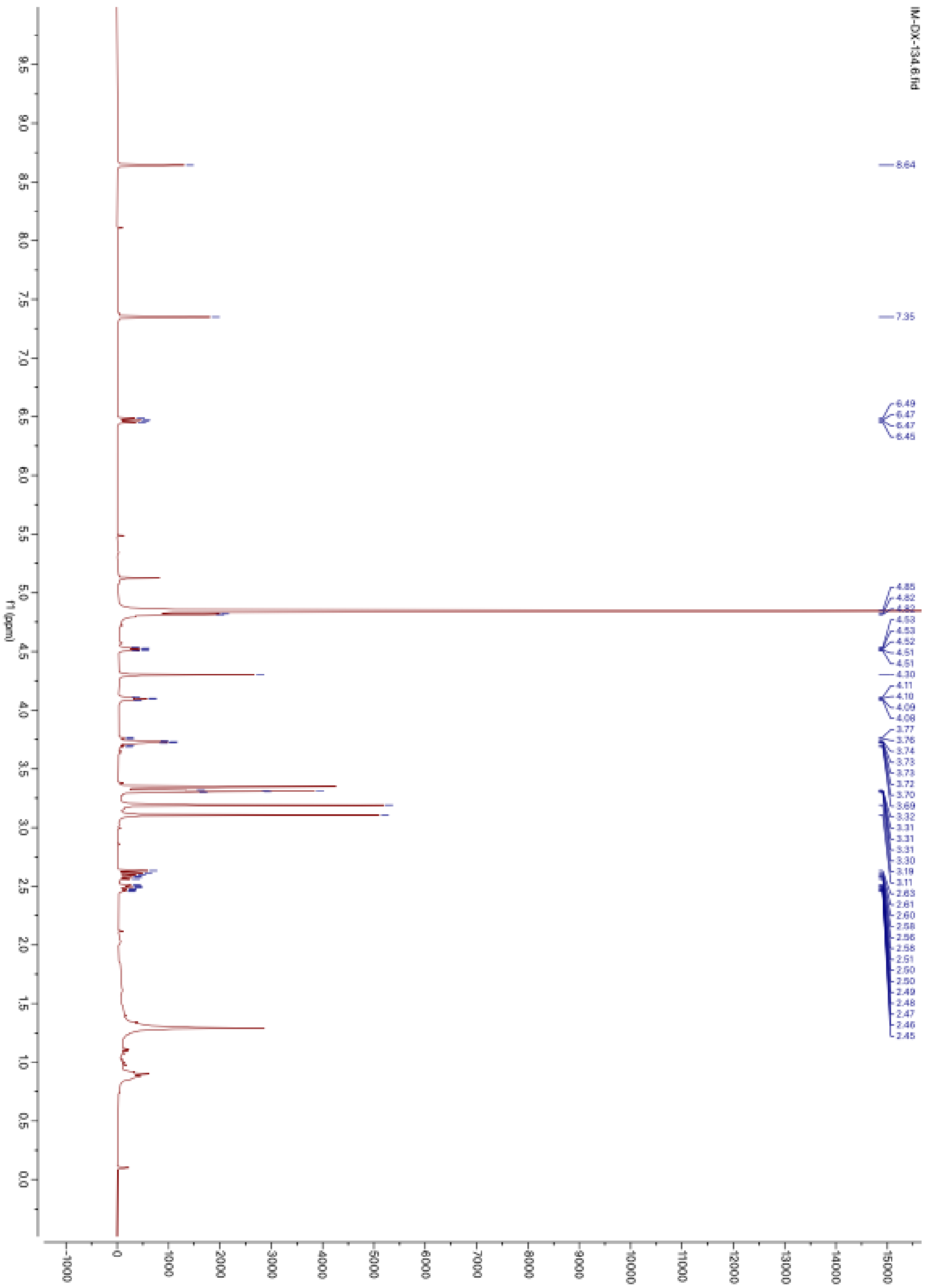

**Supplementary Figure 55.** $^{1}H$ NMR of compound **11** in $CD_3OD$

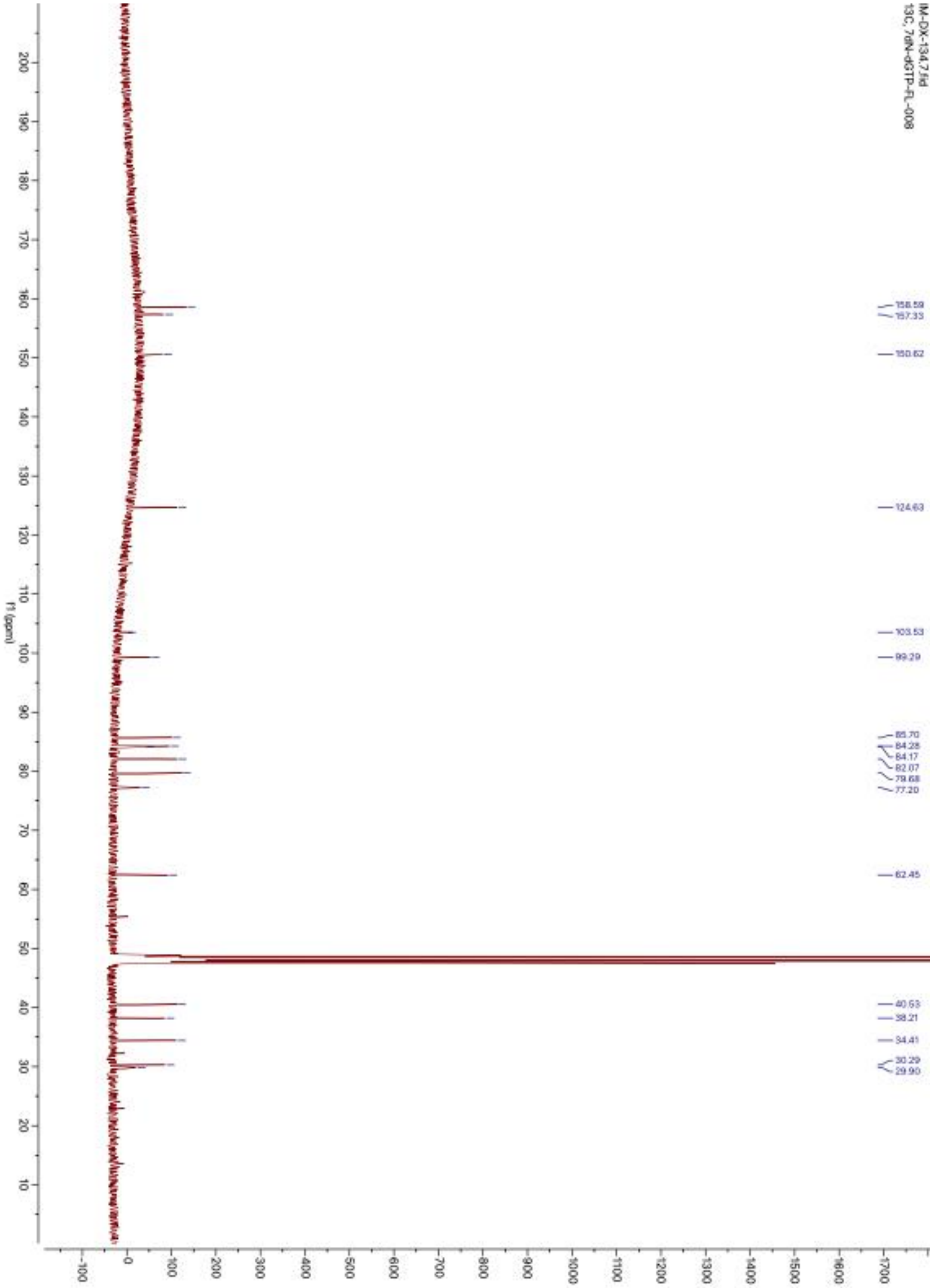

**Supplementary Figure 56.** $^{13}C$ NMR of compound **11** in $CD_3OD$

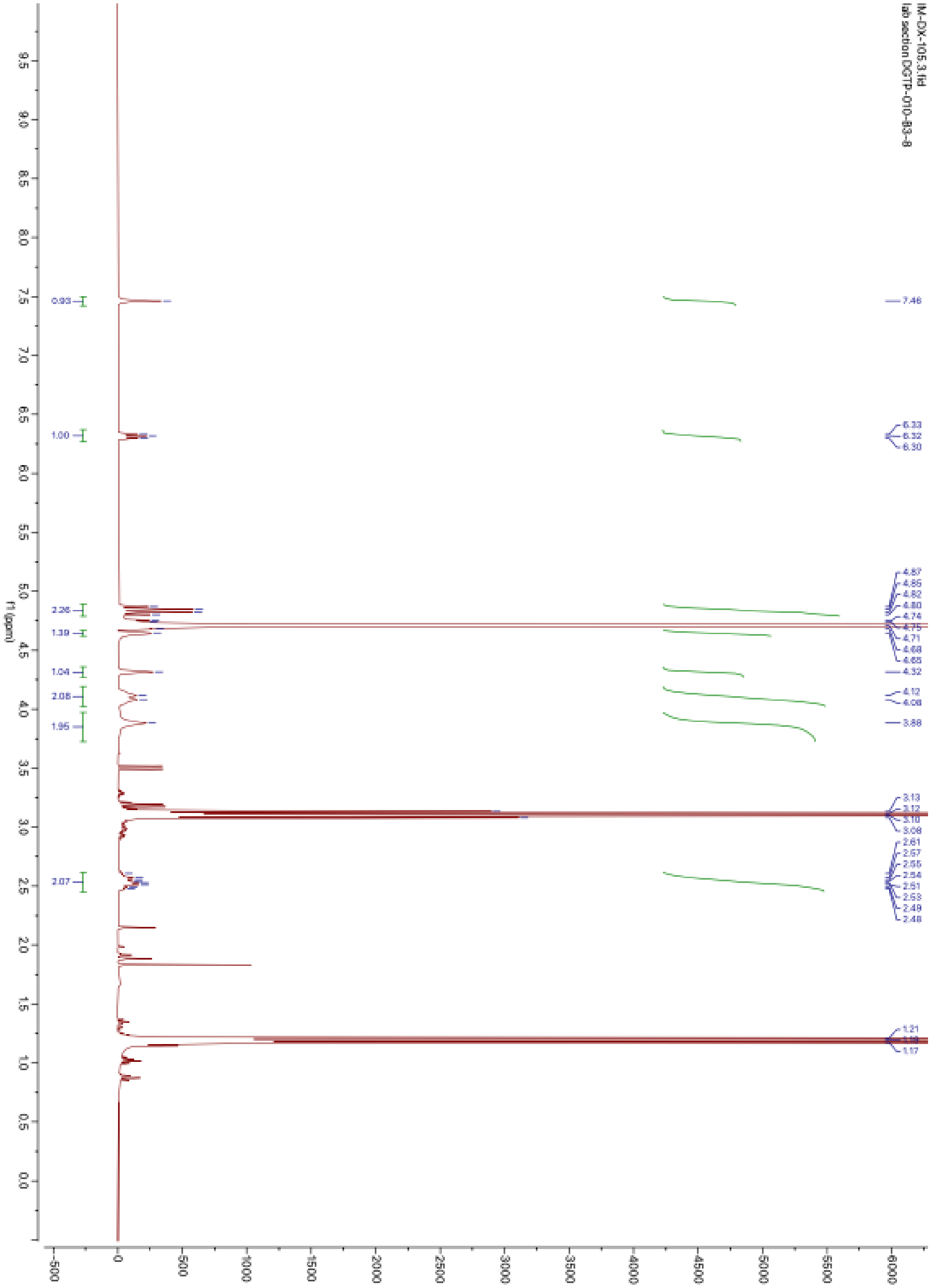

**Supplementary Figure 57.** $^{1}$H NMR of compound **12** in $D_2O$

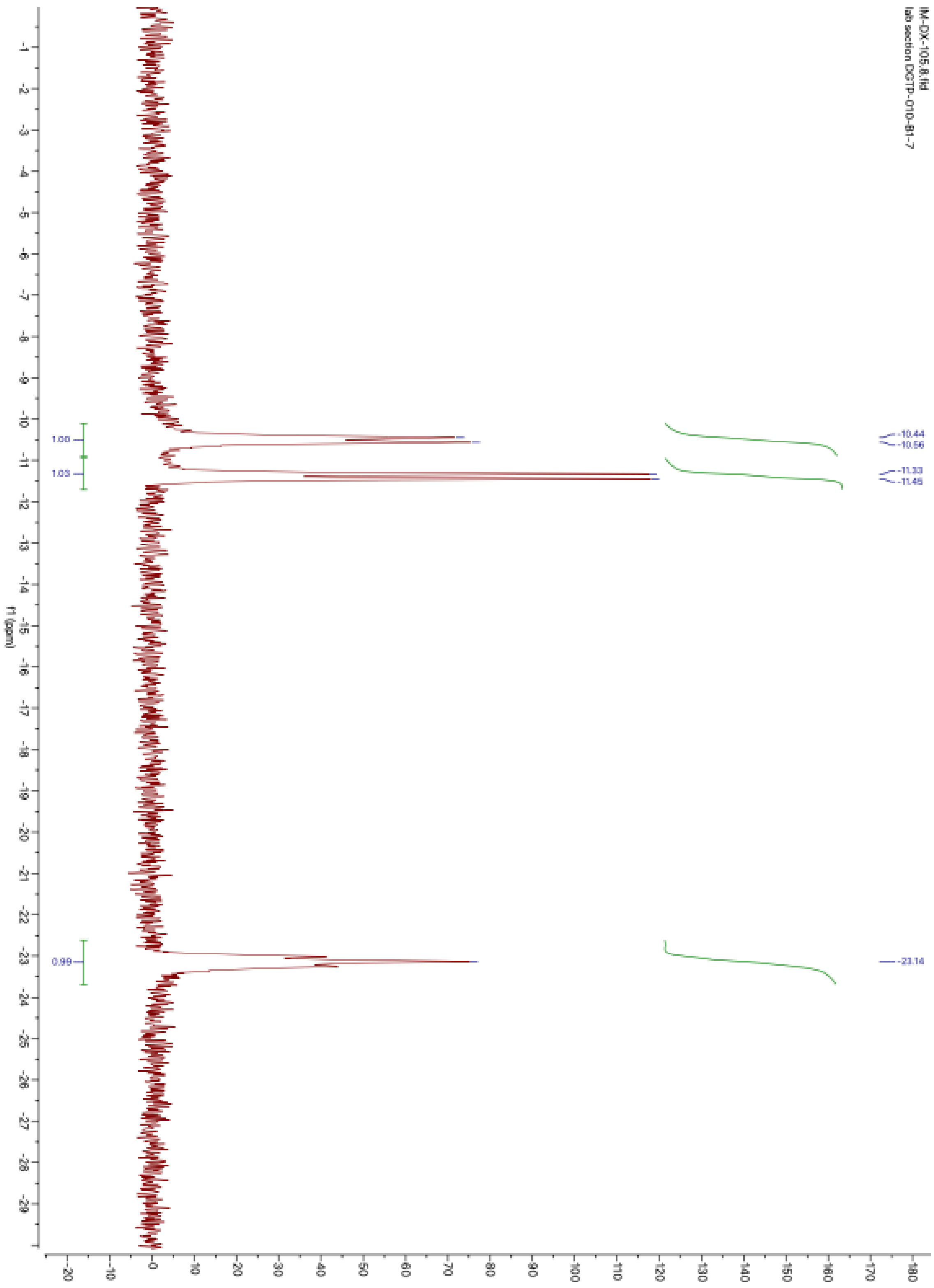

**Supplementary Figure 58.** $^{31}P$ NMR of compound **12** in $D_2O$

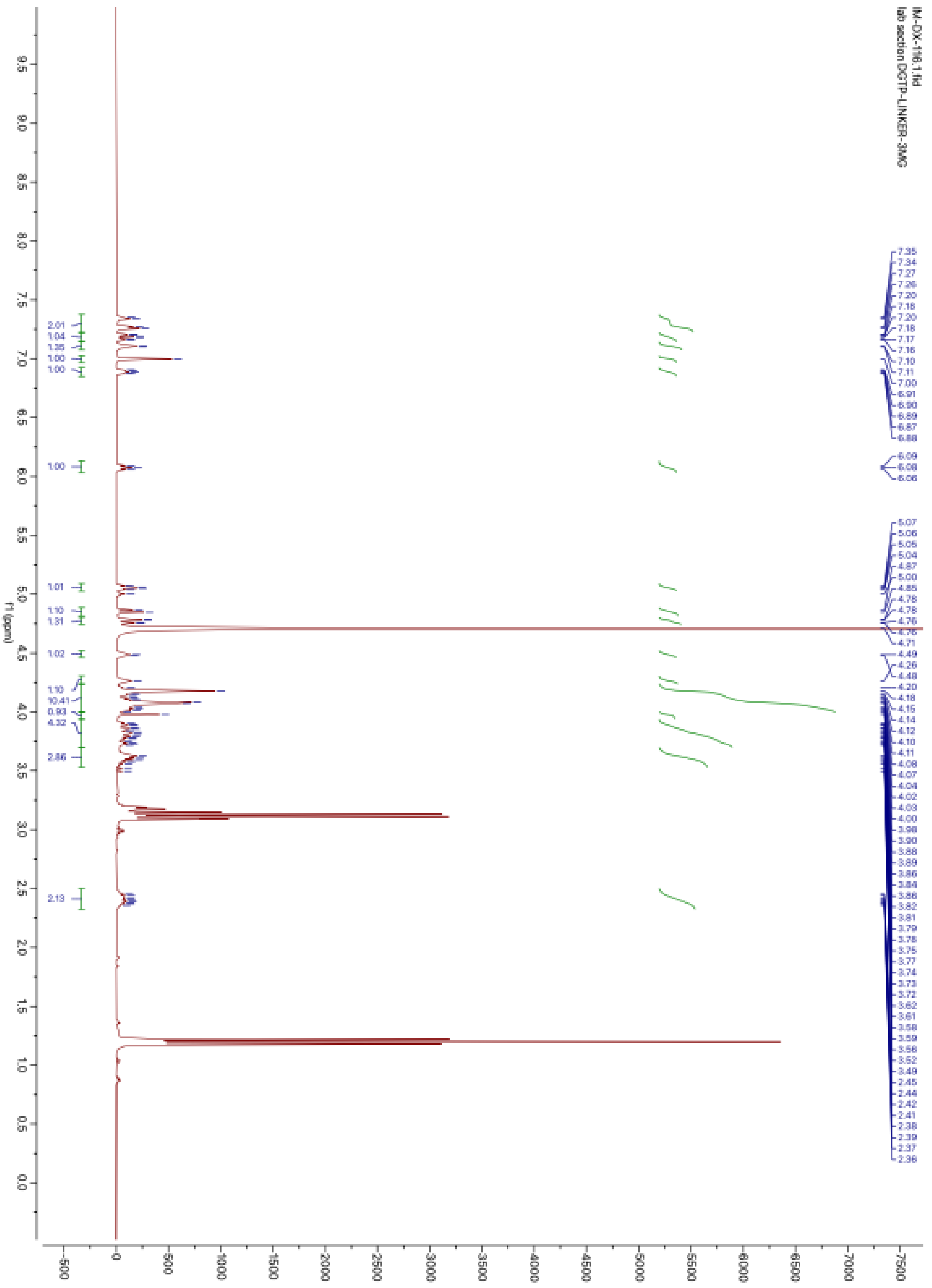

**Supplementary Figure 59.** $^{1}H$ NMR of compound **13** in $D_2O$

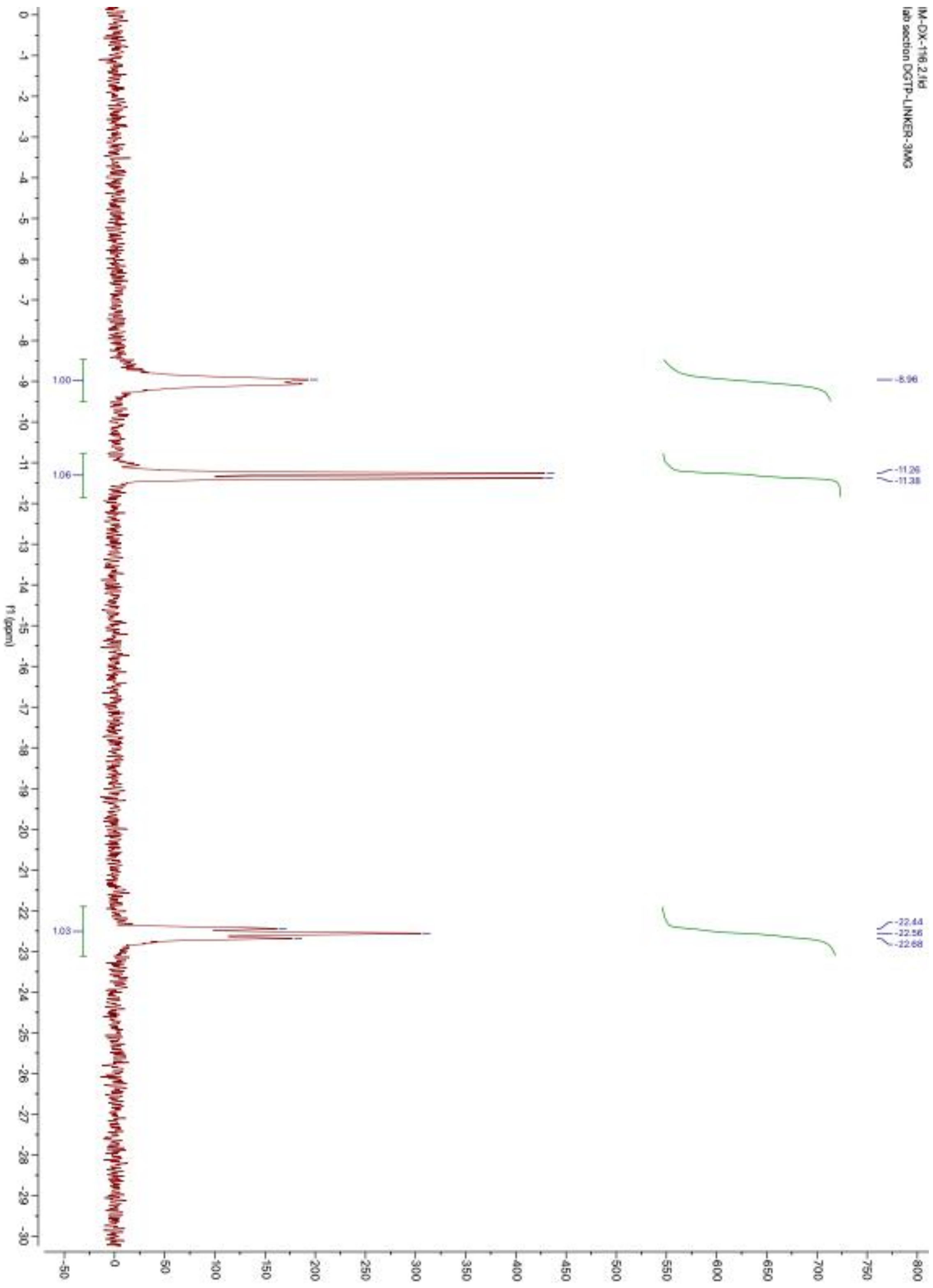

**Supplementary Figure 60.** $^{31}P$ NMR of compound **13** in $D_2O$

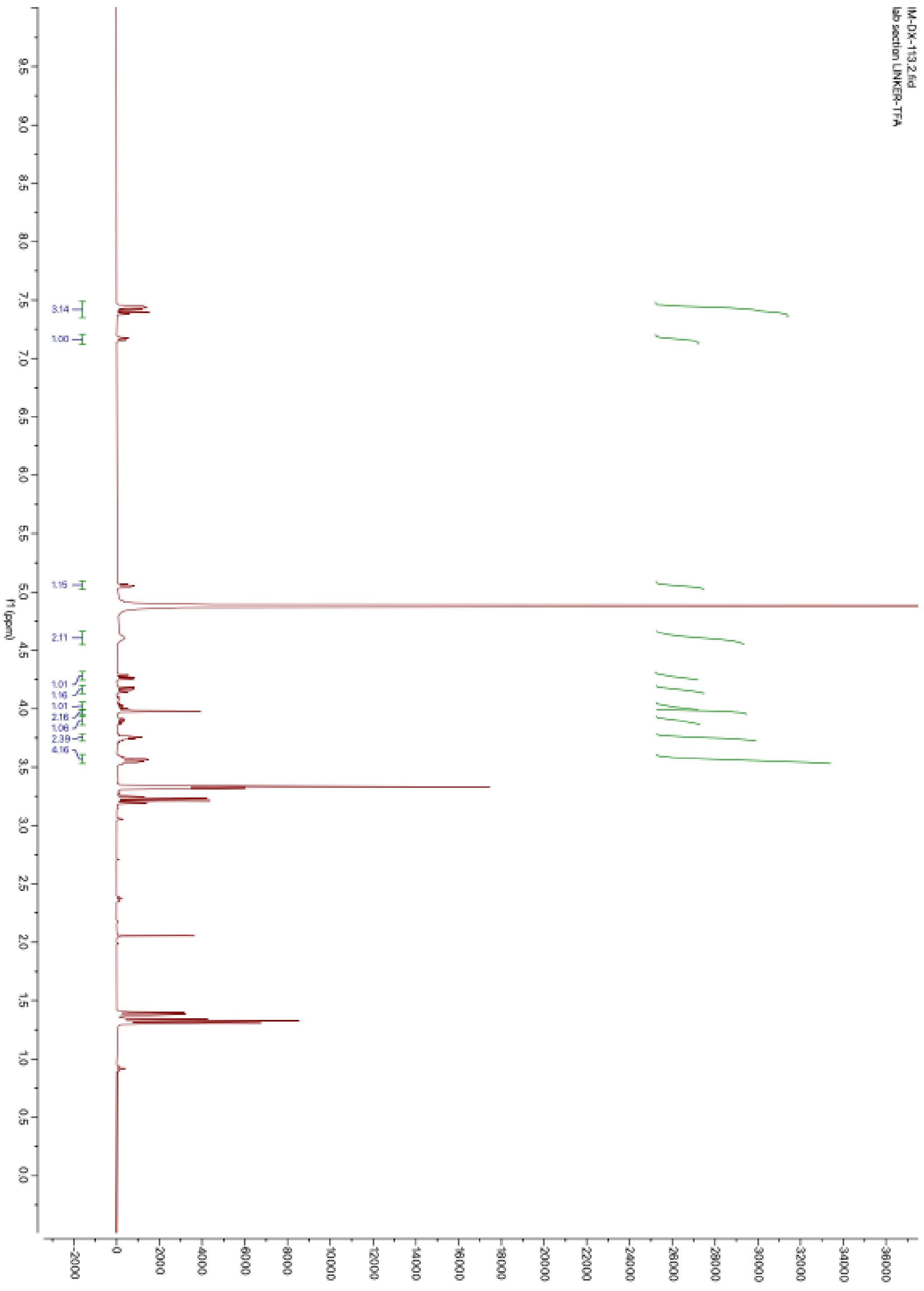

**Supplementary Figure 61.** $^{1}$H NMR of compound **S-2** in $CD_3OD$

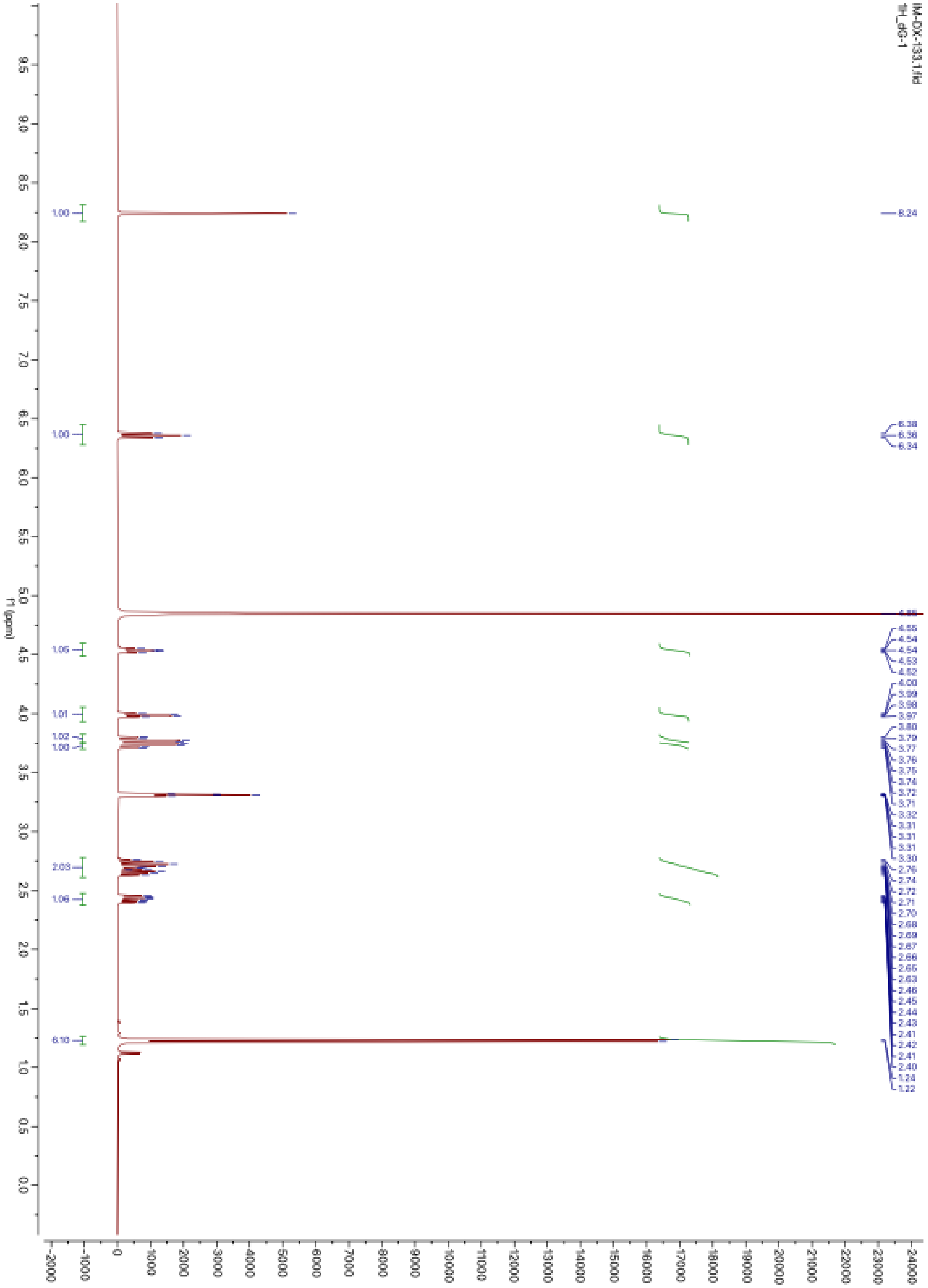

**Supplementary Figure 62.** $^{1}$H NMR of compound **S-4** in $CD_3OD$

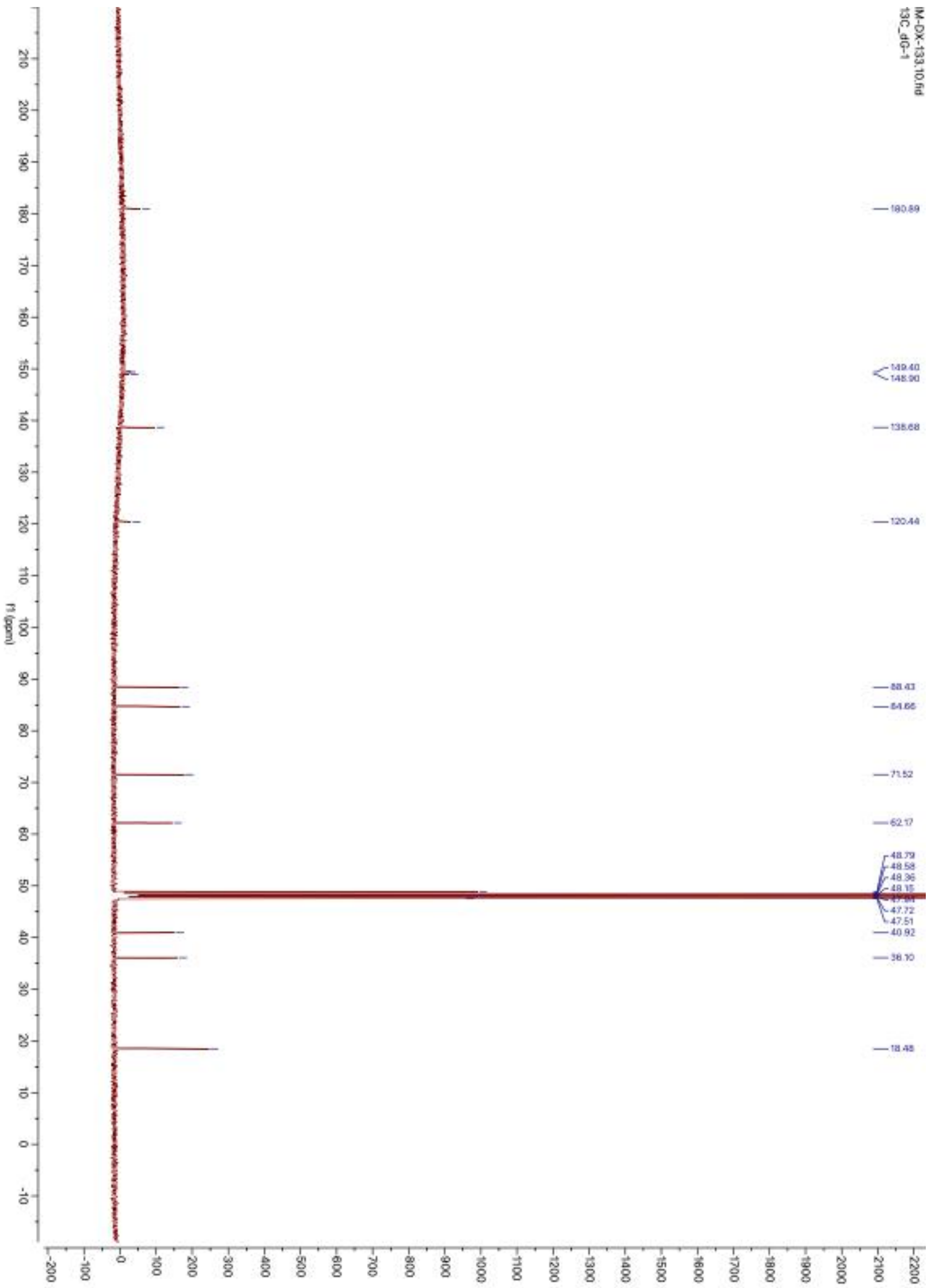

**Supplementary Figure 63.** $^{13}$C NMR of compound **S-4** in $CD_3OD$

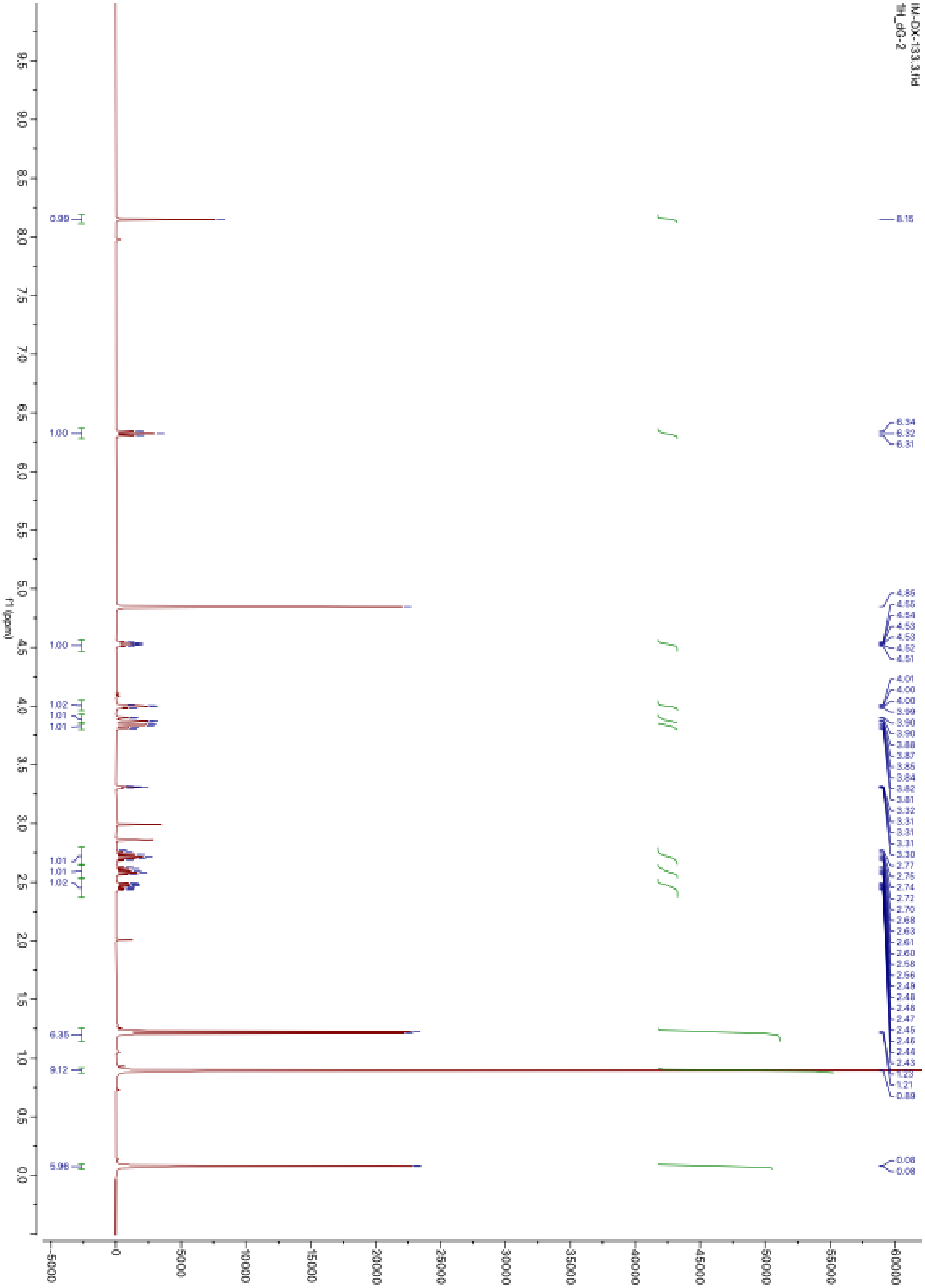

**Supplementary Figure 64.** $^{1}$H NMR of compound **S-5** in $CD_3OD$

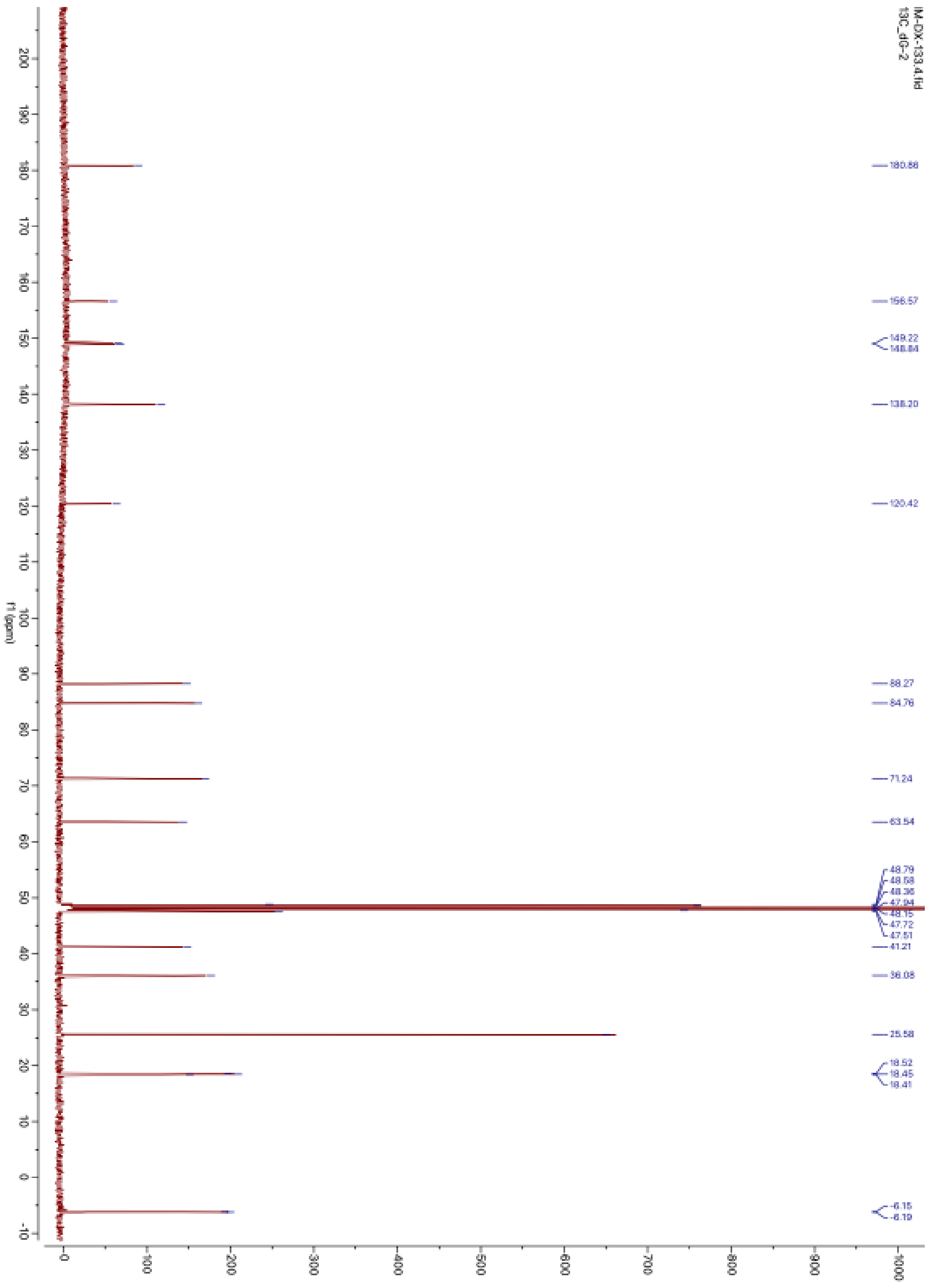

**Supplementary Figure 65.** $^{13}C$ NMR of compound **S-5** in $CD_3OD$

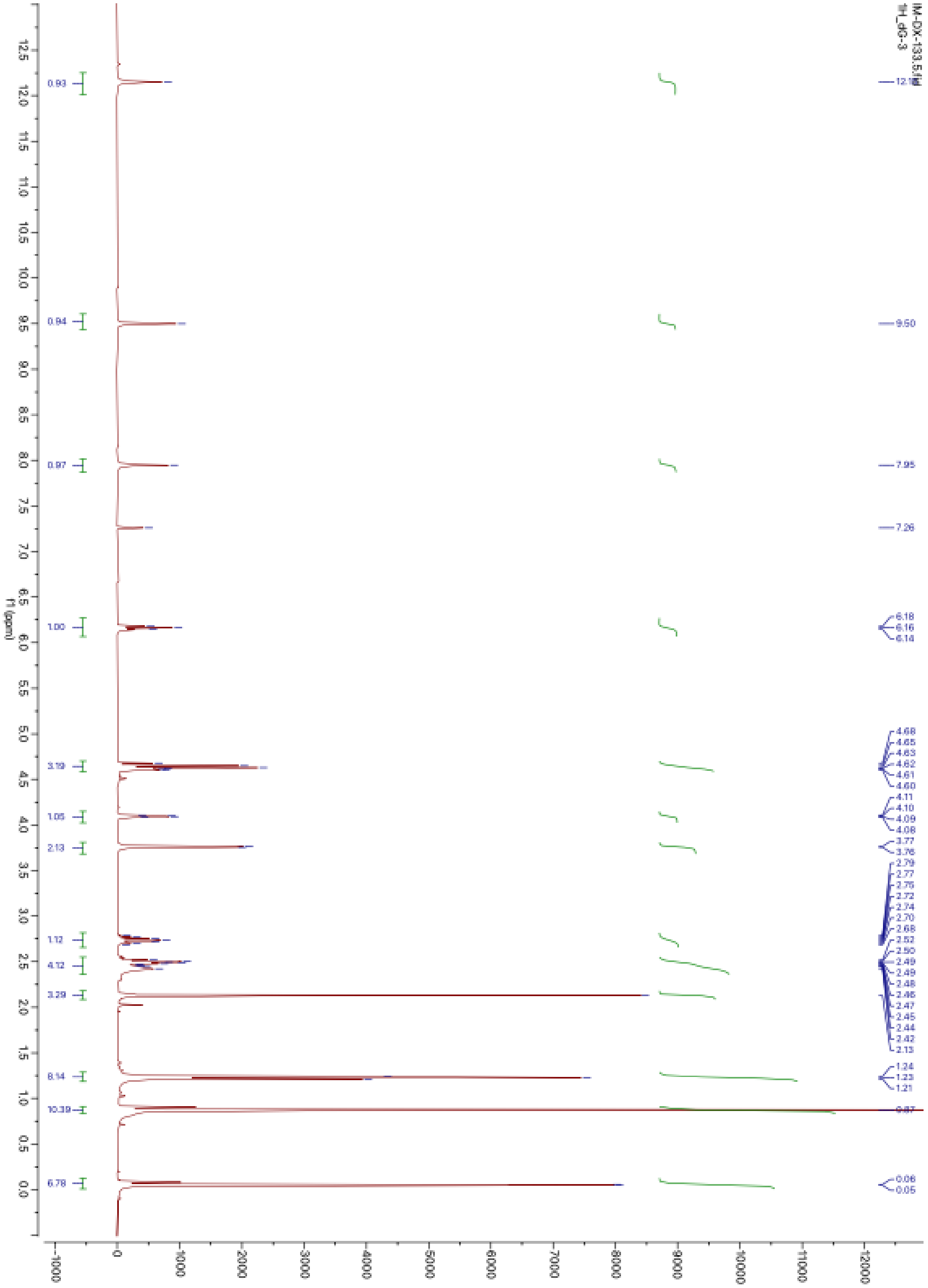

**Supplementary Figure 66.** $^{1}$H NMR of compound **S-6** in $CDCl_3$

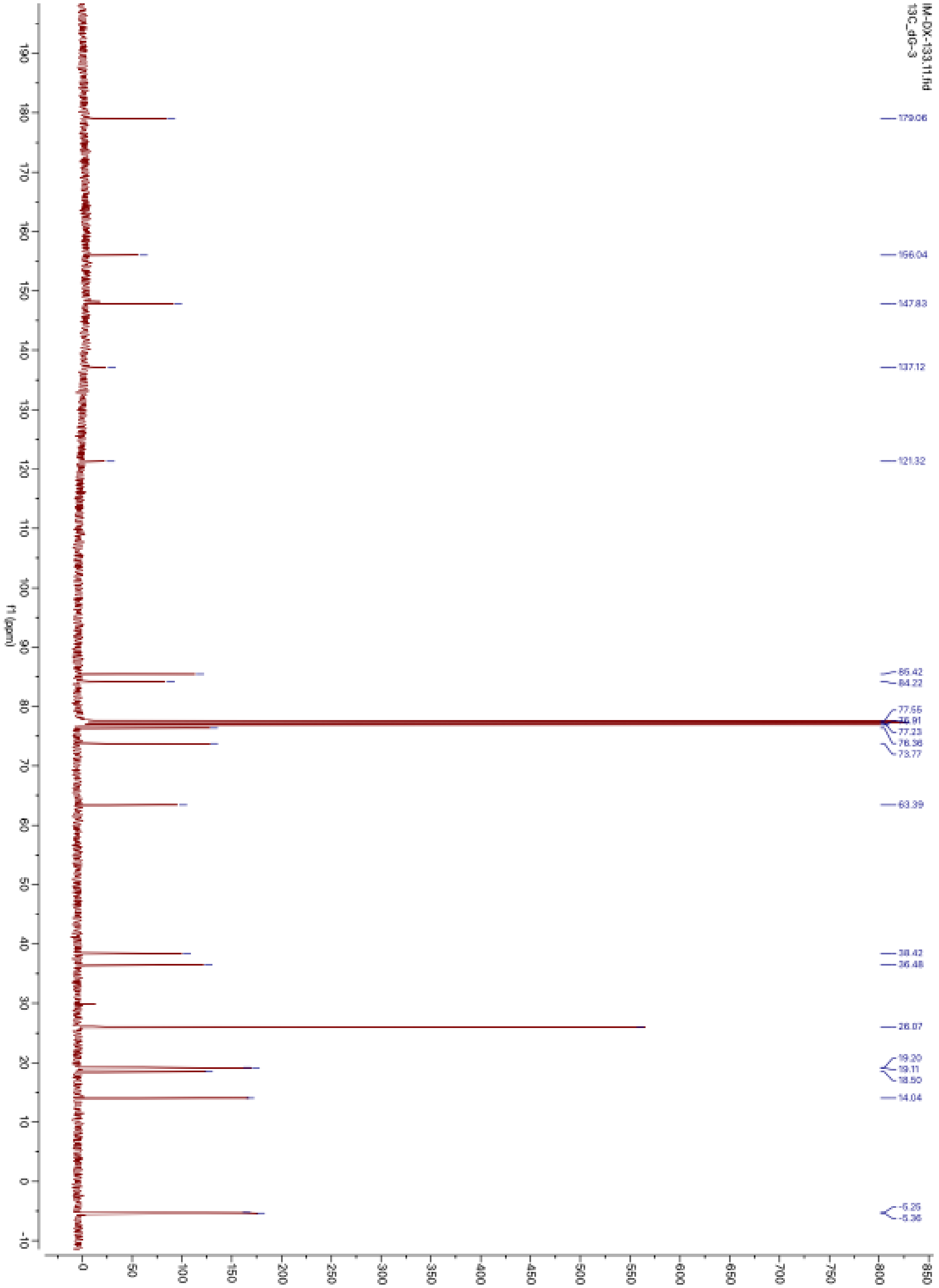

**Supplementary Figure 67.** $^{13}C$ NMR of compound **S-6** in $CDCl_3$

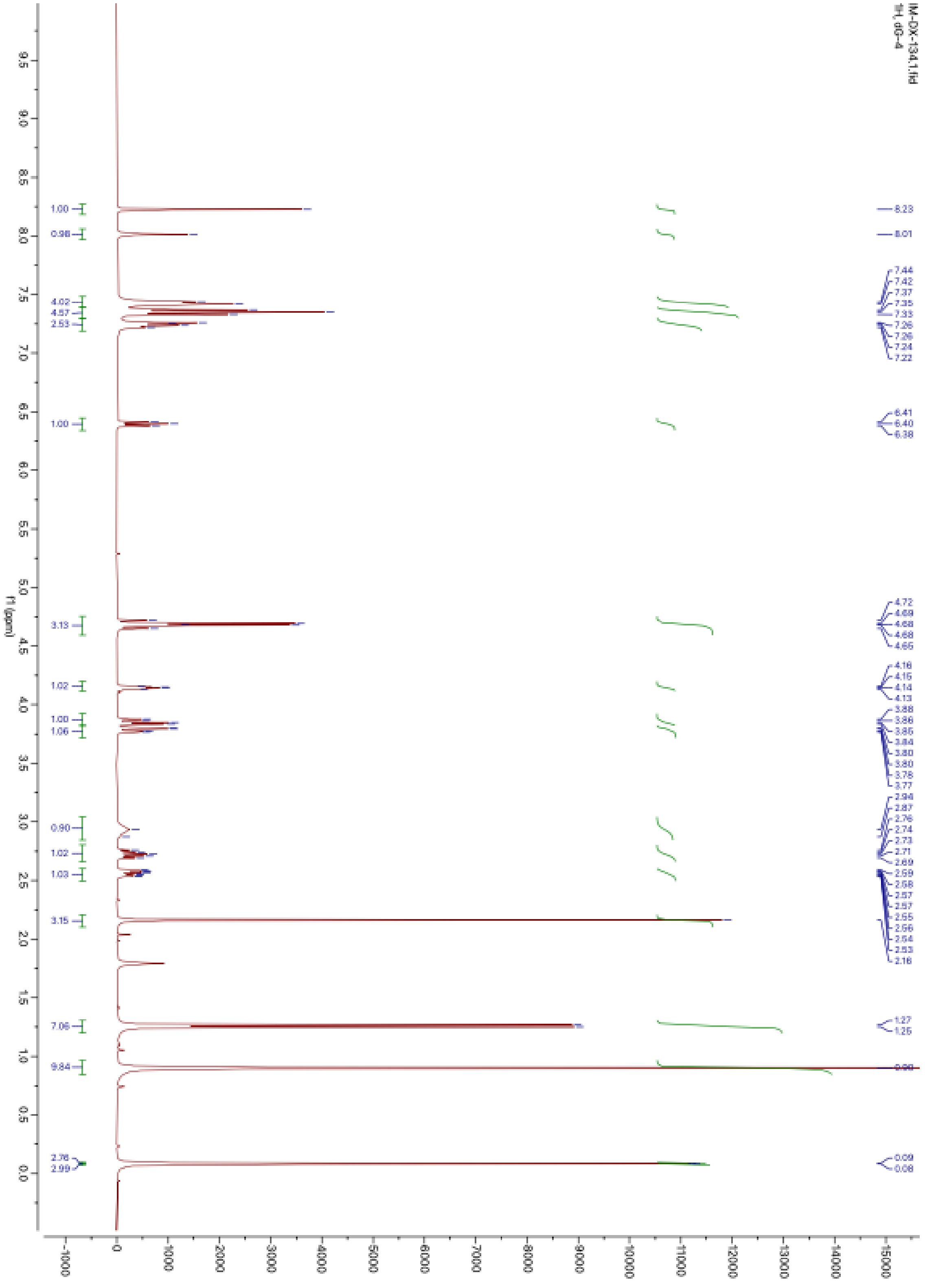

**Supplementary Figure 68.** $^{1}$H NMR of compound **S-7** in $CDCl_3$

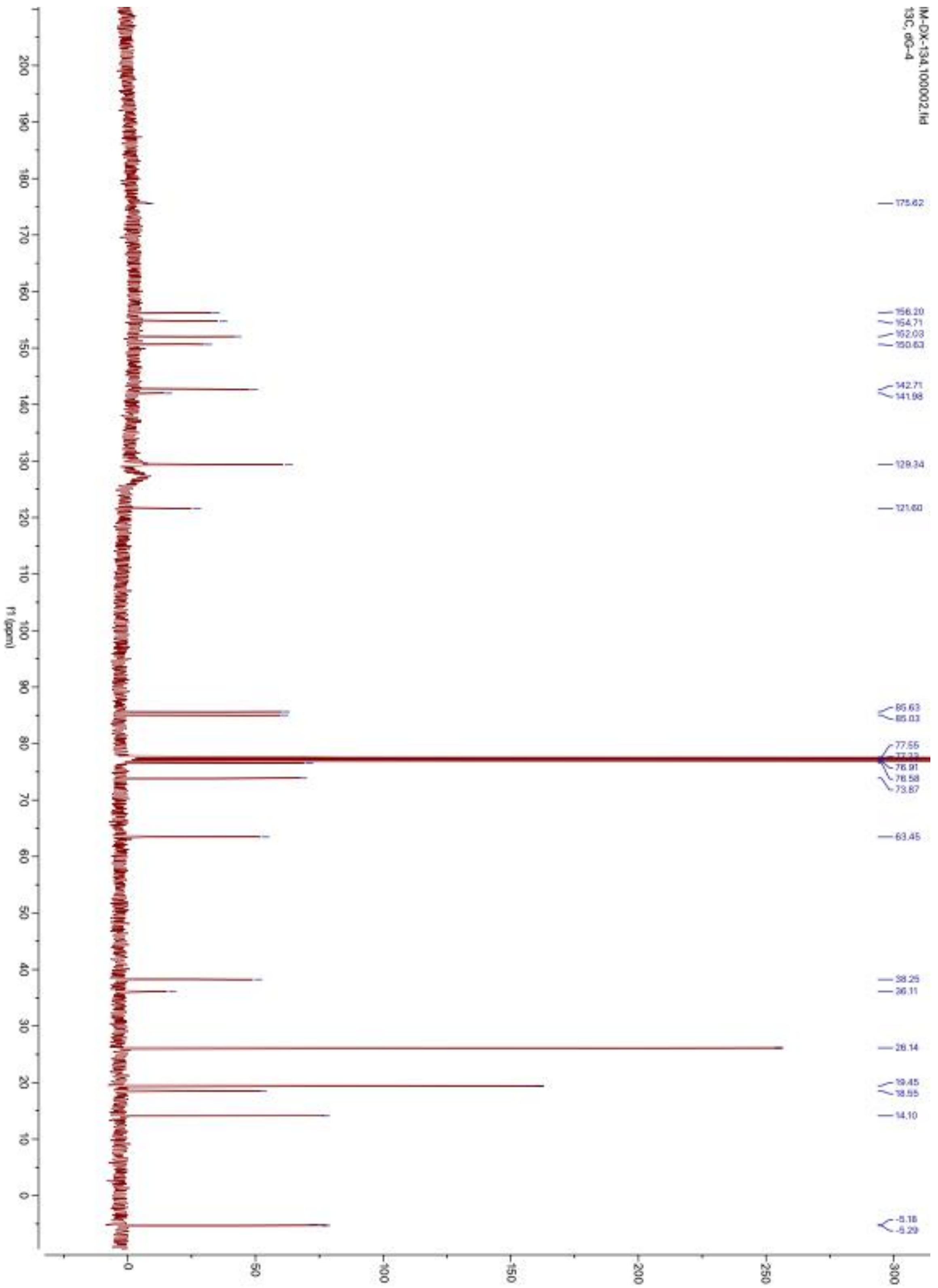

**Supplementary Figure 69.** $^{13}C$ NMR of compound **S-7** in $CDCl_3$

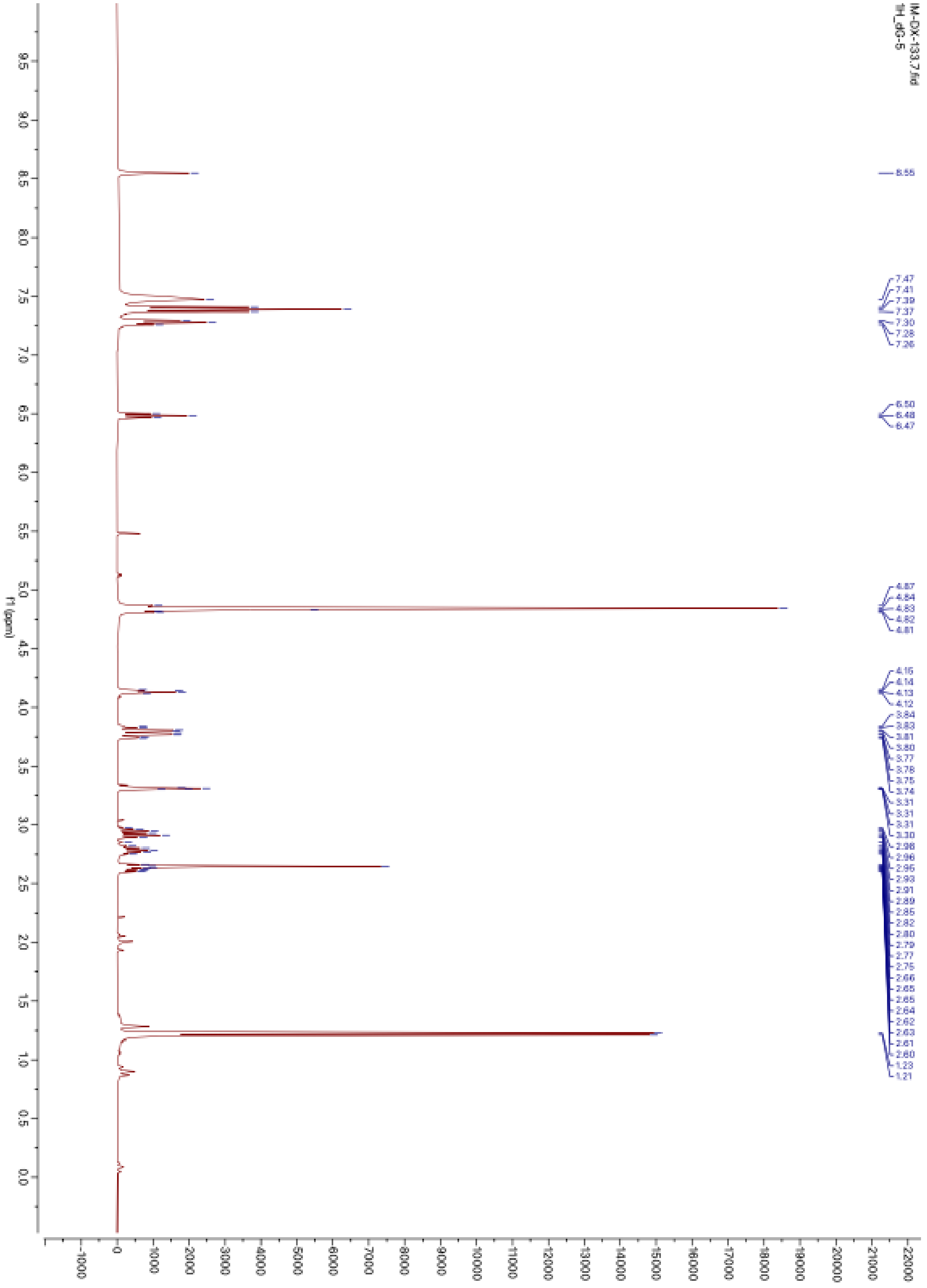

**Supplementary Figure 70.** $^{1}$H NMR of compound **S-8** in $CD_3OD$

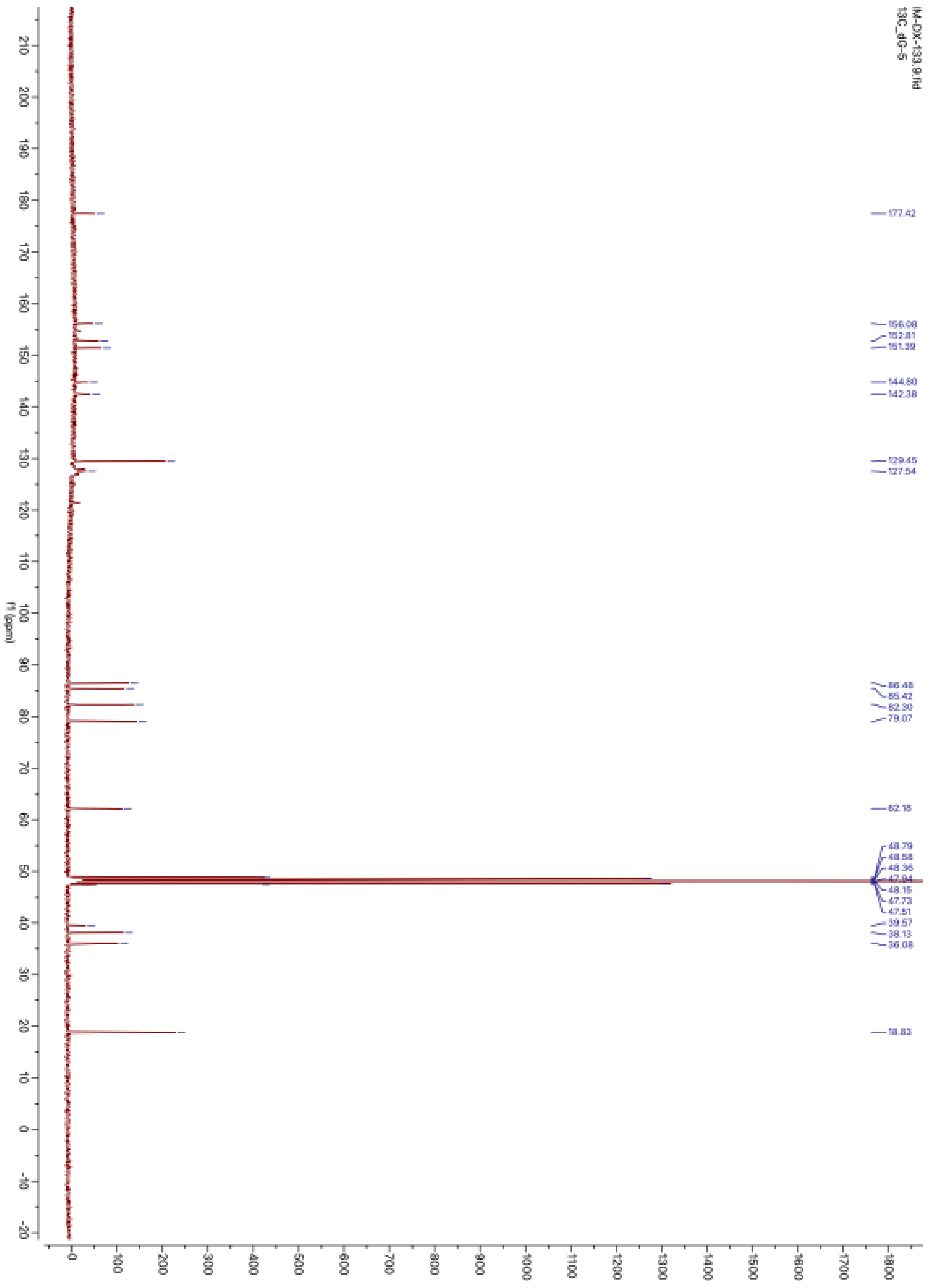

**Supplementary Figure 71.** $^{13}$C NMR of compound **S-8** in $CD_3OD$

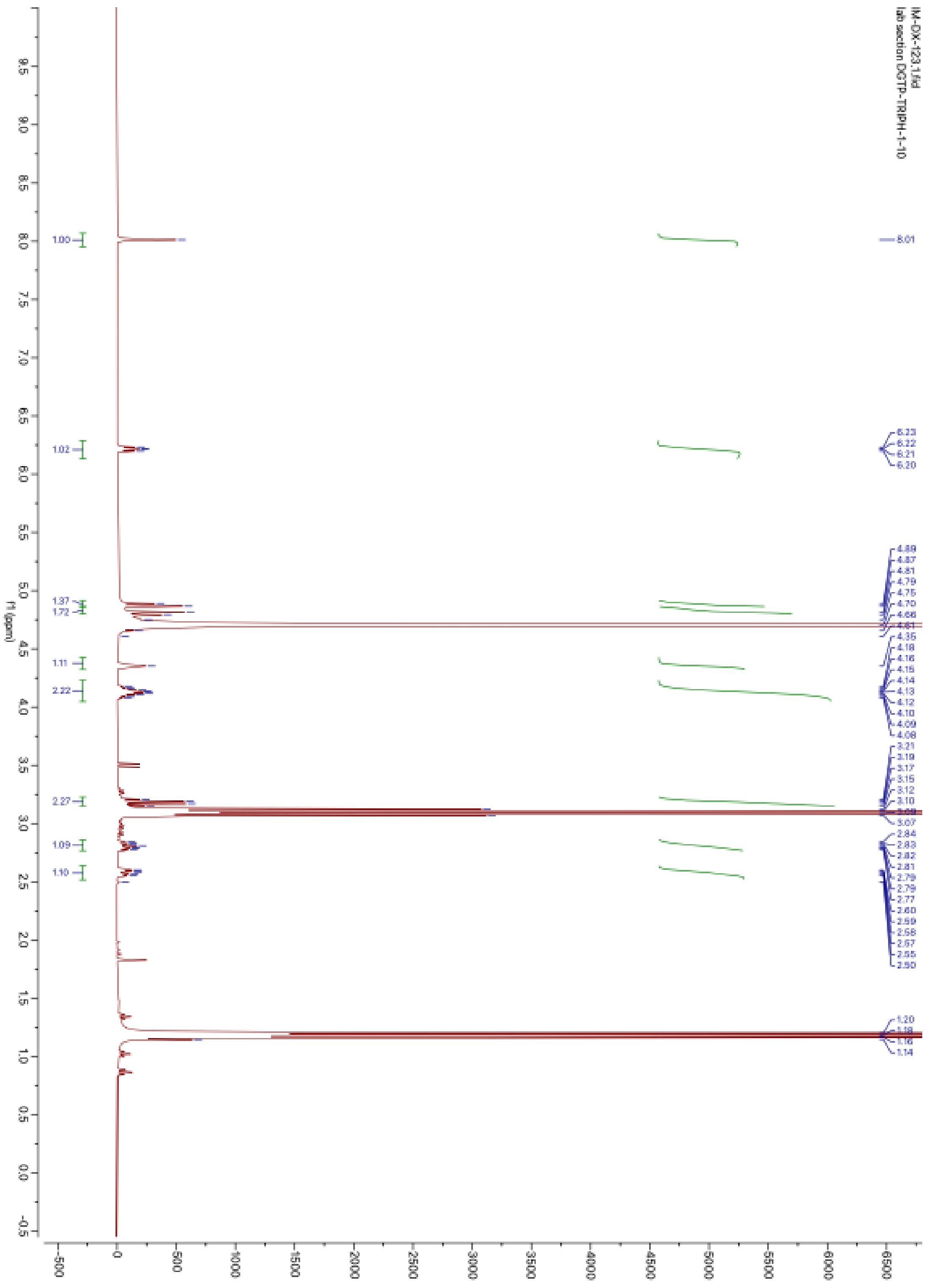

**Supplementary Figure 72.** $^1$H NMR of compound **S-9** in $D_2O$

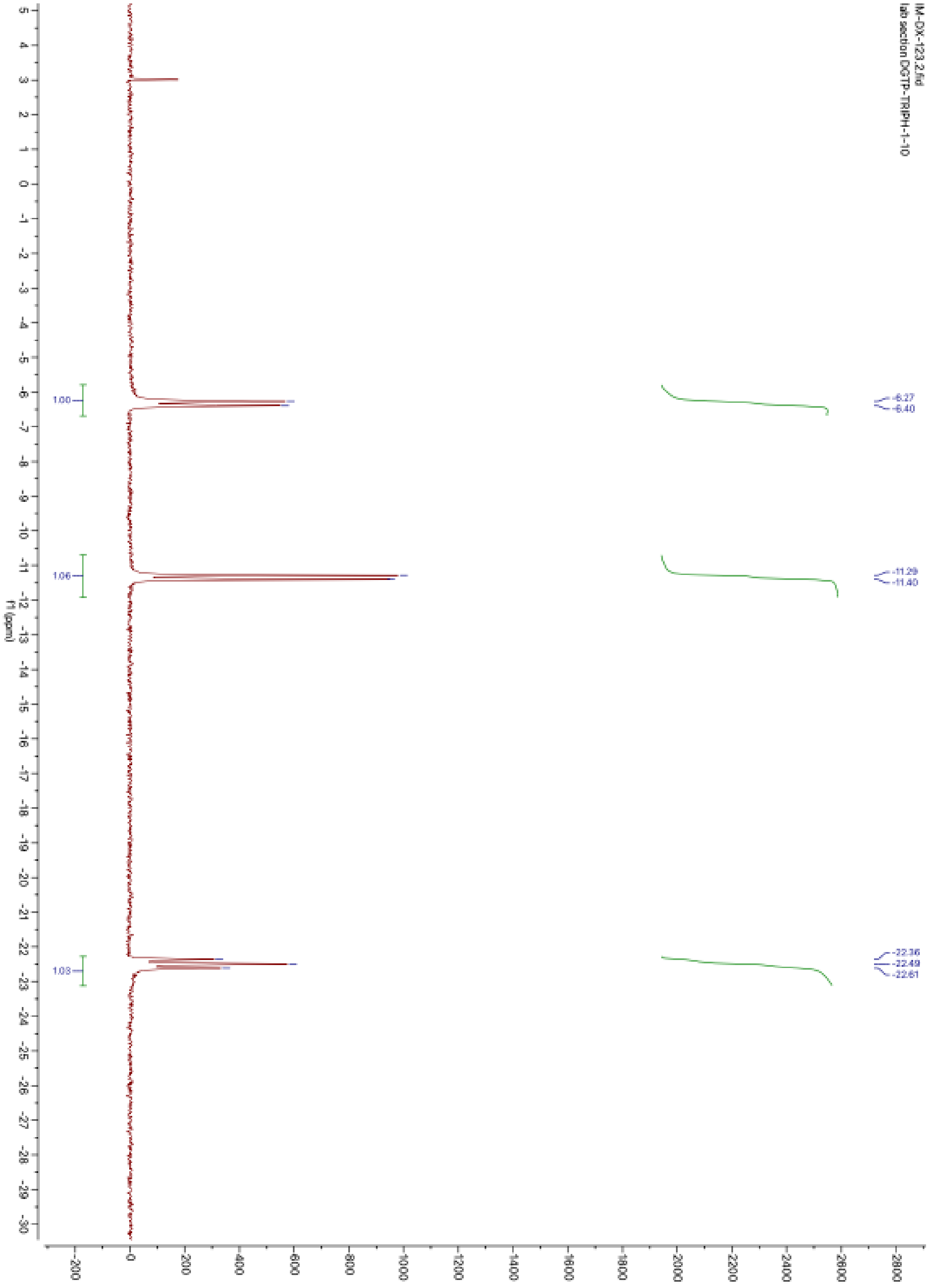

**Supplementary Figure 73.** $^{31}P$ NMR of compound **S-9** in $D_2O$
